# Denisovan leg bones from Taiwan reveal large body size

**DOI:** 10.64898/2026.08.07.743438

**Authors:** Yousuke Kaifu, Chun-Hsiang Chang, Yumeko Tarusawa, Rikai Sawafuji, Shiori Yonemoto, Shigeru Shimamura, Masanaru Takai, Reiko T. Kono, Cheng-Han Sun, Cheng-Hsiu Tsai, Minoru Yoneda, Takumi Tsutaya

## Abstract

Denisovans are an extinct archaic *Homo* group whose lineage diverged from the Neanderthal lineage approximately 550,000 years ago and were widely distributed across eastern Asia until ∼45,000 years ago^1–7^. Their morphological features are known directly from the existing cranio- dental and phalangeal remains^1,2,8–12^. However, the body size and postcranial morphology of the Denisovans remain largely unknown. We here report that hominin femoral and tibial fossils recovered from the Penghu Channel, Taiwan, are Denisovans in their proteomic profiles. Morphologically, these specimens are among the largest leg bones known in Pleistocene *Homo*. They exhibit generally archaic features, but also show some modern human-like morphology, including a strong femoral pilaster. Our findings demonstrate that the Denisovan population at the northern circle had larger body size than earlier *Homo erectus* as well as Late Pleistocene *Homo sapiens* in eastern Asia. This challenges the generally held expectation that Pleistocene *Homo* followed Bergmann’s rule that anticipates latitudinal decline of body size, and suggests that the large Denisovan brain resulted from their large body size at least partly. The strong pilaster developed in the Penghu femur suggests some behavioral similarities between the Denisovans and the Upper Palaeolithic modern humans and/or gene flow from the latter to the former.

## Main Text

Just before the appearance of *Homo sapiens* in the Late Pleistocene, eastern Asia was home to morphologically diverse hominin groups^13,14^ (**Fig. 1**). Whilst diminutive species, *H. floresiensis* and *H. luzonensis*, inhabited isolated islands in Southeast Asia, Java was home to a regional *H. erectus* group, who possessed a body size comparable to that of modern humans. Some researchers have emphasized that the situation on the Asian mainland was also complex from the latter half of the Middle Pleistocene onwards, following the disappearance of *H. erectus* represented by the fossils from Zhoukoudian Lower Cave^15^. For example, while very large cranial specimens are known from the northern China (Harbin, Xujiayao and Xuchang: 45– 34°N), fossils from Hualongdong (30°N) have been argued to exhibit some features foreshadowing modern human morphology (e.g., a pronounced mental trigone of mandible)^16^, and Maba and Narmada (22–23°N) show some similarities to Neanderthals in cranio-facial morphology^13^.

**Fig. 1.**
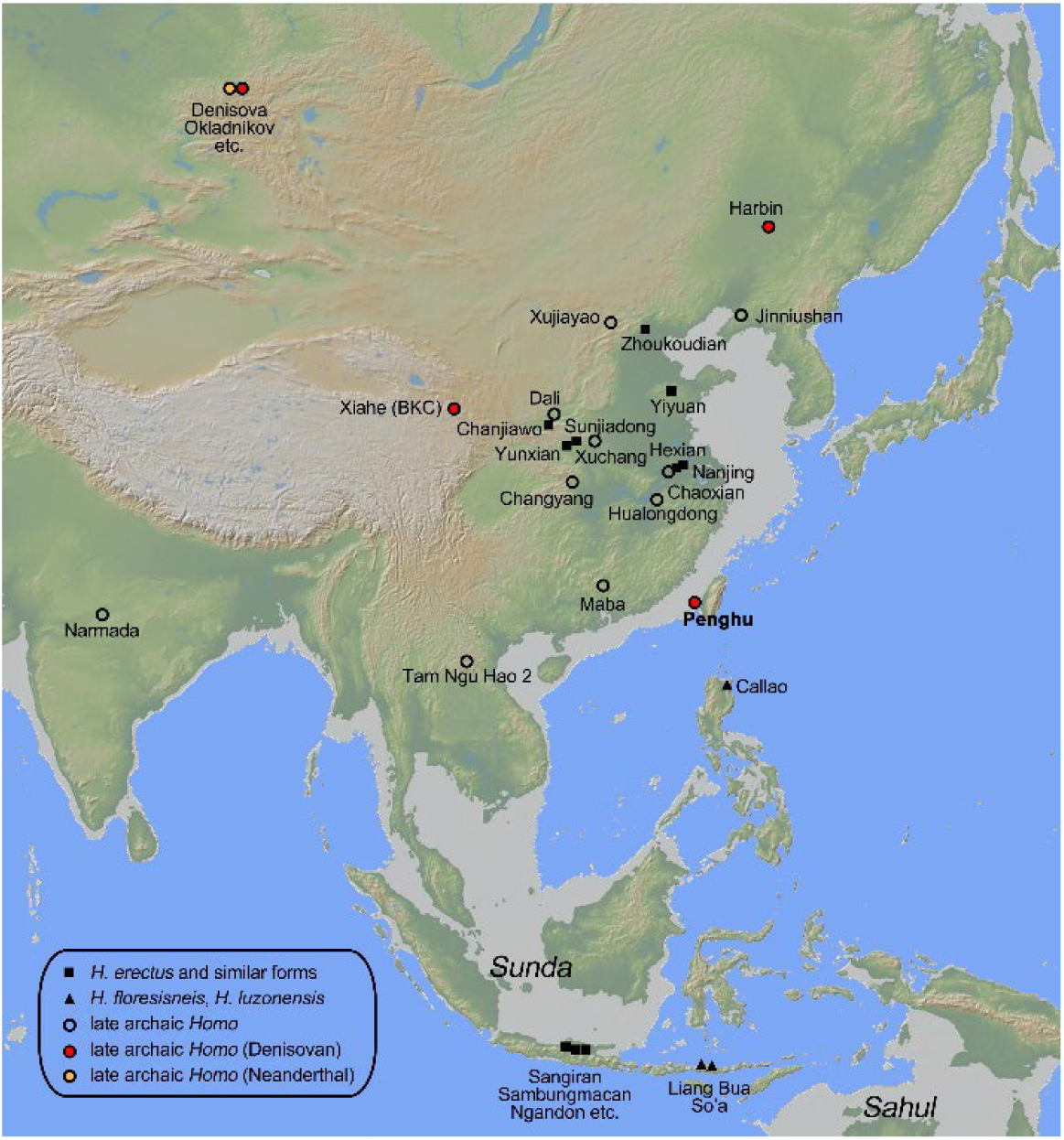
Map of major hominin fossil sites in eastern Asia. The light brown areas in the sea on the map are the continental shelves 0–100 m below sea level. The base map created with GeoMapApp (www.geomapapp.org) / CC BY / CC BY (ref. 53).

A key to understanding this diversity and its evolutionary process is the concept of Denisovans (or ‘Denisova clade’), a sister group of Neanderthal proposed from ancient genomic evidence^1,17^. So far, a well-preserved cranium, two mandibles, dental remains, a phalanx, and other fragmentary fossils from Denisova Cave (southern Siberia), Xiahe (Tibet), Harbin (Northeast China), and Penghu (Taiwan) have been biomolecularly identified as belonging to this clade. Based on the previous morphological studies of these fossils, diagnostic morphological features of the Denisovans now include a large brain size, a robust mandibular body, large teeth, a three-rooted lower second molar, and a reduced/deficient third molar, non-Neanderthal-like (plesiomorphic) phalange, among others^1,2,8–12^.

However, Denisovan’s body architecture such as body size and proportion, skeletal robusticity, as well as their general postcranial morphology remain unknown, except for hypothetical inference based on their DNA^18^, due to the paucity of their postcranial remains. More generally, in the western hemisphere, it is known that large-bodied *Homo* individuals appeared around 1.6 million years ago in Africa (*H. erectus/ergaster*) and this trend continued to the Middle and early Late Pleistocene in Africa and Europe (late archaic *Homo* and early *H. sapiens*)^19–21^, but the situation in eastern Asia is ambiguous.

Penghu fauna is represented by vertebrate fossil remains dredged from the –60 to –120m deep Penghu submarine channel in the Taiwan Strait (**Extended Data Fig. 1**). Located at ∼23°40’N, slightly south of the northern circle, it consists predominantly of terrestrial mammals from cold episodes (i.e., during the low sea-level stands) of late Middle Pleistocene to Late Pleistocene. From this faunal assemblage, we previously reported Penghu 1, a mandible from the male Denisovan, which was obtained from a local antique shop^3,11^. In 2010, we identified fossilized hominin femur (Penghu 2) and tibia (Penghu 3) among a large Penghu fossil collection donated by Mr. Li-Ren Hou to the National Museum of Natural Science, Taiwan (**Fig. 2, Extended Data Fig. 2**). Since 2012, we have conducted a step-by-step research project to analyze morphology and take bone samples for dating, isotopic, genomic and proteomic analyses while minimizing the damage to the fossils (**Supplementary Note 1**). New radiocarbon dates from three Penghu faunal fossils, including Penghu 3, indicate ages ranging from 45,000–43,000 years ago (cal BP)^7^. Therefore, some, if not all, of the Penghu faunal fossils are from Marine Isotope Stage (MIS) 3. In this paper, we report the results of palaeoproteomic and morphological analyses of the two leg bones. There is no record to know when and where in Penghu Channel each of these leg bones was collected, but their different collagen^7^ and overall proteome (**Supplementary Note 3**) preservation imply that Penghu 2 and 3 are from different individuals, or at least from different depositional microenvironments.

**Fig. 2.**
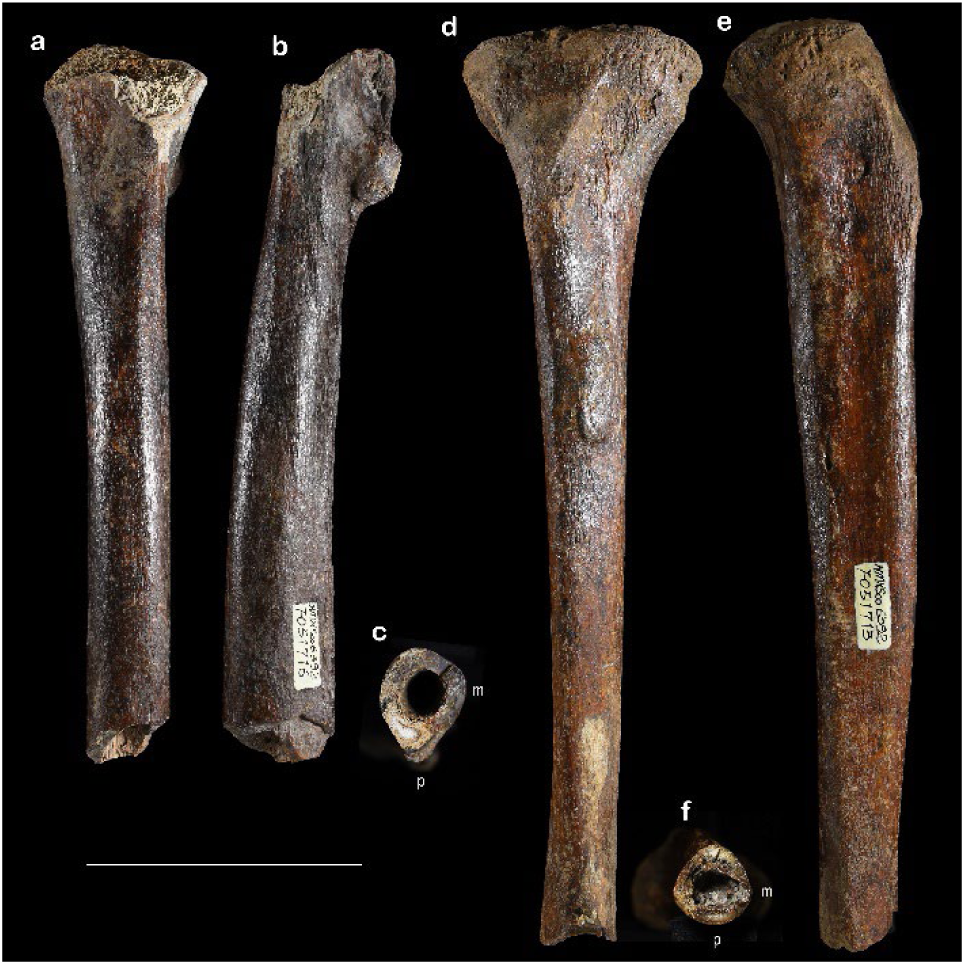
Penghu 2 and Penghu 3 hominin leg bones. Anterior (a), medial (b), and distal (c) views of the Penghu 2 right femoral shaft. Anterior (d), lateral (e) and distal (f) views of the Penghu 3 right tibia. Symbols: m=medial, p=posterior. Scale bar=10cm.

## Proteomic profiles of Penghu 2 and 3

Ancient proteomes of the Penghu 2 femur and Penghu 3 tibia were obtained by liquid chromatography tandem mass spectrometry (LC-MS/MS) to determine their taxonomic attribution. Protein extraction methods and mass spectrometry measurements were optimized based on the results of a previous study^3^ and experiments using a faunal remain from Penghu (**Supplementary Note 2**). Three different methods, including direct application of trypsin or Glu-C and a digestion-free strategy, yielded 21 and 36 endogenous proteins with ≥2 razor+unique peptides from Penghu 2 and Penghu 3, respectively (**Supplementary Note 3**). Totals of 1856 and 3283 amino acid (AA) residues were sequenced from Penghu 2 and Penghu 3, respectively, with validated peptide-spectrum matches (PSMs) (**Supplementary Note 3**).

These values are lower than those sequenced from the bone of Penghu 1 (3546 AA residues^3^) but still represent high-quality ancient proteomes among published hominin bone and dentine proteomes of the Middle to Late Pleistocene^2,4,22,23^. Asparagine (N) and Glutamine (Q) deamidation rates, which increase with the thermal age of the specimen^24^, were higher in the endogenous proteins identified from Penghu 2 and Penghu 3 (i.e., ≥88% in N and ≥67% in Q) than those in contaminant proteins identified from experimental blanks (i.e., ≤25% in N and ≤33% in Q) (**Supplementary Note 4**), suggesting the ancient origin of the retrieved proteomes of Penghu 2 and Penghu 3.

A derived AA variant in type I collagen alpha 2 specific to Denisovans (COL1A2 R996K) was confidently identified both from Penghu 2 and Penghu 3, with the depths of supporting validated peptides of four and ten, respectively (**Extended Data Fig. 3**). No confident peptide with the ancestral R variant was identified at this position (**Supplementary Note 5**). Molecularly identified Denisovans carry the derived K variant at this position^2–4,23,25^, but Neanderthals, and almost all modern humans carry the ancestral R variant in this position (**Extended Data Table 1**). Additional phylogenetically informative variants were sequenced from Penghu 2 and Penghu 3, but the confidence of these variants is ambiguous due to their relatively low depth (**Supplementary Note 5**).

Next, we constructed phylogenetic trees for 15 proteins using maximum-likelihood and Bayesian methods. These trees accurately reflect the phylogenetic relationships among *H. sapiens*, Neanderthals, Denisovans, and great apes. Both Penghu 2 and Penghu 3 clustered with Penghu 1 as well as Denisova 3, a Denisovan individual with a published high-quality nuclear genome sequence from Denisova Cave in Siberia (**Fig. 3**). The same result was obtained with stricter sequencing criteria that require at least two peptides with the spectral support to call the amino acid residue of the position (**Supplementary Note 6**). The results of both the Denisovan-specific variant and the overall phylogeny support the attribution of both specimens to the Denisovan lineage.

**Fig. 3.**
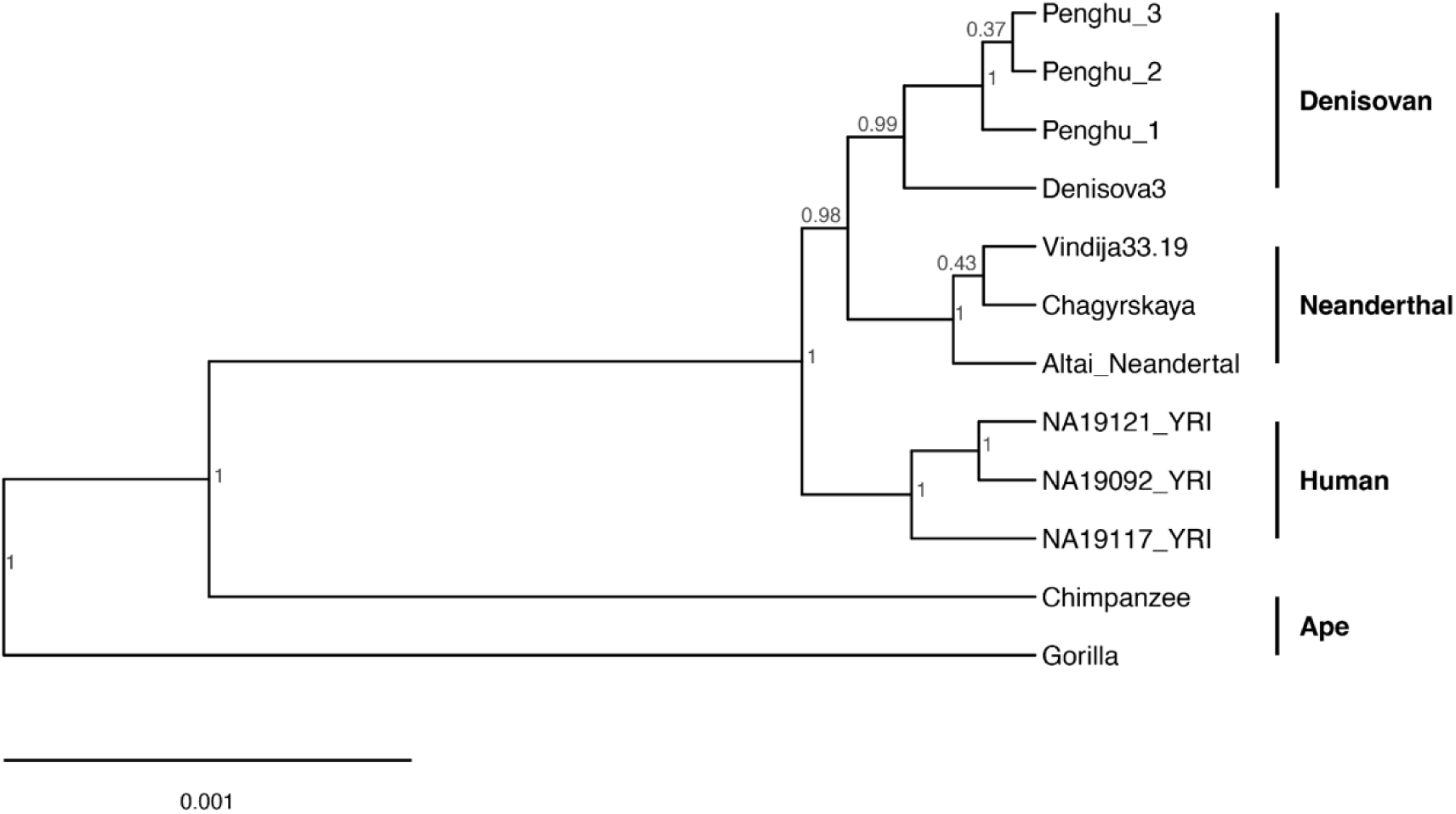
Bayesian phylogenetic tree of hominins and great apes inferred using BEAST 2. Numbers at nodes indicate posterior probabilities (0–1). The orangutan sample was excluded from the displayed tree.

## Morphology of Penghu 2 (femur)

Penghu 2 is a well-preserved, proximal half of the right femur that lacks much of the neck and greater trochanter. Given its unusual combination of features (a large, archaic hominin femur with a modern human-like strong pilaster, as detailed later), as illustrated in **Extended Data Fig. 4**, we estimated the original length of Penghu 2 by referring to selected large modern and archaic hominin femora that are morphologically and dimensionally similar to Penghu 2. The reconstructed three models yielded femoral maximum lengths (FMLs) and biomechanical lengths (FBLs) of 493‒489 and 466‒456 mm, respectively. We use their means (491 and 461 mm) in the following comparisons. These estimates are consistent with the general shaft morphology of modern and archaic hominin femora (**Extended Data Fig. 5**).

When the length and midshaft thickness (TA) are compared together, Penghu 2 is one of the largest femora among the global Pleistocene *Homo* sample (**Fig. 4a, Extended Data Fig. 5a,b**). Likewise, the area proximal to the large lesser trochanter is dimensionally comparable to or slightly greater than long modern human femora in our sample (max. length ∼500 mm: **Extended Data Fig. 4b**). Relative cortical area of Penghu 2 is moderate among the archaic hominins, but is at the upper end of variation exhibited by the Holocene modern human hunter-gatherers (**Extended Data Fig. 5c,d**).

**Fig. 4.**
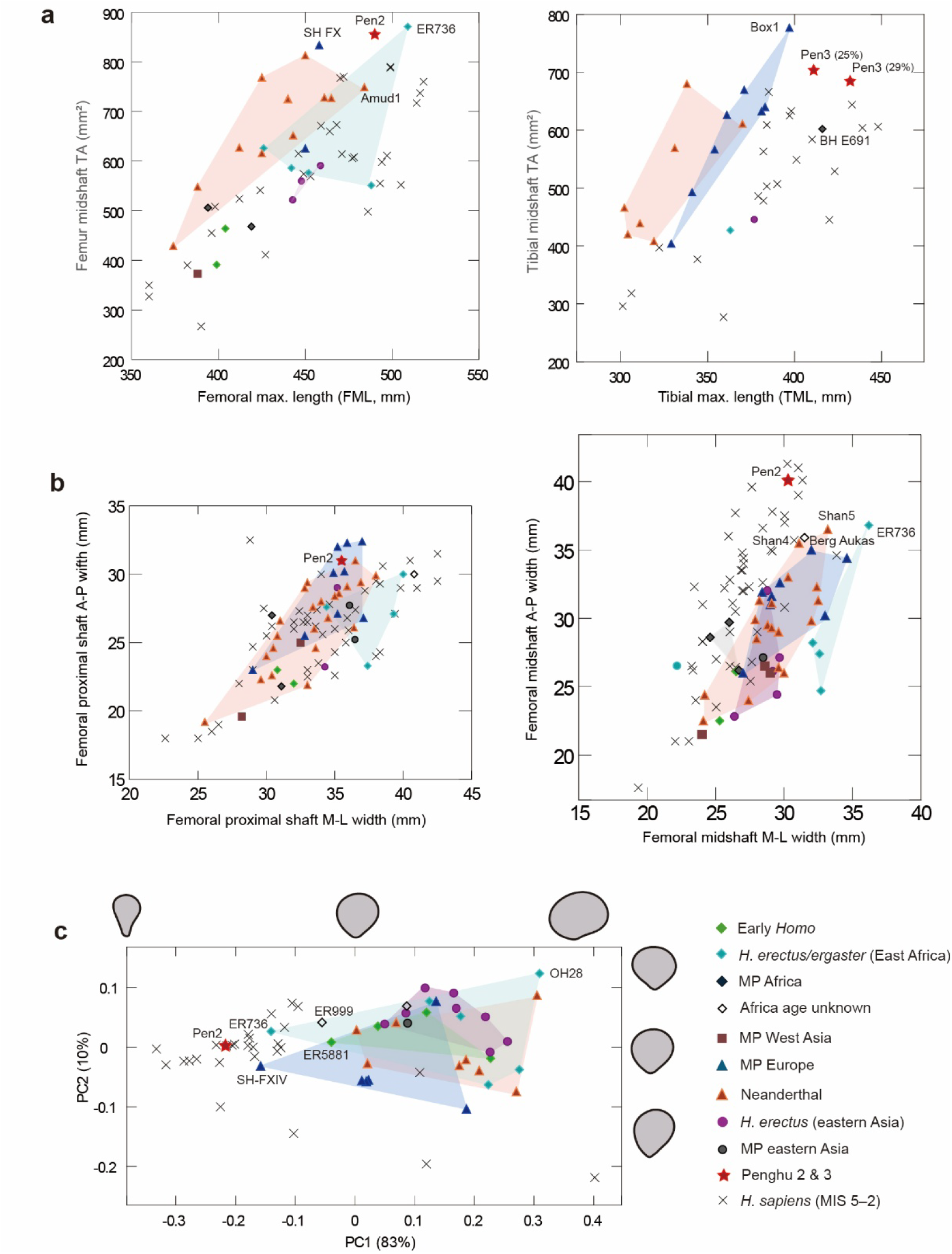
Metric comparisons of Penghu 2 and 3 with modern and archaic *Homo* groups. a, Scatter plots of femoral/tibial maximum length and midshaft thickness (total cross-sectional area: TA) for Penghu 2 and 3. The values based on the two length estimates are shown for Penghu 3. **b**, Anteroposterior (A-P) and mediolateral (M-L) shaft dimensions of femora. **c**, Plots of PC (principal component) scores derived from the normalized Elliptic Fourier Analyses (EFAs), based on midshaft cross-sectional contours. The proportion of variance explained by each PC are in parentheses. Shape differences along PC axes are shown on right femora, with anterior directed up and lateral right, two standard deviations from the origin. Convex hulls are depicted for some archaic *Homo* groups with larger sample sizes.

The preserved basal part of the neck of Penghu 2 is oriented superiorly rather than transversely. Viewed anteriorly, the base of the greater trochanter flares laterally, as commonly seen in *H. erectus* and later *Homo* species^26^. The posterior face of the proximal shaft is marked by moderately developed gluteal tuberosity, hypotrochanteric fossa, pectineal line and other muscle markings. It is known that modern humans typically show gradual mediolateral widening of the femoral shaft, from a level proximal to the midshaft toward the distal metaphysis. The midshaft of Penghu 2 does not show this trend. As commonly seen in archaic *Homo* including Asian *H. erectus*^27,28^, its mediolateral width remains stable beyond its midshaft down to the broken distal end (**Extended Data Fig. 5h**). Among the premodern *Homo* groups, an anteroposteriorly thickened (non-platymeric) proximal shaft characterizes Neanderthals and in particular Middle Pleistocene European *Homo*, as compared to African Early-Middle Pleistocene *Homo* (**Fig. 4b**). Penghu 2 is similar to the Middle Pleistocene European *Homo* in this respect. Such a trend is not evident in the available small sample of *H. erectus* and Middle Pleistocene *Homo* from eastern Asia.

Despite the above generally archaic characters, Penghu 2 has, unlike any other archaic hominin femora^29–32^, a distinct pilaster on its posterior surface. It is thick, angular, and projects strongly posteriorly, as seen in many individuals of Upper Palaeolithic *H. sapiens* (**Figs. 4c** and **5**).

**Fig. 5.**
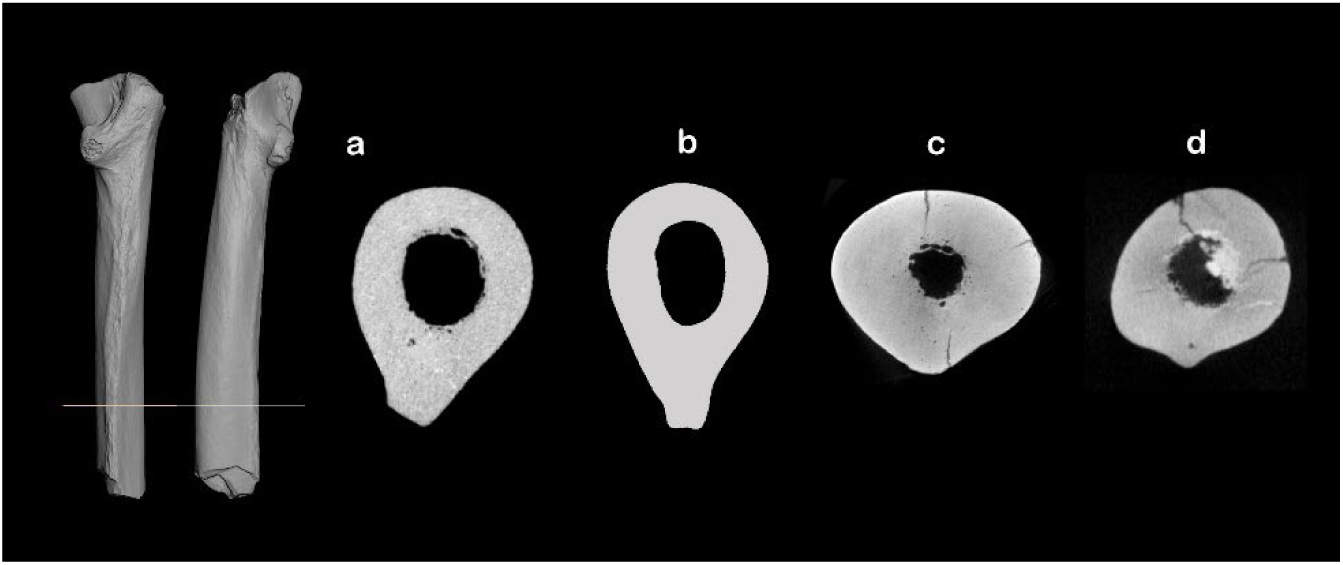
Femoral midshaft cross-sections of archaic and modern humans showing varying development of pilaster. Mid-shaft sections of Penghu 2 (based on CT) (**a**), MIS 3 *H. sapiens* from Zhoukoudian Upper Cave (Femur II, based on drawing) (**b**), late archaic *Homo* from Arago, France (A141, based on CT) (**c**), and Neanderthal from Ehringsdorf (based on CT) (**d**). Anterior directed up and lateral right. Not to scale. Sectioned level is indicated for Penghu 2. Note the strong development of femoral pilaster, which can be defined as “a distinct crest of bone along the posterior midshaft, producing a teardrop-shaped cross-section and straight or concave surfaces posteromedially and posterolaterally”^29^, in Upper Cave Femur II and Penghu 2. **b**: modified from ref. 27. **c** and **d**: courtesy of Tony Chevalier (CT scan of d obtained from NEOPOS).

## Morphology of Penghu 3 (tibia)

Penghu 3 is a well-preserved right tibia that lacks the distal end. We removed the nodules that had covered its external surfaces (**Extended Data Fig. 1**). On the medial surface, approximately 50 mm distal to the tibial tuberosity is a bulging healed trauma of 40 mm long and 16 mm wide. We virtually removed it for our digital measurements.

Because the distal break occurs proximally to the fibular notch and the cortical bone is still thick at this level (**Fig. 2**, **Extended Data Fig. 6h**), the original length of Penghu 3 must have been substantially longer than its preserved maximum length of 340 mm. We estimated its original length following the data presented in **Extended Data Fig. 6**. When aligned in the standard orientation^33^, the shaft of Penghu 3 becomes thinnest in terms of total cross-sectional area (TA) at a distance of 290 mm from the proximal landmark of tibial biomechanical length (TBL) (**a**). In our sample of modern humans (N=66) and a Neanderthal (N=1), such thinning occurs at 25−38% (mean=32%) level of their TBLs from the distal landmark (**b**), with a weak negative correlation with the TBL (*r*=−0.280, *p*=0.024) (**d**). By applying the above minimum (25%) and the ratio of the longest tibia in our sample (29%, TBL=392 mm), the TBL of Penghu 3 is calculated as >386.7 or 408.5 mm, respectively. The 29% model seems to be better because the 25% model does not conform well to the typical pattern of shaft-thickness transition: The slopes between the 20−25% TBL levels in our comparative specimens tend to be steeper in thicker tibiae (**b**, **e**), and the thick Penghu 3 would be an outlier in this trend if the 25% model is taken. Finally, by adding 24 mm (the average difference between MTL and BML for long modern human tibiae: BML=386−425 mm, N=73)^34^ to the estimated TBLs, the maximum tibial length (MTL) of Penghu 3 was calculated to be >411 mm (25% model) or probably ∼432 mm (29% model).

In terms of length and midshaft thickness, Penghu 3 is one of the largest known Pleistocene *Homo* tibiae (**Fig. 4a**). The great length of Penghu 3 contrasts particularly with generally short tibiae of Neanderthals (**Extended Data Table 2**). The proximal articular end of Penghu 3 is dimensionally comparable to that of the large Middle Pleistocene tibia from Zambia (Broken Hill E691)^35^ (their mediolateral widths are 88.4 mm for Penghu 3 and 88.1 mm for E691). The relatively straight shaft (both in anterior and lateral views) is slightly thinner compared to Boxgrove 1^36,37^, but thicker than most other archaic *Homo* specimens including Neanderthals, Atapuerca SH and Broken Hill (**Extended Data Fig. 6c**). The relative cortical area (CA/TA in **Extended Data Fig. 6f,g**) is comparatively high among the archaic *Homo*, but Penghu 3 does not show a tendency of elevated CA/TA values in its proximal shaft, which is seen in all three African tibiae (KNM-ER 803, 1481, and Broken Hill E691) and one of the European specimens (A120)^31^. Penghu 3 is similar in the latter respect to other Eurasian archaic tibiae (as well as the Holocene modern human hunter-gatherers of the Japanese Archipelago).

Penghu 3 exhibits a strong tibial posterior pilaster, a shared feature of European Middle Pleistocene *Homo* and Neanderthals, as well as some Upper Palaeolithic modern humans^38–40^. In cross-sectional morphology of the tibial shaft (**Extended Data Fig. 7**), Upper Palaeolithic modern humans tend to show, with some overlap with archaic *Homo*, mediolateral flattening and sharp anterior, interosseous and posterior crests/margins, as well as a marked concavity of the lateral surface^29,38^. Penghu 3 shows a slight tendency toward this modern human condition (**Extended Data Fig. 7b**), but also exhibits archaic characters such as a thickened anterior margin and a blunt interosseous crest.

## Implications of large body size

Based on the molecularly identified Denisovan fossils from Denisova Cave, Xiahe, Penghu and Harbin, their morphological features currently include a large brain size, a robust mandible, large teeth, a three-rooted lower second molar, and a reduced or absent third molar, among others^2,11,12^. The two Penghu leg bones reported here add a large body size to this potential list. Although we currently do not have biomolecular data to suggest their sexes, the very large dimensions of Penghu 2 and 3 suggest that these are from male individuals.

The estimated body sizes of Penghu 2 (stature: ∼180 cm, weight: ∼83 kg) and Penghu 3 (stature: ∼190 cm, weight: ∼91 kg), which are comparable to the largest known Pleistocene *Homo* from Africa and Europe, stand out among the reported estimates of archaic hominins from mainland eastern Asia (**Extended Data Table 3**). The estimated statures and body masses for ∼0.75 million years ago *H. erectus* from Zhoukoudian, northern China are 20–30 cm shorter and 20–30 kg less, respectively, compared to Penghu 2 and 3. As one of the representative sites of late archaic *Homo* fossils in mainland eastern Asia, ∼0.3 million years ago Hualongdong site in Anhui, China, yielded one femoral midshaft with estimated body mass comparable to those of Zhoukoudian *H. erectus*^21^; Two other proximal femoral shafts from this site are likewise not large^30^ (**Fig. 4**). In contrast, Jinniushan 1, a partial female skeleton from northern China weighed ∼74.2 kg and was ∼169 cm tall^21,41^. If this individual belonged to the Denisovan clade as anticipated by some researchers^13,42^ (but see ref. 43), it implies that large-bodied Denisovan individuals inhabited both northern and southern regions of Asian mainland, and large brain sizes seen in some Denisovan individuals (Harbin 1, and possibly other specimens such as Xujiayao 6 and Xuchang 1) was at least partly a consequence of their large body sizes, as has been suggested for Neanderthals^19^.

A widely held expectation is that Bergmann’s rule, which anticipates latitudinal decline of body size, applies to Pleistocene *Homo* in general and to both Pleistocene and Holocene hominin populations in eastern Asia^19,44–46^. However, the large body size of the Taiwanese Denisovans emphasizes the potential role of other factors. The uniqueness of the Penghu fossils, an assemblage from cold periods including MIS 3 (based on the radiocarbon dates of Penghu 3 and two other faunal remains^7^), is further highlighted when compared to the local Late Pleistocene modern humans. MIS 5‒3 *H. sapiens* from the western hemisphere are characterized by large body size but such a trend is not known among the available small sample from eastern Asia^21,45^ (**Extended Data Table 3**). This means that the glacial climate does not solely explain the extremely large body size of the Penghu Denisovans. Given the recent evidence that Denisovan population in Tibetan Plateau exploited a wide range of animal taxa including Caprinae, megaherbivores and carnivores^23^, and isotopic evidence that the Penghu 3 individual heavily relied on meat^7^, one possible hypothesis is that the large body size was essential for their hunting strategy relying on a tool kit different from those of the local, MIS 3 *H. sapiens*, who achieved explosive geographic expansion beyond the distribution of the Denisovans, and invented sophisticated tools such as seagoing crafts and fishhooks^47–49^.

## Regional features of late archaic *Homo*

The lineages of Denisovan and Neanderthal split from their common ancestral population 585,000‒504,000 years ago^6^, and afterward they developed contrasting cranial, mandibular, and dental features as mentioned above. The present study adds some postcranial features to this list. Penghu 2 shares with European Middle Pleistocene archaic *Homo* and Neanderthals an anteroposteriorly wide proximal femoral shaft and reduced relative cortical area in proximal tibial shaft. However, Penghu 3 does not show marked shortening of tibial length, a derived feature of Neanderthals^40^.

Apart from the above regional features, the Penghu leg bones show generally archaic appearance, such as a large size, general robusticity with relatively thick cortical bones, a distal position of the femoral shaft minimum breadth, as well as non-sharp cresting and non-deep concavity of the tibial surfaces. However, the Penghu 3 tibia exhibits a slightly modern human-like midshaft cross-sectional contour (**Extended Data Fig. 7b**), and remarkably, the Penghu 2 femur has a well-developed pilaster (**Figs. 2, 4 and 5**), a hallmark of modern human hunter-gatherers.

## Implications of femoral pilaster in Penghu 2

Some archaic hominin femora have sharply raised *linea aspera* (e.g., KNM-ER 736, Atapuerca SH F-XIV: **Fig. 4**), but these are distinct from the conditions seen in *H. sapiens* and Penghu 2, which are characterized by thick, angular, strong posterior projection. Why Penghu 2 developed a femoral pilaster, a distinct modern human feature, among its generally archaic morphology is an intriguing question. Because strong femoral pilaster, or higher femoral midshaft A-P/M-L bending rigidity, is associated with greater terrestrial mobility of modern human hunter- gatherers^50,51^, one explanation would be that the Taiwanese Denisovans experienced such a mobile lifestyle similar to early modern humans. However, this hypothesis may not be compatible with the above suggested different hunting strategies with different tool kits. Another factor that may promote strong femoral pilaster development is a narrow body shape^45^, but we currently do not have fossil materials to directly assess Denisovan pelvic breadth, and this hypothesis contradicts with an expectation from the reconstructed Denisovan pelvic size based on DNA methylation patterns^18^. Lastly, the other non-mutually exclusive explanation is that the Taiwanese Denisovans acquired femoral pilaster (and possibly the tibial cross-sectional morphology) through gene flow from early modern humans. Although genetic basis of femoral pilaster remains unknown and this hypothesis must be tested ultimately by genomic data from Denisovan fossils, it does not contradict with the genomic evidence of Denisovan introgression into modern humans in southern Asia^6,52^, and chronological overlap between Penghu 3 and the earliest modern humans in this region^7^.

## Methods

Transfers of Penghu 2 and 3 to Tokyo for CT scan and/or bone sampling in 2012, 2019 and 2024 were conducted with permissions issued from the National Museum of Natural Science, Taiwan.

### Proteomic analysis

The full details of the proteomic analysis are described in **Supplementary Note 7**.

### Protein extraction and measurement

Ancient proteins were extracted from a total of 37.9 mg and 22.4 mg of bone powder samples drilled from the inner part of the Penghu 2 femur and the Penghu 3 tibia, respectively. Based on the newly conducted evaluations of protein extraction and measurement efficiency using faunal material from Penghu (**Supplementary Note 2**), as well as the result of previous study^3^, the bone powder was divided into four aliquots and treated with the direct application of two different proteases (i.e., trypsin and Glu-C)^54^, as well as a digestion-free method^55^ to obtain the maximum coverage of amino acid residues^56^. Following the best practices of palaeoproteomics^57^, sampling and extraction procedures were carried out in a clean laboratory dedicated to ancient biomolecules. The extracted proteins/peptides were purified with C18 StageTips and sequenced with LC-MS/MS (Ultimate 3000 RSLCnano and Fusion Tribrid Orbitrap mass spectrometer).

### Sequence reconstruction

The resulting .raw files were searched with pFind^58^ and MaxQuant^59^ to detect variants and identify proteins. The protein sequence databases containing proteins with archaic variations were generated by PaleoProPhyler and used for database searching^60^.

Identified peptide-spectrum matches were computationally validated based on the R script *MS2Ladder*^61^, and only residues that were supported by tandem mass spectra were used to construct ancient protein sequences. Modern contaminant proteins were removed from the datasets. Deamidation rates were calculated based on the R script *rDeamidation* ported from the previously reported Python script^62^. Proteomic raw data and results are uploaded to the PRIDE repository with the identifier PXD080741^63^. The key to raw file attribution is found in sdrf- vertebrates.txt. Sequence reconstruction was performed in the R software environment (R Core Team, 2023) with R scripts^64^ that are accessible in the Zenodo repository.

### Phylogenetic analysis

Phylogenetic trees were constructed based on the sequences of endogenous proteins (1627 residues in Penghu 2 and 2892 residues in Penghu 3) that have ≥7 razor+unique peptides in either Penghu 2 or Penghu 3. Corresponding sequences from Penghu 1 were included for comparison. Protein sequences of modern humans from the Yoruba population (n=3), archaic humans (n=4), and great apes (chimpanzee, gorilla, orangutan) were obtained from the Hominid Palaeoproteomic Reference Dataset^60,65^, UniProt, and GenBank. Maximum-likelihood and Bayesian phylogenetic analyses were performed using IQ-TREE 2^66^ and BEAST 2^67^, respectively. The scripts^68^ used for phylogenetic analysis are accessible in Zenodo repository.

### Morphological analysis

#### Comparative sample

Penghu 2 and 3 were compared with various taxa/groups of Early to Late Pleistocene *Homo* (early *Homo*, *H. erectus/ergaster*, late archaic *Homo*, and early *H. sapiens*) except for small-bodied species from the Middle and Late Pleistocene contexts (i.e., *H. floresiensis*, *H. luzonensis*, and *H. naledi*) (**Supplementary Table 8**). To estimate the original bone length of Penghu 2 and 3, we referred to CT-derived metric data of mixed sex, adult samples of the Holocene hunter-gatherers in the Jomon period of Japan (N=21 for femur and N=48 for tibia, housed at the University Museum, The University of Tokyo) (see **Source Data** for the details). Additional comparative materials to estimate the original lengths of Penghu 2 and 3 were a few large femora from the Yayoi period of Japan (Doigahama site, Yamaguchi Prefecture) and casts of Neanderthal femur (Feldhofer 1) and tibia (Spy 2). We did not discriminate sex in our comparative analyses because the sex of each fossil specimen is generally unknown.

#### CT scan and 3D mesh data

Penghu 2 and 3 were μCT-scanned using TX225-ACTIS (Tesco Co.) at the University Museum, University of Tokyo, at 150 kV and 200 μA, at final resolutions of 95.40225 and 133.5449 μm/voxel, respectively. CT scans of the Holocene Jomon femora and tibiae were performed with the resolution of 104.248‒156.3721 μm/voxel. Threshold-based and manual segmentation of the CT data as well as production of 3D mesh models were conducted using Avizo 3D (Thermo Fisher Scientific). The traumatic bony bulge of Penghu 3 was digitally removed during this process to recover the original, unaffected bone surface, before measuring cross-sectional geometry. Further preparation of 3D mesh and extraction of cross-sectional images were performed using the MeshLab and Geomagic Wrap (3D Systems) software.

#### Metric analysis

Linear and angle measurements of the original specimens were taken directly using a digital caliper (Mitsutoyo, Inc.) and a calibrated osteometric board, or by meshLab based on 3D surface mesh created from CT scans. Serial cross-sectional metrics and properties along the femoral and tibial shaft (total area, relative cortical area, etc.) were obtained using the morphomap R code package^69^ with some modifications (**Software 1**), following the standard procedure defined by Ruff^33^ and based on 3D mesh data. Alignment of complete femora and tibiae to the standard plane was done using the R code offered in morphomap^69^ and **Software 2**, respectively. Incomplete Penghu 2 and Penghu 3 as well as a few Jomon femora were aligned by fitting them to the aligned complete leg bones with similar size and general morphology, using MeshLab.

Midshaft (50% level of biomechanical length) cross-sectional contours of femora and tibiae were analyzed by normalized (i.e., size-standardized) Elliptic Fourier Analysis (EFA), using the images from the right side or horizontally flipped images of the left bones. To compute normalized elliptical Fourier descriptors (EFDs), each contour was aligned so that the linear aspera faces posteriorly (femur) or along the long axis of its first ellipse (tibia).

Calculation of EFDs and PCAs of the normalized EFDs were conducted using the software SHAPE v. 1.3^70^.

## Data and code availability

All data generated or analyzed during this study are included in this published article (and its supplementary information files) or as a Source Data file. The **Source Data** file includes raw data used for **Figs 2 and 3, and Extended Data Figs. 3, 5, 6 and 7.** The Penghu hominin fossils are housed at the National Museum of Natural Science, Taiwan. The 3D data of Penghu 2 and 3 may be shared on request to Chun-Hsiang Chang. Source data are provided in this paper.

Proteomic raw data and analytical outputs were uploaded to the PRIDE repository. R codes used in this study are available in published scripts or as supplementary information files associated with this paper.

## Acknowledgments

C-H.C. sincerely thanks Mr. Li-Ren Hou for his generous donation of the Penghu faunal fossils to the National Museum of Natural Science, Taiwan. We also thank Takashi Gakuhari, Hiroki Oota and Kae Koganebuchi for bone sampling for chemical analyses, Ken Takai for coordinating the mass spectrometry analyses, and Sora Oniki and Asuka Shirakami for CT data processing, and Katherine Hampson for editorial assistance. This work was supported by JSPS KAKENHI Grant Numbers 22H00421, 23K17404, 23H00009, 19H05350, 20KK0166 and 21H00337, NSTC 112-2116-M-178-001-, NTU FD107028, and JST FOREST program Grant Number JPMJFR233D.

## Author contributions

C-H.C. and Y.K. conceived the study. Y.K., C-H.C. and S.Y. collected morphological data. Y.K. analyzed the morphological data. T.T., Y.T., R.S. and S.S. collected and analyzed proteomic data. Y.K., Y.T., R.S, and T.T. wrote the codes for computer-based data analysis. Y.K. and T.T. wrote the manuscript with critical input from the remaining authors.

## Competing interests

The authors declare no competing interests.

---

**Extended Data Fig. 1.**
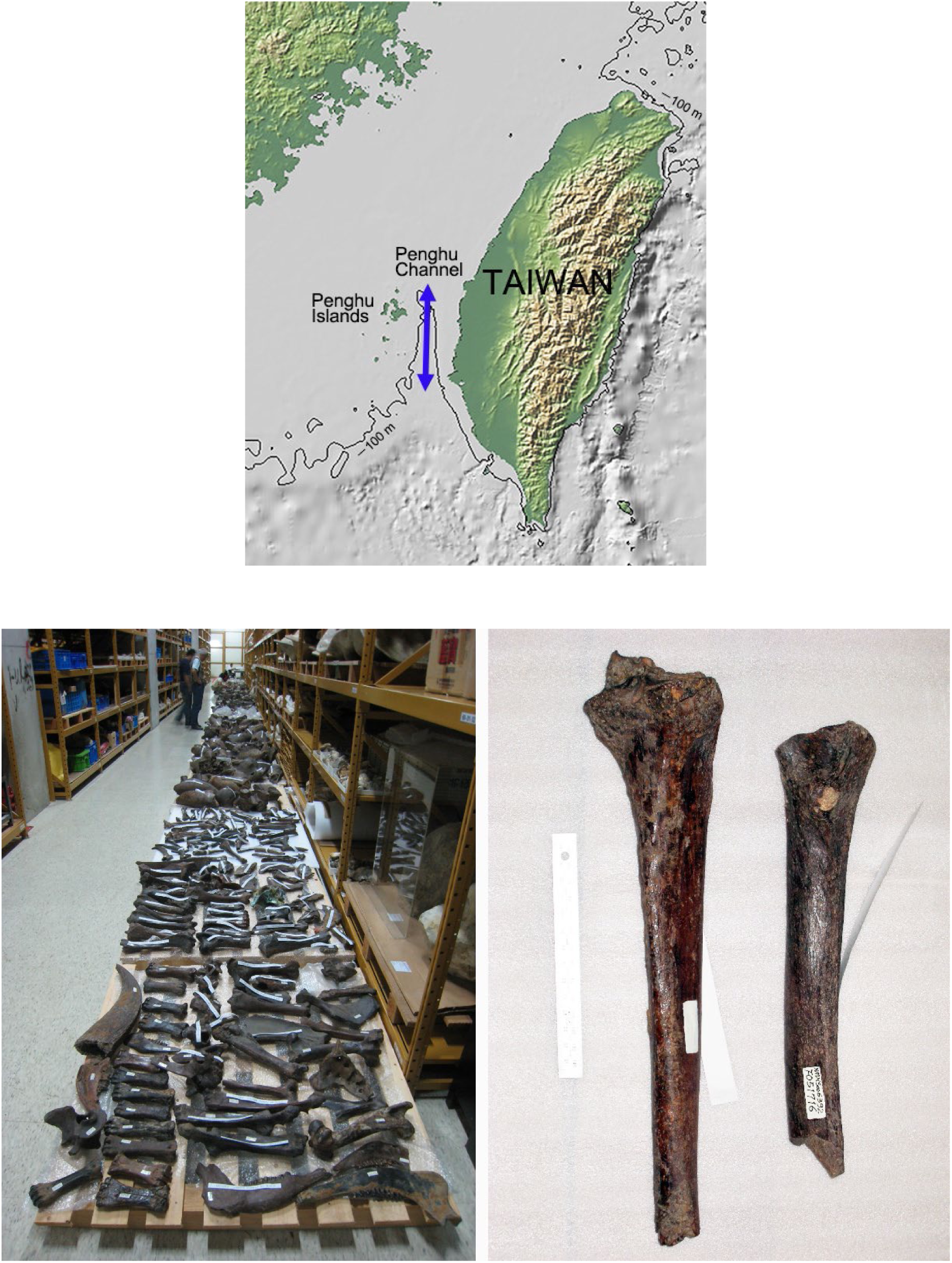
Map of the Penghu Channel and the discovery of Penghu 2 and 3. Upper: Submarine topography around the Penghu Channel. The base map created with GeoMapApp (www.geomapapp.org) / CC BY / CC BY (ref. 53). Lower left: A part of Mr. Li- Ren Hou’s Penghu fossil collection, which was donated to the National Museum of Natural Science, Taiwan, in 2011 (this photograph was taken in 2010 during preparation work). Lower right: Penghu 2 (right) and 3 (left) at the time of our identification in 2010. Proximal articular surface and some other parts of Penghu 3 were covered by nodules. We cut their distal ends in 2012 for U-series dating^71^.

**Extended Data Fig. 2.**
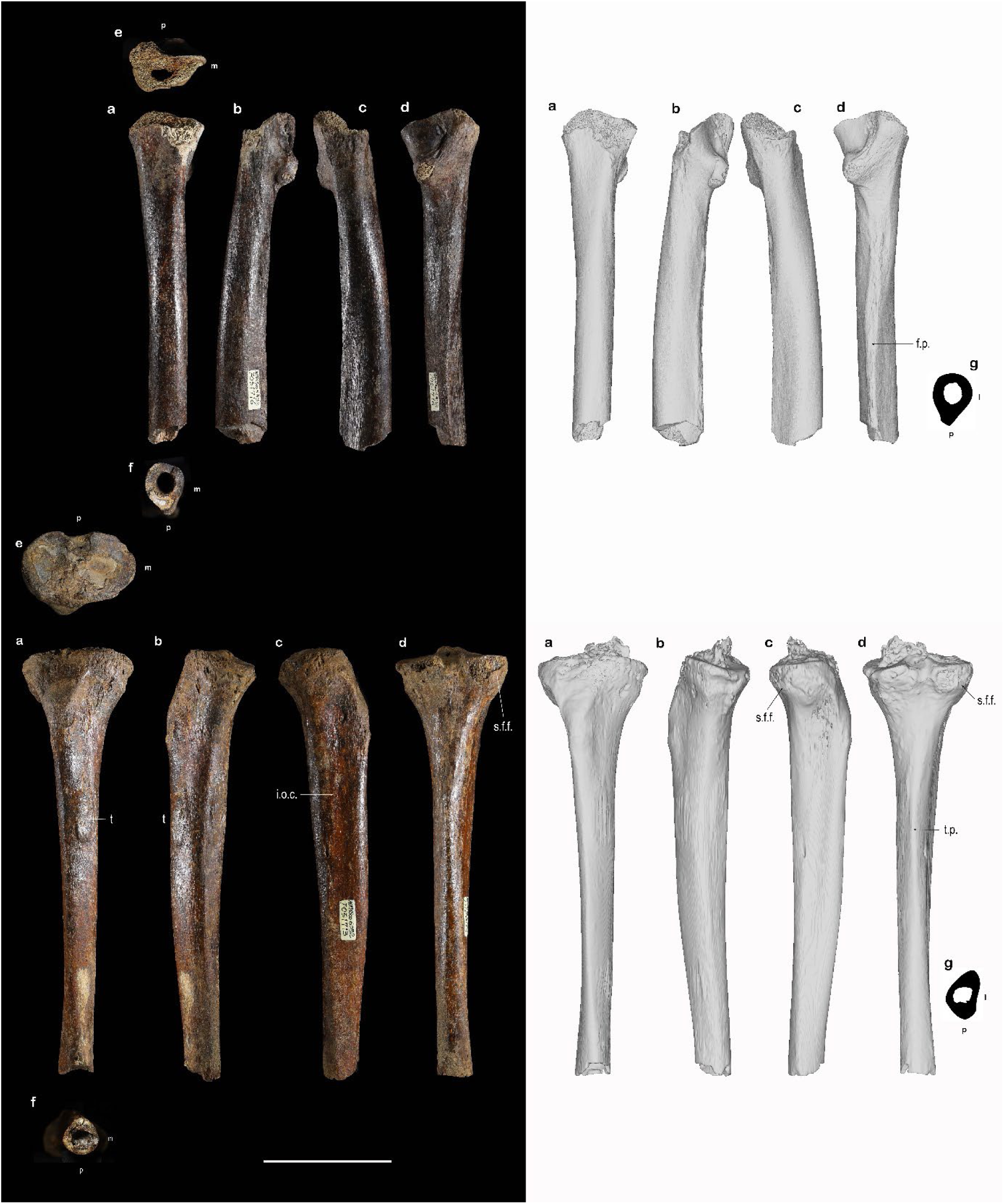
Pictures (left) and surface rendered images (right) of the Penghu 2 femur (upper) and Penghu 3 tibia (lower). Anterior (a), medial (b), lateral (c), posterior (d), superior (e) and inferior (f) views, as well as superior views of mid-shaft cross-sections (g). The pictures of Penghu 3 are after the physical cleaning in 2025. The surface rendered images of Penghu 3 are before that cleaning and after digital removal of some (but not all) nodules and the traumatic swelling on the medial face. Symbols: l=lateral, m=medial, p=posterior, f.p.=femoral pilaster, t=traumatic swelling, i.o.c.=interosseus crest, s.f.f.=superior fibular facet, t.p.=tibial pilaster. Scale bar=10 cm.

**Extended Data Fig. 3.**
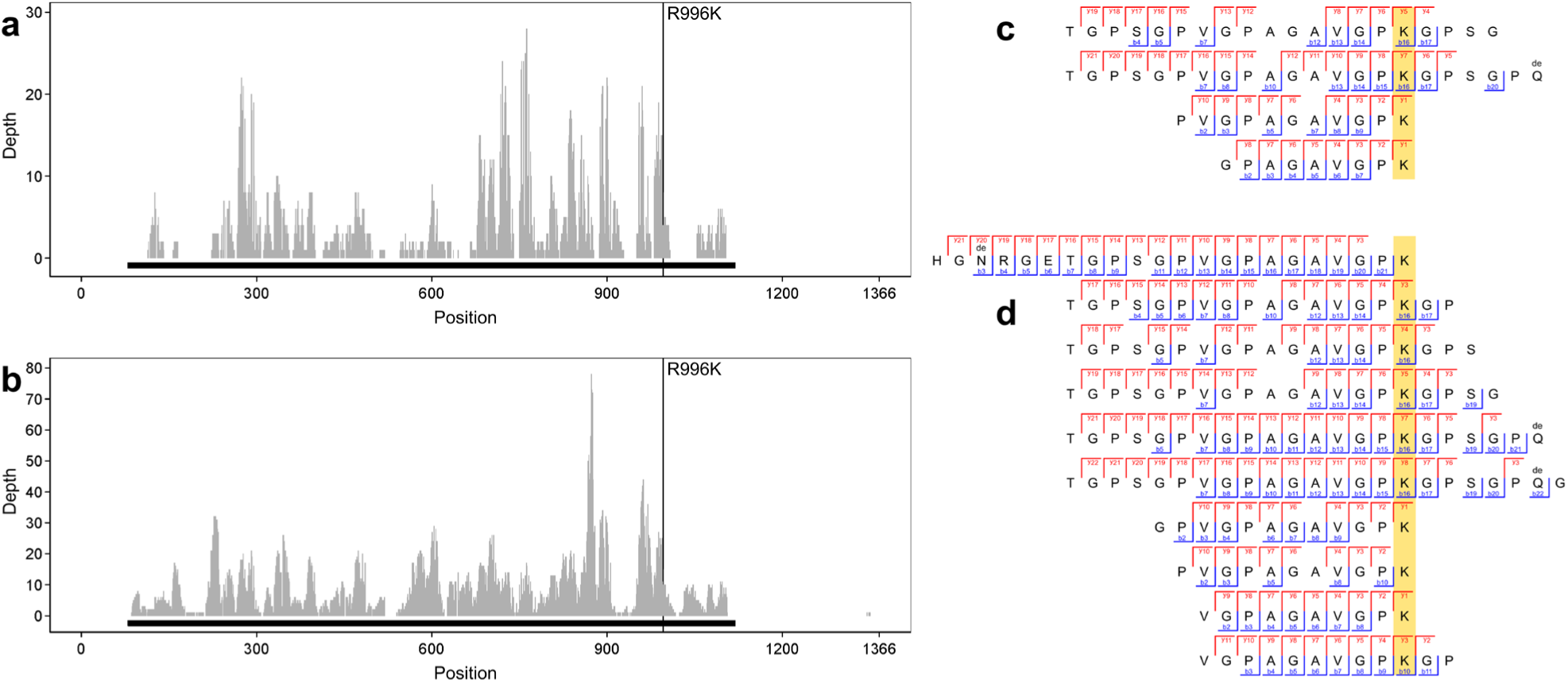
Details of the Denisovan-specific peptides identified from COL1A2 of Penghu 2 and Penghu 3. The depth and amino acid coverage of Penghu 2 (**a**), Penghu 3 (**b**) based on computationally validated peptide counts. The regions of the mature protein without the signal sequence and propeptide are indicated with the black solid bars under the histograms. The position with Denisovan-specific variants in COL1A2 is indicated by a vertical solid line. Supporting peptides of the Denisovan-specific residue (R996K) in COL1A2 of Penghu 2 (**c**) and Penghu 3 (**d**), aligned and highlighted for the subject position.

**Extended Data Fig. 4.**
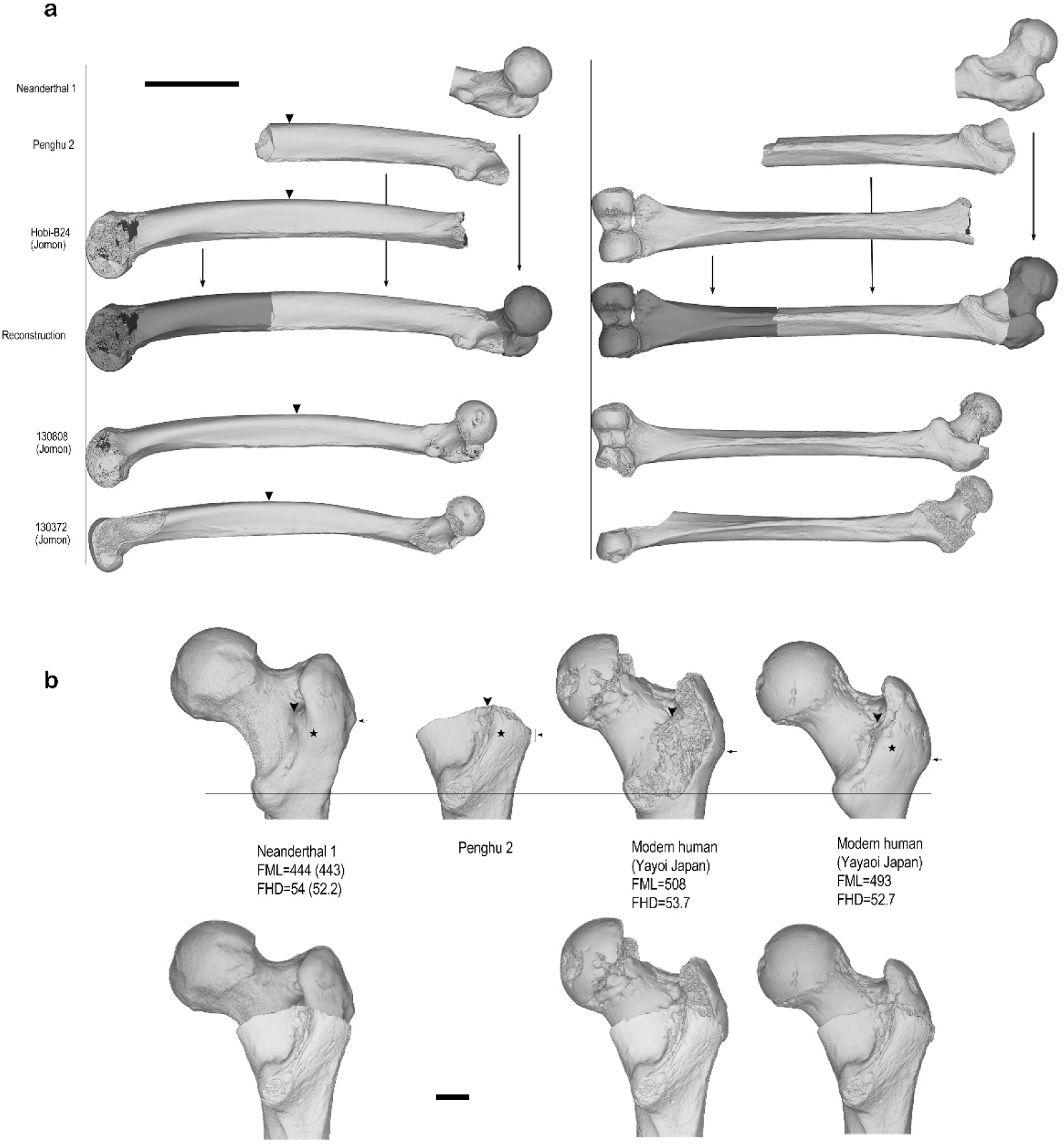
Process of the length estimation of Penghu 2. **a**, Reconstruction of the missing parts. The missing distal portion was digitally reconstructed using Hobi-B24, an extremely robust modern human femur from the Holocene Jomon hunter-gatherers of Japan, which shows markedly similarities to Penghu 2 in shaft dimensions, curvature, and pilastric development. After aligning Hobi-B24 in standard orientation [Ruff], Penghu 2 was fitted to Hobi-B24 using Meshlab, with final fine adjustment to align the posterior borders of pilaster and the anterior vertices of the two midshafts (indicated by inverted triangles). Two other Jomon femora (130808 and 130372) exhibit comparable pilaster development, but we did not use them due to differences in size and general morphology. Scale bar=10 cm. **b**, Posterior views of femoral neck of Penghu 2 and selected large hominin femora (upper) and reconstructions of the proximal portion of Penghu 2 using these specimens (lower). FML and FHD (femoral head diameter) for each specimen are indicated in mm. Neanderthal 1 is a cast of the left femur, with the dimensions from the original specimen [T&R] in the parentheses. The specimens were aligned at the centers of their lesser trochanters (the transverse line). Large arrows=base (posterior edge) of trochanteric fossa. Small arrows=lateral projection of the greater trochanter (possible position is indicated for the damaged Penghu 2). Stars=center of the bulge on the intertrochanteric crest. Scale bar=2 cm. Note large vertical distances of these landmarks from the lesser trochanter in Penghu 2. Reconstructions in the lower row were performed by aligning the bases of the trochanteric fossae, although minor gaps at the junctions leave the possibility that the proximal end of Penghu 2 was slightly larger than all or most of these reconstructions. Thus reconstructed three mesh models of Penghu 2 yielded FML and FBL of 492.6 and 455.8 mm (Yayoi left), 488.8 and 461.4 mm (Yayoi right) and 490.9 and 466. 2 mm (Neanderthal), respectively.

**Extended Data Fig. 5.**
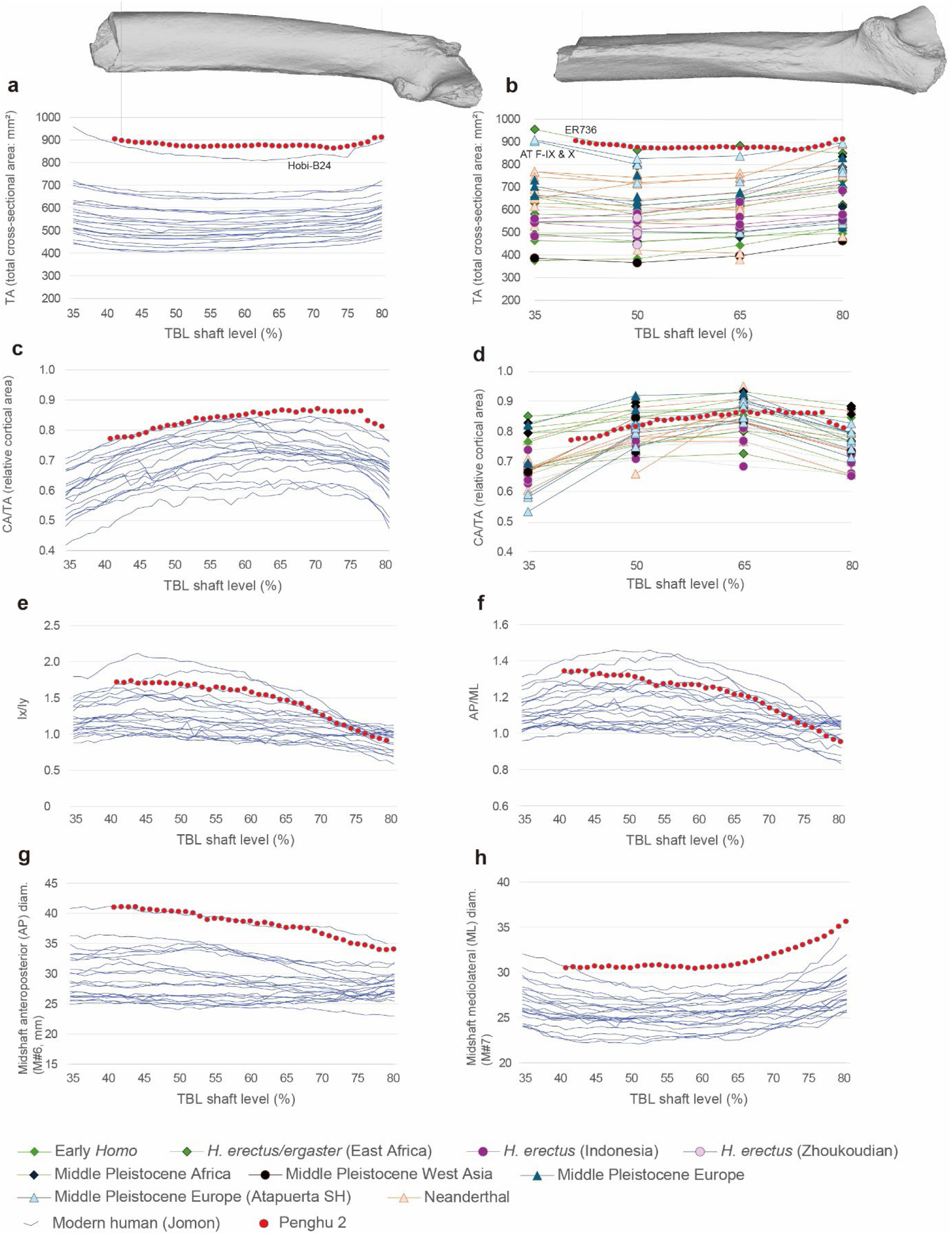
Femoral sequential cross-sectional metrics of Penghu 2, modern human hunter-gatherers (Jomon) and pre-modern *Homo* specimens. Penghu 2 and Jomon are sampled at 1% and the other fossil specimens at 15% intervals between 20% (distal) and 80% (proximal) of femoral biomechanical length (FBL). The values of the distalmost (far left) point of Penghu 2 are based on reconstruction using molding clay for the missing small part. Based on the reconstruction described in the text and in Extended Data Fig. 4, the distal break of Penghu 2 (the level indicated by the left vertical line) is assumed to have occurred at 42% FBL shaft level. No clear evidence is found in the panels in this figure to suggest that the above is an overestimate and the actual length of Penghu 2 was shorter. TA and ML tend to increase and AP/ML and Ix/Iy decrease when approaching the distal articular surface, but no such trend is seen in Penghu 2, although the distally tapering ML width of Penghu 2 affects these trends to some extent. In particular, the robust archaic *Homo* specimens with similar TA values to Penghu 2 (KNM-ER 736, Atapuerca SH-FIX and FX) show substantial increase in TA from 50 to 35% levels. If Penghu 2 shared this feature, its broken distal end was not very close to 35% level.

**Extended Data Fig. 6.**
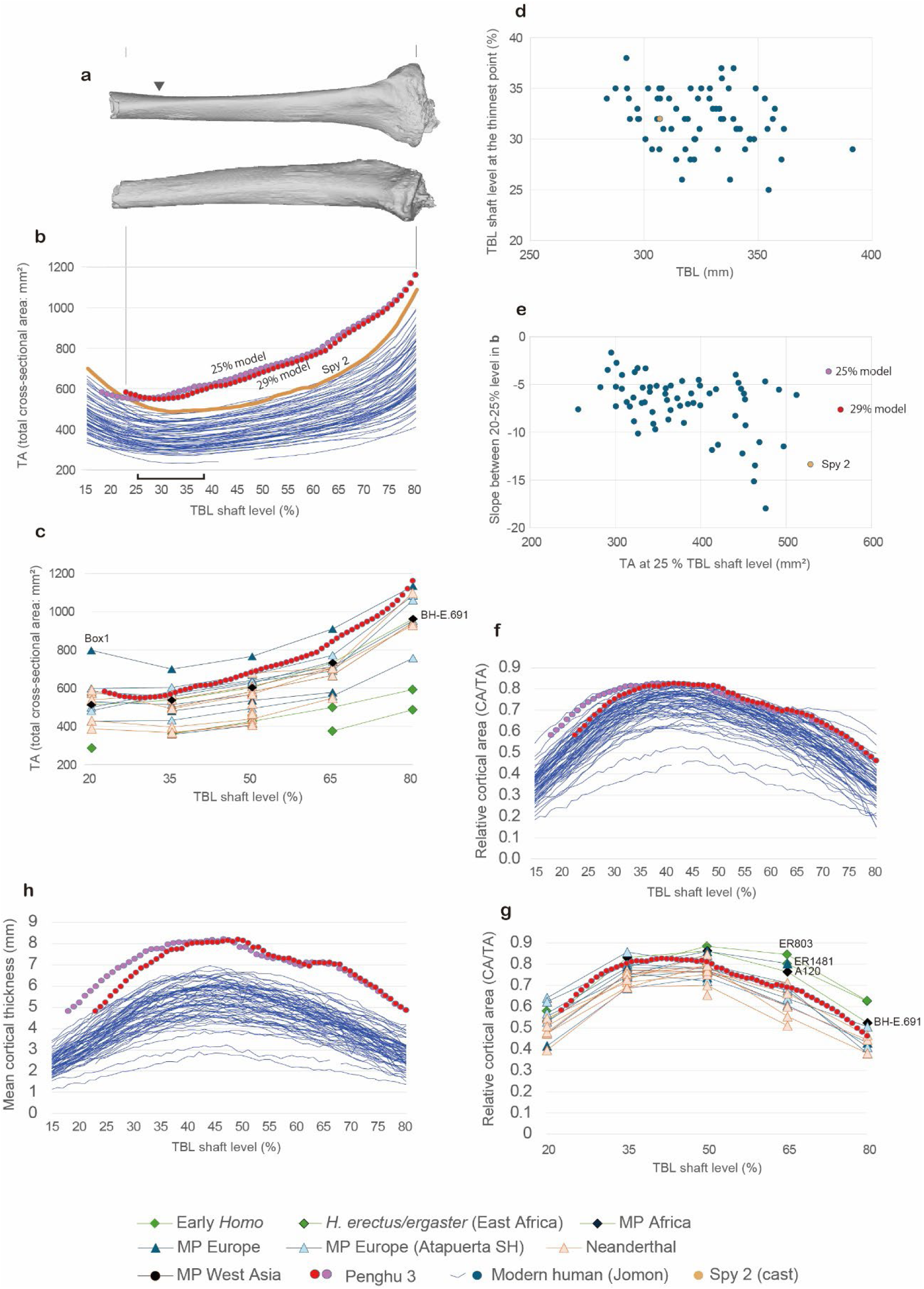
Process of length estimation of Penghu 3. **a**, Anterior (upper) and medial (lower) views of Penghu 3 aligned in the standard orientation (Ruff) by fitting it to an aligned, long modern human tibia (TML=415 mm). The triangle indicates the position of minimum total cross-sectional area (TA). **b** and **c**, TA sampled at 1% (b) and 15% (c) intervals between 20% (distal) and 80% (proximal) of tibial biomechanical length (TBL) for extant (b) and premodern (c) *Homo*. The transverse bar at the bottom in b is the range of minimum thickness positions for modern humans and Spy 2 (25−38%). **d**, Relationship between TBL and the level of the minimum TA. **e**, Relationship between shaft thickness and thickness transition (slope) in the distal segment of the shaft. **f** and **g**, Cortical area (CA) relative to TA along TBL, sampled at 1% (f) and 15% (g) intervals. **h**, Mean cortical thickness in each section sampled at 1% interval of TBL.

**Extended Data Fig. 7.**
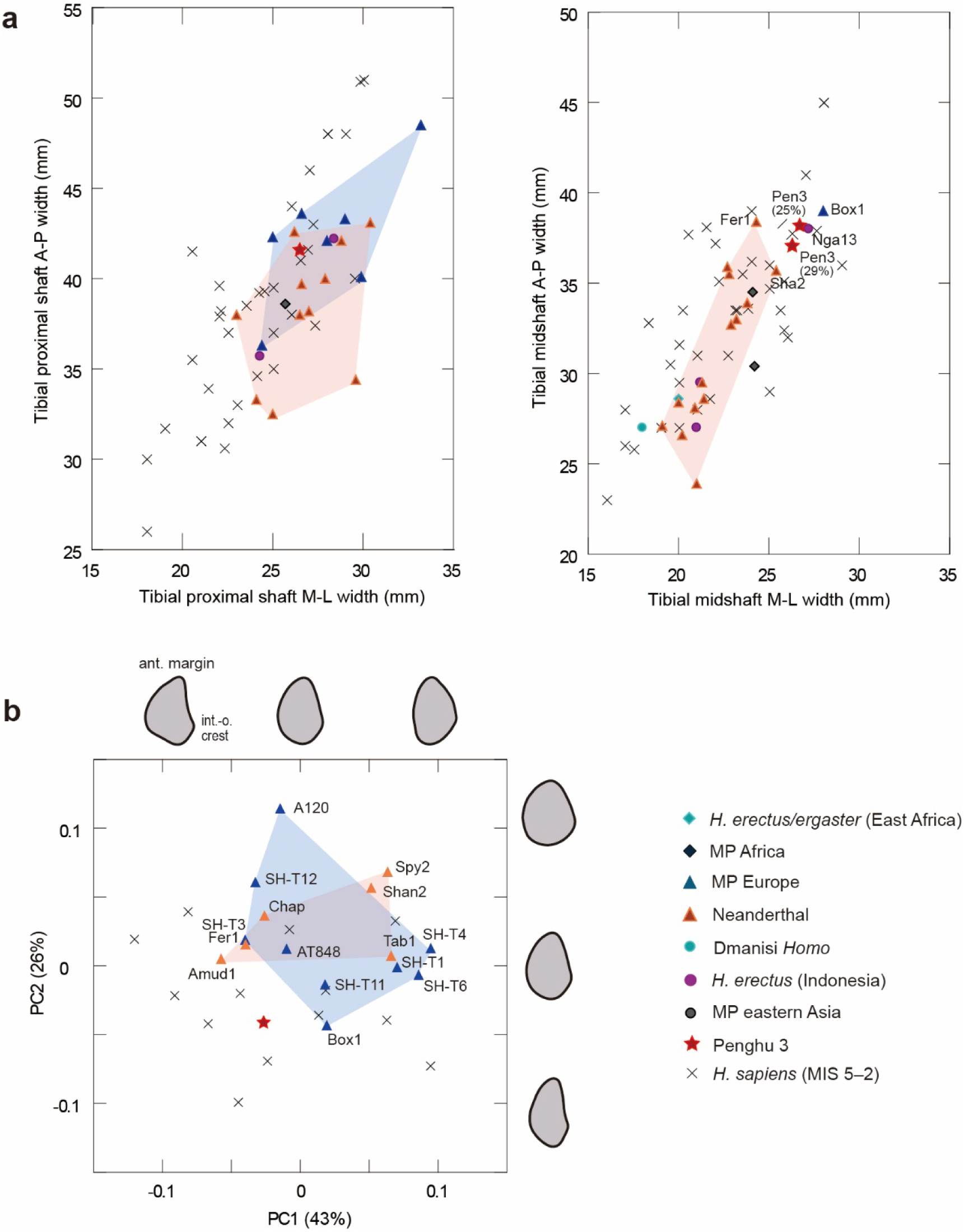
Metric comparisons of tibial shafts. **a**, Scatter plots of anteroposterior (A-P) and mediolateral (M-L) diameters at the proximal shaft (at nutrient foramen) and midshaft (50% TBL shaft level). Values based on the 29% and 25% model reconstructions of Penghu 3 (Extended Data Fig. 6) are indicated for the midshaft analysis. **b**, Plots of PC (principal component) scores derived from the normalized Elliptic Fourier Analyses (EFAs), based on midshaft cross-sectional contours. Penghu 3 is based on the 29% model. Proportions explained by each PC are in parentheses. Shape differences along PC axes are shown on right tibiae, with anterior directed up and lateral right, two standard deviations from the origin. Convex hulls are depicted for some archaic *Homo* groups with larger sample sizes.

**Extended Data Table 1.**
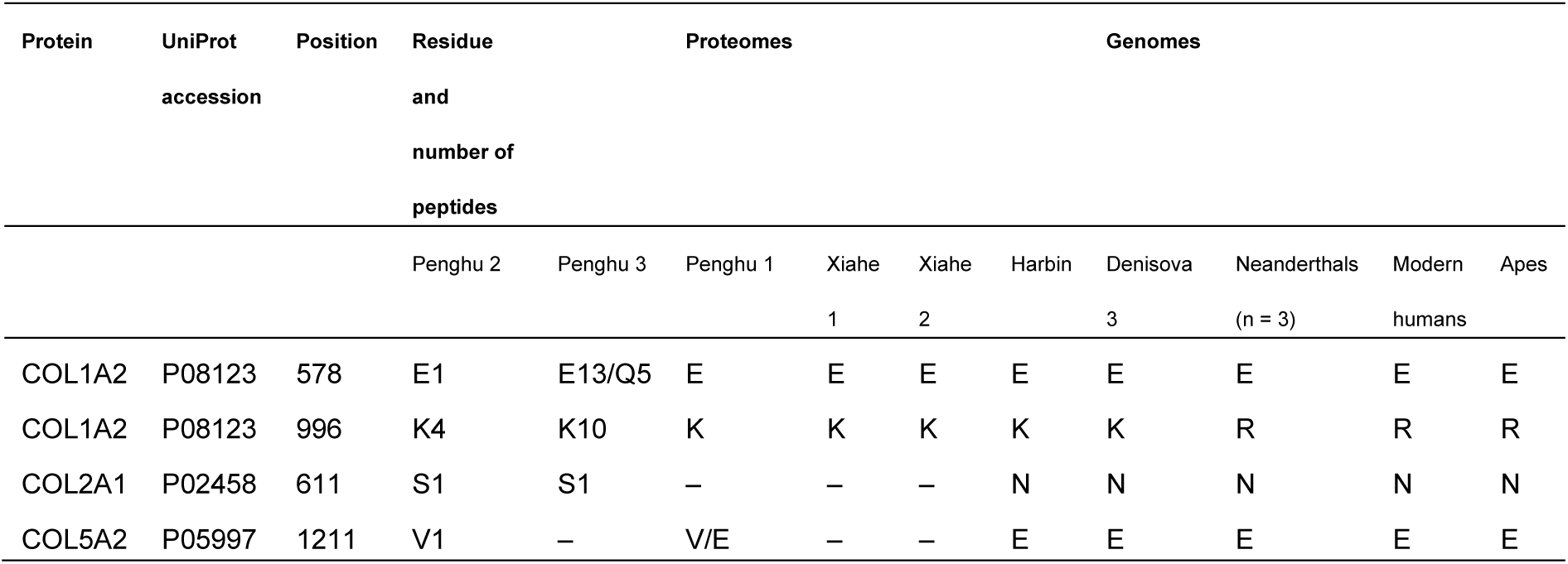
Denisovan-specific or related variants identified in this study. Two different AA residues that are shown with a hyphen mean heterozygosity. Numbers shown with some AA residues mean the depth of the non-redundant peptides that cover the subject position in the Penghu 2 or Penghu 3. The corresponding amino acid residues in other Denisovan individuals, Neanderthals, modern humans, and apes are also shown.

**Extended Data Table 2.** Measurements of Penghu 2 and 3, and metric comparisons with Neanderthals. The numerals in the left column are Matrin’s number. For Penghu 3, our best and minimum estimates (in parentheses) of the length, as well as midshaft dimensions based on these figures, are reported. Z scores are deviations from the Neanderthal means relative to the standard deviations for Neanderthal. Data of Neanderthal are from refs [40,72].

|  | Penghu |  | Neanderthal |  |  |
| --- | --- | --- | --- | --- | --- |
|  |  | Z-score | M | Mean | S.D. |
| **Penghu 2 (femur)** |  |  |  |  |  |
| 1 Maximum length | (491) | 1.93 | 13 | 429.23 | 32.04 |
| Biomechanical length | (461) |  |  |  |  |
| 6 Midshaft AP diameter (50%) | 40.3 | 2.78 | 17 | 29.49 | 3.89 |
| 7 Midshaft ML diameter (50%) | 30.5 | 0.46 | 17 | 29.29 | 2.61 |
| 8 Midshaft circumference (50%) | 121 |  |  |  |  |
| Midshaft AP diameter (41%) | 41.1 |  |  |  |  |
| Midshaft ML diameter(41%) | 30.5 |  |  |  |  |
| 10 Proximal AP diameter | 31.0 | 1.30 | 16 | 26.94 | 3.13 |
| 9 Proximal ML diameter | 35.5 | 0.70 | 16 | 33.25 | 3.21 |
| Midshaft cross-sectional total area (TA50%) | 875.9 | 2.04 | 14 | 670.43 | 100.83 |
| **Penghu 3 (tibia)** |  |  |  |  |  |
| 1a Maximum length | 432 (>411) | 4.47 (>3.60) | 7 | 325.00 | 23.92 |
| Biomechanical length | 408 (>387) |  |  |  |  |
| 3 Proximal max. breadth | 88.4 | 1.19 | 4 | 81.50 | 5.78 |
| 3a Medial condyle breadth | 39.1 |  |  |  |  |
| 3b Lateral condyle breadth | 37.1 |  |  |  |  |
| 4a Medial condyle depth | 54.1 |  |  |  |  |
| 4b Lateral condyle depth | 49.5 |  |  |  |  |
| Tuberosity projection (Trinkaus) | 48.2 | 0.44 | 5 | 44.83 | 7.72 |
| 8 Midshaft AP diameter | 37.2 (<38.2) | 1.30 (<1.57) | 12 | 32.19 | 3.84 |
| 9 Midshaft ML diameter | 26.3 (<26.7) | 2.33 (<2.57) | 12 | 22.41 | 1.67 |
| 10 Midshaft circumference | 107 |  |  |  |  |
| 8a Proximal AP diameter | 41.6 | 0.80 | 10 | 38.86 | 3.44 |
| 9a Proximal ML diameter | 27.0 | -0.05 | 10 | 27.10 | 2.19 |
| 9aV Proximal ML diameter | 27.0 |  |  |  |  |
| 10b Distal min. circumference | 86 |  |  |  |  |
| 12 Medial retroversion angle | 16 |  |  |  |  |
| Midshaft cross-sectional total area (TA50%) | 684.8 (<703.8) | 1.78 (<1.99) | 11 | 524.91 | 89.91 |

**Extended Data Table 3.** Stature and body mass estimates for Penghu and other Pleistocene hominins from mainland eastern Asia. ^a^Based on Ruff’s formulae^73^ developed for Holocene Europeans, using the models for femur (F), tibia (T), femur and tibia (F+T), males (m) and females (f) of North (N) or South (S) of Europe.^b^Based on Sjøvold’s formulae^74^ designed for extant humans of all ethnic groups and both sexes. ^c^Based onRuff’s formulae developed using worldwide modern humans^21^, using the models for femoral midshaft cortical area (FCA), femoral maximum length (FL), tibial plateau mediolateral breadth (TPML), femoral head superoinferior breadth (FHSI), FHSI estimated from the acetabular breadth (FHSI’), femoral distal articular breadth (FDML) and TPML estimated from FDML (TPML’). ^d^The minimum estimates are in parentheses. ^e^Similar values are reported previously using different methods (168.8 cm, ∼78.6kg)^41^. ^f^Cited from ref. 75. ^g^Cited from ref. 45.

|  | Bone | Latitude | Age | Sex | Stature A<sup>a</sup> | model<sup>a</sup> | Stature B<sup>b</sup> | Body mass<sup>c</sup> | model<sup>c</sup> |
| --- | --- | --- | --- | --- | --- | --- | --- | --- | --- |
|  |  |  | (ka) |  |  |  |  |  |  |
| **Penghu** |  |  |  |  |  |  |  |  |  |
| Penghu 2 | femur | 23°N | 190-130 | M? | 176.4 | F: N+S, m | 178.9 | 82.5 | FCA+FL |
|  |  |  | or 70-10 |  |  |  |  |  |  |
| Penghu 3 | tibia | 23°N | 45 | M? | 180.9 (>175.0)<sup>d</sup> | T: S, M | 189.5 (>182.6)<sup>d</sup> | 90.8 | TPML |
| ***H. erectus* (northern China)** |  |  |  |  |  |  |  |  |  |
| Zhoukoudian Femur I | femur | 39°N | 780-680 | M? | 152.4 | F: N+S, m+f | 155.3 | 62.9 | FCA+FL |
| Zhoukoudian Femur II | femur | 39°N | 780-680 | F? |  |  |  | 60.7 | FCA |
| Zhoukoudian Femur IV/V | femur | 39°N | 780-680 | M? | 154.3 | F: N+S, m+f | 157.2 | 60.6/61.3 | FCA+FL/FCA |
| Zhoukoudian Femur VI | femur | 39°N | 780-680 | M? |  |  |  | 62.9 | FCA |
| **MP archaic *Homo*** |  |  |  |  |  |  |  |  |  |
| Hualongdong 11 | femur | 30°N | 300 |  |  |  |  | 60.8 | FCA |
| Jinnuishan 1 | pelvis, ulna | 40°N | 200 | F |  |  | 168.8<sup>e</sup> | 74.2<sup>e</sup> | FHSI' |
| ***H. sapiens* (mainland including the Pleistocene Sundaland)** |  |  |  |  |  |  |  |  |  |
| Tianyuan 1 | femur, tibia | 39°N | 40 | M? | 168.4/169.9 | F+T: S/N, m |  | 73.4 | FHSI |
| Zhoukoudian-UC 67 | femur | 39°N | 36 |  |  |  |  | 67.5 | FCA |
| Zhoukoudian-UC 68 | femur | 39°N | 36 |  |  |  |  | 60.0 | FCA |
| Zhoukoudian-UC 105 | femur | 39°N | 36 |  |  |  |  | 69.9 | FCA |
| Zhoukoudian-UC 117 | femur | 39°N | 36 |  |  |  |  | 58.6 | FCA |
| Liujiang | femur | 24°N | 28 | M |  |  |  | 50.9 | FHSI |
| Maomaodong GM7506 | femur | 26°N | 15 |  |  |  |  | 63.6 | FCA |
| Maomaodong GM7507 | femur | 26°N | 15 |  |  |  |  | 59.8 | FCA |
| Maomaodong GM7508 | femur | 26°N | 15 |  |  |  |  | 59.8 | FCA |
| Niah | femur | 4°N | 35 | F | 146<sup>f</sup> |  | 151.6 | 44.6 | FDML, TPML' |
| Trinil 1 | femur | 7°S | ? |  | 168.5 | F: S, m+f | 171.1 | 62.6 | FHSI |
| Trinil 9 | femur | 7°S | 36 | M | 156<sup>g</sup> |  | 158.9 | 55.4 | CAFL |
| Wajak W-H-27 | femur | 8°N | 33 |  |  |  |  | 65.4 | FHSI |

## Notes

### Competing Interest Statement

The authors have declared no competing interest.

