## Supplementary Information for "Denisovan leg bones from Taiwan reveal large body size"

This PDF file includes:

Supplementary Notes 1 to 8
and associated Supplementary Figures
Supplementary Tables
Supplementary references

**Supplementary Note 1:**
**Research history of Penghu 2 and 3**

Soon after we (C-H.C., Y.K., M.T. and R.T.K.) started our work on the Penghu 1 mandible that derived from an antique shop in Tainan City, we identified Penghu 2 and 3 in October 2010 at the National Museum of Natural Science of Taiwan located in Taichung City (hereafter 'NMNS Taiwan'). These hominin leg bones were among a large Penghu fossil collection collected by Mr. Li-Ren Hou, a local artist, who donated it to the NMNS Taiwan, in 2011. Following this discovery, we initiated a stepwise research program in 2012 to analyze these hominin leg bones as summarized below, alongside our investigations of Penghu 1, which were published in 2015[1] and 2025[2].

In 2012, we cut off a small piece of bone from the distal end of each leg bone (as well as the central part of the Penghu 1 mandible) using a thin diamond cutter at the National Museum of Nature and Science, Tokyo (hereafter 'NMNS Tokyo'). We submitted the separated fragments to laser- ablation U-series dating, the results of which were recently published in ref. 3. Based on the bone powder generated from this cutting, we also examined organic contents and obtained negative and positive results for Penghu 2 and 3, respectively. In 2015, at NMNS Taiwan, we took additional bone samples of Penghu 2 and 3 by drilling the sections created by the 2012 work. We also collected bone samples from non-human mammalian fossils from the Penghu Channel at that opportunity. Based on these samples, in 2017, we obtained the radiocarbon dates we mentioned in the present paper[4]. This positive result for Penghu 3 prompted us to collect additional bone samples from these fossils in Tokyo for palaeoproteomic analysis and ancient DNA extraction in 2019 and 2024. The palaeoproteomic results are reported in the present paper, and the outcome of the ancient DNA analyses will be reported elsewhere by a team of The University of Tokyo and NMNS Taiwan.

For morphological analyses, we performed μCT scans of the two leg bones at The University Museum, The University of Tokyo, in 2024. Because we found that it is difficult to digitally separate the attached nodules from the bone surfaces due to ambiguous boundaries in the CT scan, we physically removed these nodules at the NMNS Taiwan in 2025.

### Supplementary Note 2:
### Optimization of protein extraction

We evaluated protein extraction methods and mass spectrometry measurement parameters using a faunal specimen from Penghu to maximize the coverage of phylogenetically informative sites in hominin fossils. Protocols for protein extraction and digestion have already been established in a previous study using faunal fossil specimens from Penghu[2]. In this study, based on the optimal protocol established in the previous study, we investigated (i) the compatibility of the non-trypsin protease (Glu-C) with the presence of denaturant (Gu), and (ii) the optimization of measurement parameters (fragmentation mode and MS2 resolution) in a different mass spectrometry platform. The number of razor+unique peptides (R+U peptides) and identified amino acid (AA) residues were used as proxies for comparison.

Denaturants disrupt the three-dimensional structure of proteins and thereby increase protein extraction efficiency, but if they remain in the solution, they reduce protease activity and digestion efficiency. Previous studies have shown that trypsin retains its activity in the presence of the denaturant guanidine (Gu) at low concentrations[5]. The same was true in earlier research using fossils from Penghu[2]. However, for proteases other than trypsin, compatibility with denaturants has not been properly verified. It is known that, when used in combination with trypsin, Glu-C can expand the proteome coverage of fossil bones, including those from Penghu [2,6]. Another previous study showed that Glu-C retains its activity even in the presence of denaturants[7]. In this study, in order to determine the optimal buffer for digestion using Glu-C, we investigated the extent to which Glu-C retains its activity in the presence of denaturants when applied to fossil proteomes.

The Orbitrap Fusion Tribrid mass spectrometer (Fusion) is equipped with both an ion trap and an Orbitrap as detectors, allowing flexible modification and combination of measurement parameters such as MS/MS spectral acquisition and fragmentation modes[8]. In this study, measurements were performed using the Fusion, whereas previous work used an Orbitrap Exploris 480 to obtain ancient proteomes from Penghu fossil bones[2]. Here, to achieve high proteome coverage on the Fusion, we investigated multiple fragmentation modes and MS2 mass resolution settings, which are known to strongly affect amino acid sequence acquisition, and examined the optimal combination of parameters. Since the data obtained from the ion trap detector is not well suited to certain parts of database searches, we decided to use only Orbitrap detectors.

#### Provenance

A fossil specimen, mandible of *Sus sp.* (NTUM-VP 210121), was also collected and dredged from the sea bottom of the Taiwan Strait by a local commercial fishery during the trawling operations. The ages of these terrestrial fossils remain uncertain, but they likely coexisted with Penghu hominins, given their shared source locality. The faunal specimen was subsequently donated to CHT in 2021 for further research and permanent curation at the National Taiwan University.

### Materials and methods

The mandible specimen of *Sus sp.* from Penghu Channel was used to optimize protein extraction methods and mass spectrometry measurement parameters. The specimen is curated at the National Taiwan University and was used with permission from the institution. Samples were cut from the specimen using diamond disks and a dental drill after surface abrasion to remove exogenous contaminants. The bone sample was cut from the bottom of the middle part of the mandibular body, in the same position shown as "Bone 2" (PF02B2) in ref. 2. The samples were then crushed into a coarse powder with a metal hammer and weighed into protein LoBind tubes. All procedures were conducted in a clean laboratory dedicated to ancient biomolecule analysis at SOKENDAI. All fractions were treated along with experimental blanks that did not contain any sample.

#### *Compatibility of Glu-C with Gu*

Bone samples (~5 mg or ~10 mg) were decalcified with 500 µL of 0.5 M EDTA for 2–3 days under rotation. The samples were then centrifuged (10,000 ×g, 5 min). Since Tsutaya et al.[2] already demonstrated higher proteome recovery with the direct application of Glu-C to the EDTA solution, the supernatant was not used for method optimization in this study. The pellet was washed three times with 100 µL of 50 mM Tris solution, centrifuged each time, and the supernatants were combined with the EDTA supernatant. Then, 50 µL Gu buffer (2 M guanidinium chloride, 10 mM tris(2-carboxyethyl)phosphine [TCEP], 20 mM chloroacetamide [CAA], and 100 mM Tris) or TEAB buffer (50 mM triethylammonium bicarbonate [TEAB], 10 mM TCEP, and 20 mM CAA) was added to the pellet fractions. The pellet fraction was incubated at 60°C for 2.5 hours. The samples with Gu buffer were then diluted with a solution of 50 mM TEAB to reduce the concentration of guanidinium chloride to less than 0.2M. Subsequently, 0.4 µg of Glu-C (the approximate weight ratio of >1:64 relative to the total protein) was added and incubated overnight at 37°C. After digestion, the solution was acidified to a pH of less than 2 to inactivate the protease. In this experiment, we examined three conditions: 5 mg of bone with Gu buffer, 10 mg of bone with Gu buffer, and 10 mg of bone with TEAB buffer.

Digested samples were purified and desalted using in-house-made StageTips with two stacked C18 membranes (CDS Analytical, USA)[9], with the same procedures described in the Supplementary Method (**Supplementary Note 7**).

Purified peptides were measured under the same conditions described in the Supplementary Method (**Supplementary Note 7**).

The resulting MS/MS spectra were searched against the entire pig proteome or the skeletal proteome using MaxQuant, version 2.0.3.0[10]. The reference proteome (UP000694727) of pig (*Sus scrofa*) was obtained from UniProt (as of 2023-05-13) and used for the analysis as the pig proteome database. A subset of 32 proteins (AHSG, ALB, APOA1, BGLAP, BGN, C3, CHAD, CLEC3B, COL10A1, COL11A1, COL11A2, COL12A1, COL1A1, COL1A2, COL22A1, COL2A1, COL3A1, COL5A1, COL5A2, COL5A3, COL9A1, DCN, F10, F2, LUM, MGP, OMD, POSTN, SERPINF1, SPARC, SPP1, THBS1) that are typically identified from ancient bone/dentine samples was curated from the pig proteome (UP000694727, as of 2023-01-14) as the pig skeletal database. The following parameters were used for the analysis.

Parent mass error and fragment mass tolerances were those pre-set for Orbitraps. Carbamidomethylation was set as a fixed modification. Oxidation of methionine, deamidation of asparagine and glutamine, hydroxyproline, and derivation of pyroglutamic acid were set as variable modifications, with up to a maximum of 5 modifications per peptide allowed. Digestion efficiency was selected as either "Specific" or "Semi-specific" with a maximum of 2 missed cleavages with the pig proteome database, and "Unspecific" with the pig skeletal database. The specific search only accepts peptides that were cleaved at both N- and C-ends by the specific protease, while the semi-specific search accepts peptides that were cleaved at either the N- or C-end by the specific protease. The unspecific search does not require specific cleavage for peptides to be identified. All peptides were automatically filtered by a false discovery rate (FDR) of 1.0%. Contaminant accessions (i.e., keratins and trypsin) were excluded from further analysis using the contamination.fasta file provided by MaxQuant, which includes common laboratory contaminants. The protein groups identified with non-overlapping ≥2 razor+unique peptides were considered to be present in the samples. The number of AAs was calculated by multiplying the sequence length by razor+unique sequence coverage for convenience.

*Mass spectrometry measurement parameters*

Bone samples (~5 mg) were processed with the digestion-free method, and proteins were purified with C18 StageTips with the procedures described in the Supplementary Method (**Supplementary Note 7**).

LC-MS/MS analysis was basically performed under the same conditions described in the Supplementary Method (**Supplementary Note 7**). For optimization of the fragmentation mode, we evaluated two conditions: collision-induced dissociation (CID) and higher energy collisional dissociation (stepped HCD). For the MS2 mass resolution, we tested two settings: 15,000 full width at half maximum (15k) and 60,000 full width at half maximum (60k). Using samples prepared from the same specimen PF02B2 and processed with the same digestion-free protocol, we conducted four measurements combining fragmentation mode and MS2 resolution (CID-15k, CID-60k, HCD-15k, and HCD-60k).

The resulting MS/MS spectra were searched against the pig skeletal database described above using MaxQuant, version 2.6.3.0[10]. The following parameters were used for the analysis. Parent mass error and fragment mass tolerances were those pre-set for Orbitraps. Oxidation of methionine, deamidation of asparagine and glutamine, hydroxyproline, and derivation of pyroglutamic acid were set as variable modifications, with up to a maximum of 5 modifications per peptide allowed. Digestion efficiency was selected as "Unspecific". All peptides were automatically filtered by a false discovery rate (FDR) of 1.0%. Contaminant accessions (i.e., keratins and trypsin) were excluded from further analysis using the contamination.fasta file provided by MaxQuant, which includes common laboratory contaminants. The protein groups identified with non-overlapping ≥2 razor+unique peptides were considered to be present in the samples. Each AA residue was evaluated with corresponding PSMs reported in the MaxQuant output msms.txt files by using an R script[11], and unsupported positions were replaced with "X". The number of AAs was calculated from the MaxQuant output peptides.txt files. The PSM-supported residues were calculated as "checked" AA residues/sequences, and entire identified residues that are not necessarily supported by MS2 spectra were calculated as "unchecked" residues/sequences. We examined peptides with a peptide score of ≥70 and that were derived from proteins considered endogenous rather than contaminants.

**Compatibility of Glu-C with Gu**

Under conditions where the denaturant was removed for Glu-C digestion (TEAB buffer condition), the number of razor+unique peptides and AAs obtained was about twice that under conditions where the denaturant was not removed (Gu buffer condition) in the specific and semi-specific searches, and about 1.6 times higher in the unspecific search (**Supplementary Table 2_1**). When collagen and NCP were evaluated separately, the largest numbers of razor+unique peptides (**Supplementary Fig. 2_1**) and AA residues (**Supplementary Fig. 2_2**) were generally identified under conditions in which the denaturant was removed. Therefore, we decided to apply Glu-C under the condition of complete removal of Gu for Penghu 2 and Penghu 3.

**Mass spectrometry measurement parameters**

Based on analysis of AA residues supported by MS2 spectra, both the numbers of razor+unique peptides (**Supplementary Fig. 2_3**) and AA residues (**Supplementary Fig. 2_4**) were highest when the fragmentation mode was set to CID and the MS2 resolution to 60k (**Supplementary Table 2_2**). Whereas MS2 resolution at 60k produced markedly better results than at 15k, the choice of fragmentation mode had a much smaller impact. The same trend was observed when collagen and NCP were examined separately, as well as when we evaluated the total number of AA residues, including positions not necessarily supported by MS2 spectra (**Supplementary Fig. 2_4**). Therefore, we decided to operate Fusion with CID fragmentation mode and an MS2 resolution of 60k.

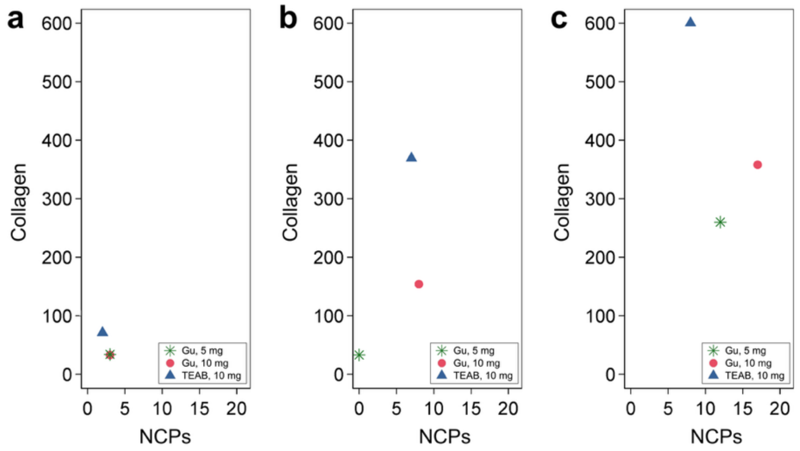


**Supplementary Fig. 2_1. The number of identified razor+unique peptides of collagen and NCPs from PF02B2 in different protein extraction methods.** Results obtained in specific (a), semi-specific (b), and unspecific (c) MaxQuant searches are shown.

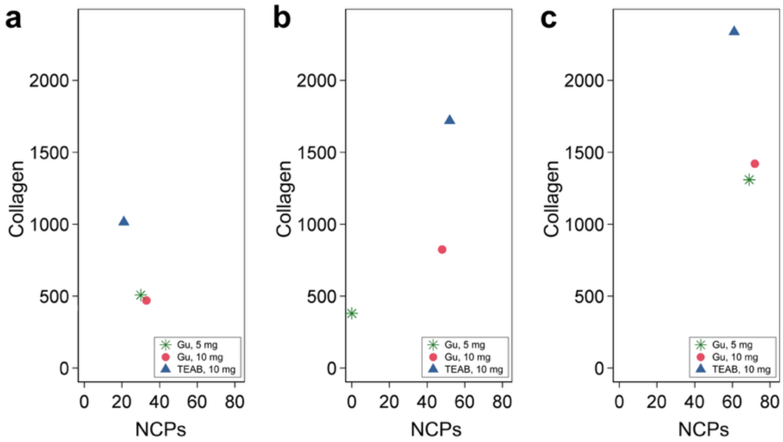


**Supplementary Fig. 2_2. The number of identified AA of collagen and NCPs from PF02B2 in different protein extraction methods.** Results obtained in specific (a), semi-specific (b), and unspecific (c) MaxQuant searches are shown.

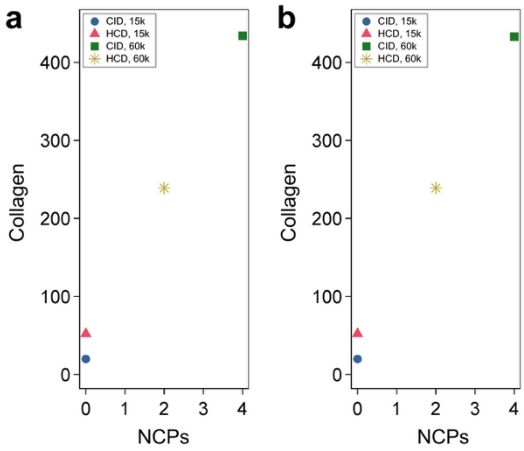


**Supplementary Fig. 2_3. The number of identified razor+unique peptides of collagen and NCPs from PF02B2 in different mass spectrometry measurement parameters.** Results obtained from unvalidated (a) and validated (b) AA peptides are shown.

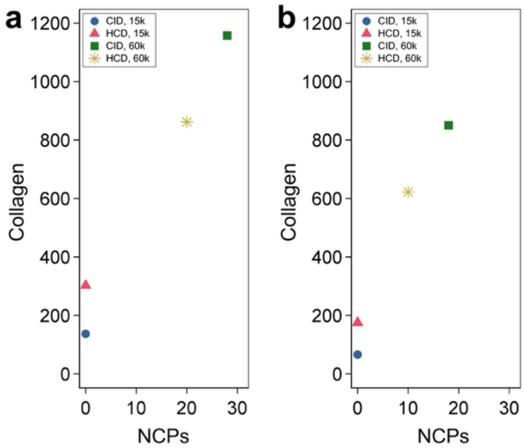


**Supplementary Fig. 2_4. The number of identified AA of collagen and NCPs from PF02B2 in different mass spectrometry measurement parameters.** Results obtained from unvalidated (a) and validated (b) AA peptides are shown.

**Supplementary Table 2_1. Results for the evaluation of the extraction protocols for Glu-C digestion.** Results of buffers used for digestion (Gu or TEAB) and the starting bone amount (approximately 5 mg or 10 mg) were shown for MaxQuant search sessions (specific, semi-specific, or unspecific protease cleavages) and protein types (collagen or NCPs).

| **Search** | **Protein** | **R+U peptides** | | | **AA** | | |
|---|---|---|---|---|---|---|---|
| | | Gu, 5 mg | Gu, 10 mg | TEAB, 10 mg | Gu, 5 mg | Gu, 10 mg | TEAB, 10 mg |
| Specific | Collagen | 34 | 33 | 71 | 507 | 470 | 1014 |
| | NCP | 3 | 3 | 2 | 30 | 33 | 21 |
| | Total | 37 | 36 | 73 | 536 | 503 | 1035 |
| Semi-specific | Collagen | 33 | 154 | 369 | 380 | 824 | 1719 |
| | NCP | 0 | 8 | 7 | 0 | 48 | 52 |
| | Total | 33 | 162 | 376 | 380 | 873 | 1771 |
| Unspecific | Collagen | 260 | 358 | 600 | 1309 | 1420 | 2336 |
| | NCP | 12 | 17 | 8 | 69 | 72 | 61 |
| | Total | 272 | 375 | 608 | 1378 | 1492 | 2397 |

**Supplementary Table 2_2. Results for the evaluation of the mass spectrometry measurement parameters.** Results of the different combinations of fragmentation mode and MS2 resolution are shown.

| **Validated** | **Fragmentation** | **MS2 resolution** | **R+U peptides** | | | **AA** | | |
|---|---|---|---|---|---|---|---|---|
| | | | Collagen | NCPs | Total | Collagen | NCPs | Total |
| No | CID | 15k | 20 | 0 | 20 | 137 | 0 | 137 |
| | HCD | 15k | 52 | 0 | 52 | 302 | 0 | 302 |
| | CID | 60k | 434 | 4 | 438 | 1157 | 28 | 1185 |
| | HCD | 60k | 239 | 2 | 241 | 862 | 20 | 882 |
| Yes | CID | 15k | 20 | 0 | 20 | 66 | 0 | 66 |
| | HCD | 15k | 52 | 0 | 52 | 175 | 0 | 175 |
| | CID | 60k | 433 | 4 | 437 | 850 | 18 | 868 |
| | HCD | 60k | 239 | 2 | 241 | 622 | 10 | 632 |

### Supplementary Note 3:
### Identified proteins from Penghu 2 and Penghu 3

#### Removal of potential contaminants

In the protein identification results, we applied conservative criteria to exclude possible contaminants. We excluded proteins predominantly expressed in skin, muscle, or the nucleus, translation factors, conserved ubiquitous proteins, and homologs of proteases used in the extraction (i.e., Glu-C and trypsin) as possible contaminants. These proteins include keratin, myosin, and ubiquitin, for example.

#### Identified protein set

After the removal of contaminant proteins and proteins with <2 razor+unique peptides, a total of 21 and 36 proteins were identified from Penghu 2 (**Supplementary Table 3_1**) and Penghu 3 (**Supplementary Table 3_2**). These proteins cover a total of 1856 and 3283 amino acid residues supported by tandem mass spectra, out of 21979 (8.4%) and 34309 residues (9.6%) in the full protein coverage of mature regions. The coverage of amino acid residues ranged from 0.2% (COL7A1 of Penghu 2) to 69.9% (COL1A2 of Penghu 3), with a heavy distribution toward lower coverage (i.e., the median was 2.5% in Penghu 2 and 4.0% in Penghu 3).

Overall, the upset plot[12] showed that 55.7% (n = 1033/1856) or 45.1% (n = 1482/3283) of all residues recovered were only identified from a single extraction method for Penghu 2 and Penghu 3, respectively, which demonstrates the increased coverage that can be gained by combining multiple extraction methods (**Supplementary Fig. 3_1**). Only 10.0% (n = 186/1856) or 21.6% (n = 709/3283) of AA residues were identified by all extraction methods in Penghu 2 and Penghu 3, respectively. On the other hand, AA residues identified only by the digestion-free, trypsin, or Glu-C methods accounted for 0.5% (n = 10), 41.6% (n =773), or 13.5% (n = 250) of the 1856 identified AA residues from Penghu 2, respectively. For Penghu 3, AA residues identified only by the digestion-free, trypsin, or Glu-C methods accounted for 1.6% (n = 53), 27.2% (n = 894), or 16.0% (n = 525) of the 3283 identified AA residues, respectively. These results are consistent with previous studies showing that using different sets of proteases increases protein sequence recovery[6].

When considering search sessions using the Bone database, AA residues identified only by specific, semi-specific, and unspecific searches accounted for 6.1% (n = 113), 21.8% (n = 398), and 1.3% (n = 25) of the identified 1856 AA residues from Penghu 2, respectively (**Supplementary Fig. 3_2a**). For Penghu 3, AA residues identified only by specific, semi-specific, and unspecific searches accounted for 3.5% (n = 116), 14.1% (n = 464), and 1.3% (n = 44) of the identified 1856 AA residues (**Supplementary Fig. 3_2b**). These are likely due to the degraded nature of the Penghu 2 and Penghu 3 proteomes and the positive relationship between search space and false discovery rates. The endogenous proteins of Penghu 2 and Penghu 3 were typically fragmented (**Supplementary Note 4**) and exist as fragmented peptides that are not necessarily cleaved at the recognition site of specific proteases at least one end. In such cases, even if the cleavage efficiency is very high, most of the resultant peptides have improper "cleavage" sites and can only be identified from semi-specific or unspecific searches. Additionally, the positive relationship between the size of the search space and false discovery rates could contribute. Peptides “cleaved” at improper sites can be identified in semi-specific and unspecific searches, but increased false discovery rates due to the larger search space hinder the identification of some properly or semi-properly cleaved peptides. This result suggests that the combination of different search specificities increases AA identification for degraded proteomes.

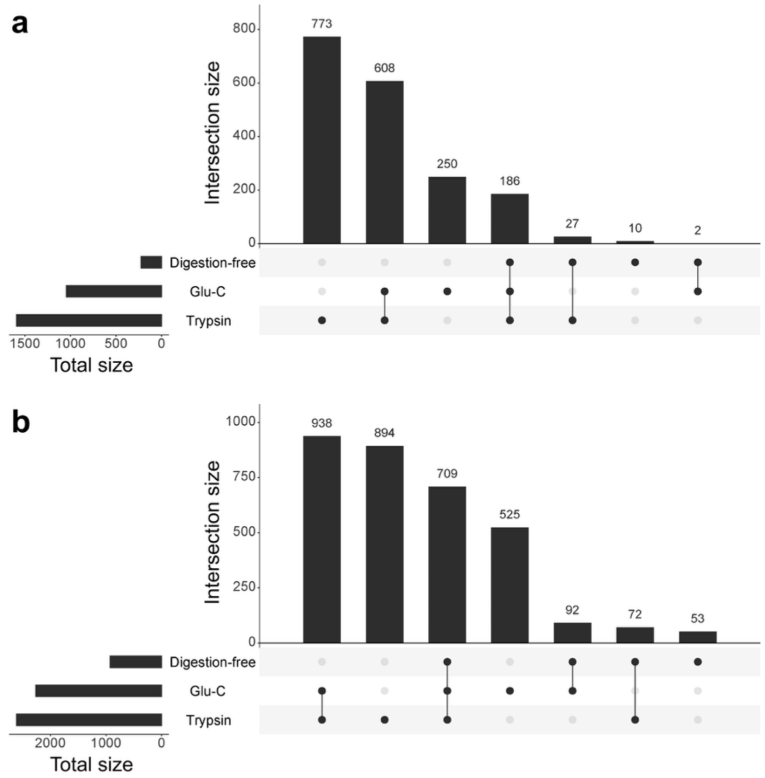


**Supplementary Fig. 3_1**. **Upset plots of the number of AA residues identified from different extraction methods.** Upset plots show from which extraction methods the validated AA residues were obtained in Penghu 2 (a) and Penghu 3 (b).

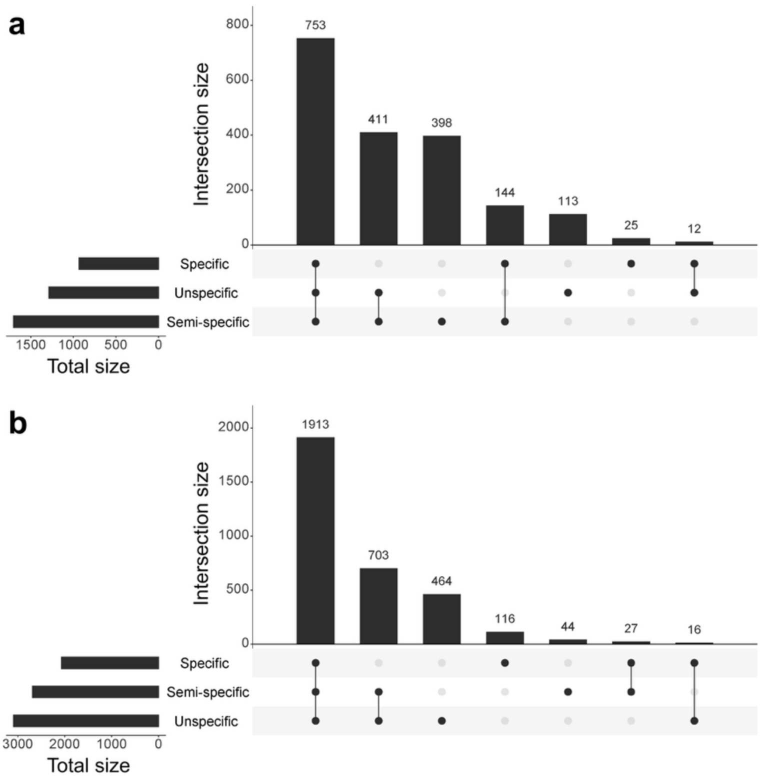


**Supplementary Fig. 3_2. Upset plots of the number of AA residues identified from different MaxQuant search sessions.** Upset plots show from which search sessions the validated AA residues were obtained in Penghu 2 (a) and Penghu 3 (b).

**Supplementary Table 3_1. Summary statistics of the identified proteins from Penghu 2.** The number of razor+unique peptides calculated from non-redundant peptides is shown as "Total", and the sum of those in other fractions does not equal "Total" due to the redundant peptides identified from multiple fractions. The non-redundant depth of the covered AA residues is also shown as "Mean" among the identified residues and as "Total", the sum of the depths of each residue divided by the protein length. The AA coverage (%) was calculated based on the AA sequence of the mature protein that does not contain a signal sequence or a propeptide, rather than the total AA sequence of the preproprotein.

| **Gene name** | **Uniprot ID** | **R+U peptides** | | | | **AA** | **Coverage (%)** | **Length** | | **Depth** | | |
|---|---|---|---|---|---|---|---|---|---|---|---|---|
| | | Total | Trypsin | Glu-C | Digestion-free | | | Total | Mature | Max | Mean | Total |
| COL1A1 | P02452 | 606 | 687 | 234 | 52 | 815 | 55.7 | 1464 | 1057 | 42 | 6.83 | 3.80 |
| COL1A2 | P08123 | 418 | 469 | 202 | 12 | 683 | 50.0 | 1366 | 1040 | 28 | 5.13 | 2.57 |
| COL2A1 | P02458 | 10 | 12 | 5 | 0 | 58 | 3.9 | 1487 | 1102 | 3 | 1.21 | 0.05 |
| COL5A2 | P05997 | 10 | 3 | 9 | 0 | 42 | 2.8 | 1499 | 1203 | 5 | 1.64 | 0.05 |
| COL5A1 | P20908 | 8 | 7 | 4 | 0 | 29 | 1.6 | 1838 | 1568 | 4 | 2.10 | 0.03 |
| COL3A1 | P02461 | 5 | 3 | 2 | 0 | 33 | 2.3 | 1466 | 1068 | 1 | 1.00 | 0.02 |
| CD63 | P08962 | 4 | 0 | 4 | 0 | 21 | 8.8 | 238 | 237 | 2 | 1.38 | 0.12 |
| COL4A3 | Q01955 | 4 | 3 | 1 | 1 | 18 | 1.1 | 1670 | 1642 | 3 | 1.67 | 0.02 |

| | | | | | | | | | | | | |
|---|---|---|---|---|---|---|---|---|---|---|---|---|
| COL12A1 | Q99715 | 3 | 4 | 0 | 0 | 9 | 0.7 | 1366 | 1040 | 2 | 1.44 | 0.01 |
| COL16A1 | Q07092 | 3 | 0 | 3 | 0 | 12 | 0.7 | 1604 | 1583 | 3 | 1.75 | 0.01 |
| EMID1 | Q96A84 | 3 | 3 | 0 | 0 | 13 | 2.9 | 441 | 419 | 3 | 2.31 | 0.07 |
| AHSG | P02765 | 2 | 0 | 2 | 0 | 13 | 3.5 | 367 | 309 | 1 | 1.00 | 0.04 |
| CD36 | P16671 | 2 | 0 | 2 | 0 | 12 | 2.5 | 472 | 471 | 2 | 1.17 | 0.03 |
| CD9 | P21926 | 2 | 1 | 2 | 0 | 14 | 6.1 | 228 | 227 | 1 | 1.00 | 0.06 |
| COL10A1 | Q03692 | 2 | 2 | 0 | 0 | 11 | 1.6 | 680 | 662 | 1 | 1.00 | 0.02 |
| COL11A1 | P12107 | 2 | 0 | 2 | 0 | 11 | 0.6 | 1806 | 1052 | 2 | 1.36 | 0.01 |
| COL18A1 | P39060 | 2 | 1 | 1 | 0 | 14 | 0.8 | 1754 | 1731 | 1 | 1.00 | 0.01 |
| COL23A1 | Q86Y22 | 2 | 1 | 1 | 0 | 17 | 3.1 | 540 | 540 | 1 | 1.00 | 0.03 |
| COL24A1 | Q17RW2 | 2 | 2 | 0 | 0 | 13 | 0.8 | 1714 | 1679 | 1 | 1.00 | 0.01 |
| COL26A1 | Q96A83 | 2 | 3 | 0 | 0 | 11 | 2.5 | 441 | 421 | 1 | 1.00 | 0.02 |
| COL7A1 | Q02388 | 2 | 2 | 0 | 0 | 7 | 0.2 | 2944 | 2928 | 1 | 1.00 | 0.00 |

**Supplementary Table 3_2. Summary statistics of the identified proteins from Penghu 3.** The number of razor+unique peptides calculated from non-redundant peptides is shown as "Total", and the sum of those in other fractions does not equal "Total" due to the redundant peptides identified from multiple fractions. The non-redundant depth of the covered AA residues is also shown as "Mean" among the identified residues and as "Total", the sum of the depths of each residue divided by the protein length. The AA coverage (%) was calculated based on the AA sequence of the mature protein that does not contain a signal sequence or a propeptide, rather than the total AA sequence of the preproprotein.

| Gene name | Uniprot ID | R+U peptides | | | | AA | Coverage (%) | Length | | Depth | | |
|---|---|---|---|---|---|---|---|---|---|---|---|---|
| | | Total | Trypsin | Glu-C | Digestion-free | | | Total | Mature | Max | Mean | Total |
| COL1A1 | P02452 | 1034 | 1000 | 614 | 222 | 1012 | 69.1 | 1464 | 1057 | 71 | 11.73 | 8.11 |
| COL1A2 | P08123 | 807 | 863 | 517 | 76 | 955 | 69.9 | 1366 | 1040 | 78 | 9.65 | 6.74 |
| AHSG | P02765 | 40 | 34 | 34 | 2 | 109 | 29.7 | 367 | 309 | 14 | 3.37 | 1.00 |
| BGLAP | P02818 | 29 | 28 | 16 | 5 | 36 | 36.0 | 100 | 49 | 19 | 6.22 | 2.24 |
| CHAD | O15335 | 25 | 32 | 15 | 2 | 72 | 20.1 | 359 | 337 | 15 | 3.19 | 0.64 |
| SERPINF1 | P36955 | 24 | 33 | 9 | 3 | 75 | 17.9 | 418 | 399 | 9 | 2.73 | 0.49 |
| OMD | Q99983 | 22 | 21 | 27 | 2 | 60 | 14.3 | 421 | 401 | 12 | 3.77 | 0.54 |
| COL2A1 | P02458 | 21 | 19 | 16 | 2 | 111 | 7.5 | 1487 | 1102 | 9 | 1.76 | 0.13 |
| BGN | P21810 | 18 | 29 | 4 | 5 | 61 | 16.6 | 368 | 331 | 11 | 2.66 | 0.44 |
| F2 | P00734 | 18 | 32 | 3 | 0 | 79 | 12.7 | 622 | 579 | 7 | 1.95 | 0.25 |
| LUM | P51884 | 17 | 24 | 5 | 1 | 42 | 12.4 | 338 | 320 | 8 | 3.05 | 0.38 |
| COL5A2 | P05997 | 15 | 28 | 5 | 0 | 108 | 7.2 | 1499 | 1203 | 3 | 1.19 | 0.09 |
| COL5A1 | P20908 | 10 | 8 | 9 | 0 | 95 | 5.2 | 1838 | 1568 | 2 | 1.06 | 0.05 |
| COL9A1 | P20849 | 8 | 6 | 6 | 0 | 35 | 3.8 | 921 | 898 | 3 | 1.26 | 0.05 |
| ALB | P02768 | 7 | 12 | 2 | 0 | 42 | 6.9 | 609 | 585 | 2 | 1.52 | 0.11 |
| COL24A1 | Q17RW2 | 6 | 2 | 5 | 0 | 29 | 1.7 | 1714 | 1679 | 2 | 1.28 | 0.02 |
| COL3A1 | P02461 | 6 | 5 | 3 | 0 | 32 | 2.2 | 1466 | 1068 | 2 | 1.13 | 0.02 |

| | | | | | | | | | | | | |
|---|---|---|---|---|---|---|---|---|---|---|---|---|
| COL4A2 | P08572 | 5 | 4 | 2 | 0 | 30 | 1.8 | 1712 | 1529 | 1 | 1.00 | 0.02 |
| COL7A1 | Q02388 | 5 | 4 | 2 | 0 | 28 | 1.0 | 2944 | 2928 | 1 | 1.00 | 0.01 |
| VTN | P04004 | 5 | 9 | 1 | 0 | 29 | 6.1 | 478 | 459 | 2 | 1.66 | 0.10 |
| COL16A1 | Q07092 | 4 | 2 | 2 | 1 | 23 | 1.4 | 1604 | 1583 | 1 | 1.00 | 0.01 |
| COL22A1 | Q8NFW1 | 4 | 1 | 3 | 0 | 24 | 1.5 | 1626 | 1599 | 1 | 1.00 | 0.01 |
| COL17A1 | Q9UMD9 | 3 | 3 | 1 | 0 | 14 | 0.9 | 1497 | 1497 | 2 | 1.36 | 0.01 |
| COL28A1 | Q2UY09 | 3 | 1 | 3 | 0 | 23 | 2.0 | 1125 | 1102 | 1 | 1.00 | 0.02 |
| COL5A3 | P25940 | 3 | 1 | 3 | 0 | 12 | 0.7 | 1745 | 1716 | 2 | 1.17 | 0.01 |
| F10 | P00742 | 3 | 3 | 2 | 0 | 21 | 4.3 | 488 | 448 | 2 | 1.38 | 0.06 |
| KIF26A | Q9ULI4 | 3 | 0 | 3 | 0 | 10 | 0.5 | 1882 | 1882 | 3 | 2.20 | 0.01 |
| C1QC | P02747 | 2 | 4 | 1 | 0 | 18 | 7.3 | 245 | 217 | 1 | 1.00 | 0.07 |
| COL12A1 | Q99715 | 2 | 2 | 1 | 0 | 9 | 0.7 | 1366 | 1040 | 1 | 1.00 | 0.01 |
| COL21A1 | Q96P44 | 2 | 2 | 0 | 0 | 9 | 0.9 | 957 | 935 | 2 | 1.44 | 0.01 |
| COL26A1 | Q96A83 | 2 | 0 | 2 | 0 | 11 | 2.5 | 441 | 421 | 1 | 1.00 | 0.02 |
| COL4A3 | Q01955 | 2 | 1 | 1 | 0 | 11 | 0.7 | 1670 | 1642 | 1 | 1.00 | 0.01 |
| F9 | P00740 | 2 | 3 | 2 | 0 | 12 | 2.6 | 461 | 415 | 1 | 1.00 | 0.03 |
| OLFML3 | Q9NRN5 | 2 | 5 | 0 | 0 | 16 | 3.9 | 406 | 385 | 1 | 1.00 | 0.04 |
| SRPX | P78539 | 2 | 2 | 2 | 0 | 19 | 4.1 | 464 | 434 | 1 | 1.00 | 0.04 |
| THBS1 | P07996 | 2 | 6 | 0 | 0 | 11 | 0.9 | 1170 | 1152 | 1 | 1.00 | 0.01 |

### Supplementary Note 4:
### Authenticity of the recovered ancient proteomes

To further validate the authenticity of the recovered ancient proteomes of Penghu 2 and Penghu 3, the deamidation rates and peptide length distribution were analyzed. The results suggest an ancient origin of the identified proteome as described below.

#### Deamidation rates

Deamidation rates of asparagine (N) and glutamine (Q) residues increase with thermal age and thus are a proxy that positively correlates with the chronological age of the specimen[13]. The deamidation rates of enamel proteins from the early Pleistocene typically reach 100%[13-15], and those of bone proteins from the Middle to Late Pleistocene typically range between 50–100%[2,16-18]. Considering the chronological age of Penghu 2 (probably Late Pleistocene) and Penghu 3 (c. 45,000 cal BP), its deamidation rates of endogenous proteins would likely be higher than 50%. Therefore, the deamidation rates were calculated to evaluate the authenticity of identified proteins from Penghu 2 and Penghu 3.

Deamidation rates of the peptides originating from the identified 21 or 36 proteins were calculated for each .raw file for Penghu 2 and Penghu 3, respectively, using an R script rDeamidation[2] that was ported from a Python script[19]. The calculation was carried out for the distinct final MaxQuant search results (i.e., specific, semi-specific, and unspecific cleavage specificity).

The calculated deamidation rates of N and Q were greater than 88% and 67%, respectively, suggesting the ancient origin of the identified peptides from Penghu 2 (**Supplementary Table 4_1**) (**Supplementary Fig. 4_1**) and Penghu 3 (**Supplementary Table 4_2**) (**Supplementary Fig. 4_2**). The calculated deamidation rates of N and Q of proteins identified from experimental blanks were less than 25% and 33%, respectively, demonstrating the elevated deamidation rates of the endogenous proteins identified from Penghu 2 and Penghu 3 (**Supplementary Table 4_3**) (**Supplementary Fig. 4_3**).

#### Peptide length distribution

Ancient proteins degrade over time, and the length of peptides identified in digestion-free fractions tends to be shorter in more ancient specimens[14]. The distribution of peptide length has been investigated for the proteins identified in digestion-free fractions in this study.

The length distribution of peptides identified in the bone of Penghu 3 peaked at 15 amino acids (**Supplementary Fig. 4_4b**). Although that of Penghu 2 is ambiguous due to the relatively small number of identified peptides, most of the peptides consisted of ≤13 AA residues (**Supplementary Fig. 4_4a**). These results indicate that the identified proteins from Penghu 2 and Penghu 3 are degraded, supporting their ancient origin.

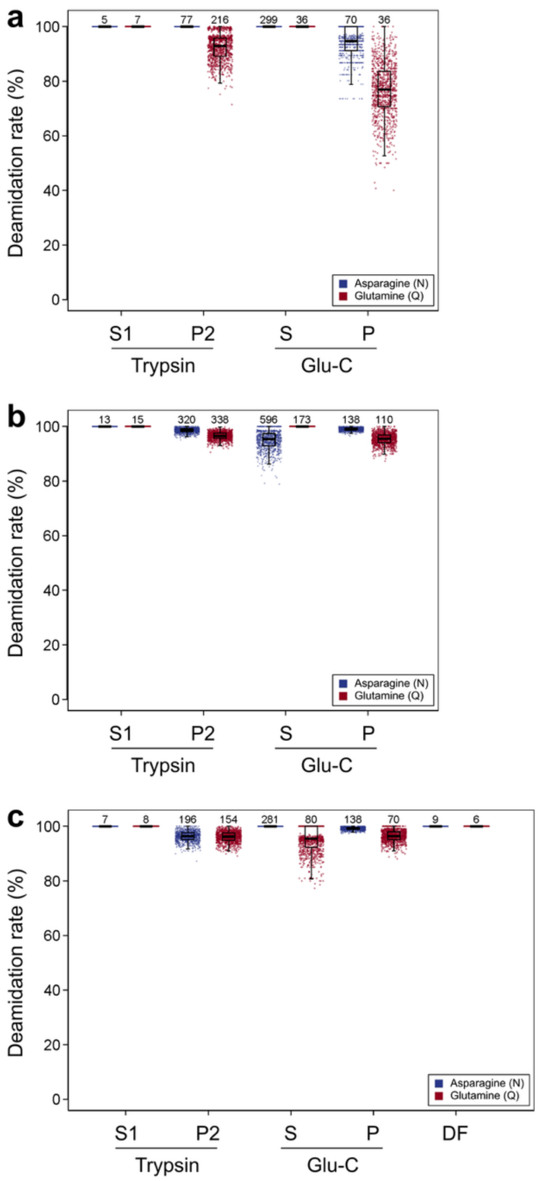


**Supplementary Fig. 4_1. Jitter and box plots of bootstrapped deamidation rates of each .raw files of Penghu 2.** Results from specific (a), semi-specific (b), and unspecific (c) searches are shown. The number of N or Q residues are shown above the jitters/boxes. “S” and “P” mean supernatant and pellet fractions, respectively.

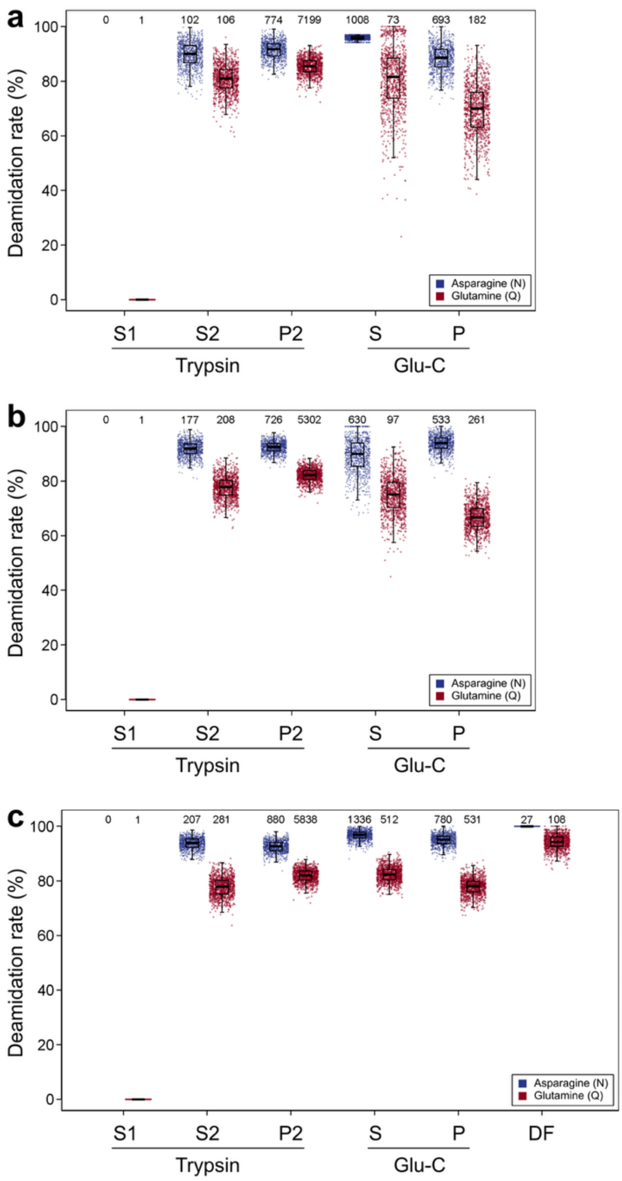


**Supplementary Fig. 4_2. Jitter and box plots of bootstrapped deamidation rates of each .raw files of Penghu 3.** Results from specific (a), semi-specific (b), and unspecific (c) searches are shown. The number of N or Q residues are shown above the jitters/boxes. “S” and “P” mean supernatant and pellet fractions, respectively.

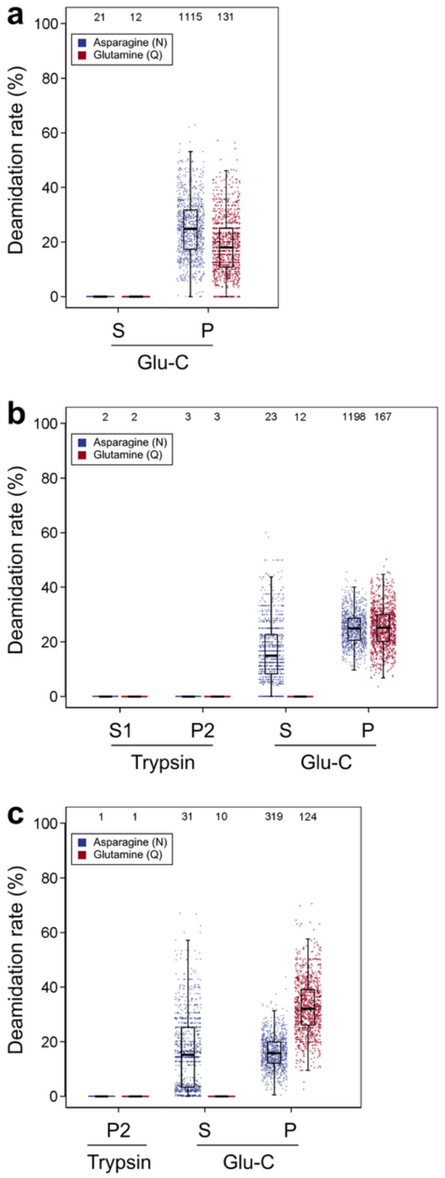


**Supplementary Fig. 4_3. Jitter and box plots of bootstrapped deamidation rates of each .raw files of experimental blanks.** Results from specific (a), semi-specific (b), and unspecific (c) searches are shown. The number of N or Q residues are shown above the jitters/boxes. “S” and “P” mean supernatant and pellet fractions, respectively.

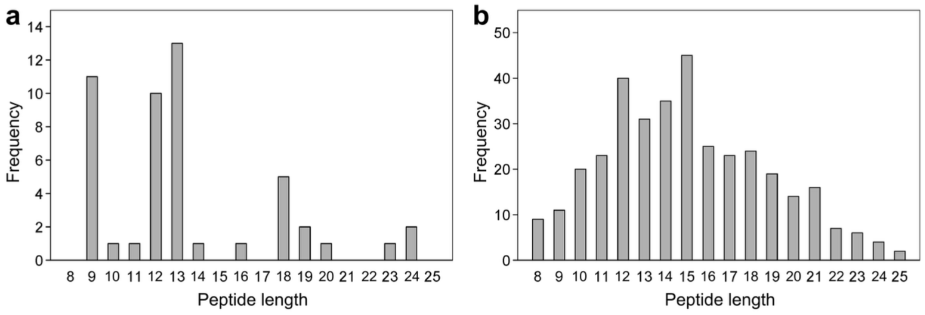

**Supplementary Fig. 4_4. Histogram showing the peptide length distribution.** The endogenous proteins identified from digestion-free fractions of Penghu 2 (a) and Penghu 3 (b) are shown.

**Supplementary Table 4_1. Summary of the calculated deamidation rates of the endogenous proteins of Penghu 2.**

| Search | Digestion | Fraction | Batch | N | | | | | Q | | | | |
|---|---|---|---|---|---|---|---|---|---|---|---|---|---|
| | | | | Mean | SD | 95% CI, lower | 95% CI, upper | n | Mean | SD | 95% CI, lower | 95% CI, upper | n |
| Specific | Trypsin | EDTA | 1 | 100.0 | 0.0 | 100.0 | 100.0 | 5 | 100.0 | 0.0 | 100.0 | 100.0 | 7 |
| | | Pellet | 2 | 100.0 | 0.0 | 100.0 | 100.0 | 77 | 92.4 | 4.8 | 82.3 | 99.7 | 216 |
| | Glu-C | EDTA | 4 | 100.0 | 0.0 | 100.0 | 100.0 | 299 | 100.0 | 0.0 | 100.0 | 100.0 | 36 |
| | | Pellet | 4 | 94.7 | 5.3 | 82.4 | 100.0 | 70 | 76.7 | 9.9 | 56.0 | 95.5 | 36 |
| Semi-specific | Trypsin | EDTA | 1 | 100.0 | 0.0 | 100.0 | 100.0 | 13 | 100.0 | 0.0 | 100.0 | 100.0 | 15 |
| | | Pellet | 2 | 98.5 | 0.8 | 96.6 | 100.0 | 320 | 96.5 | 1.4 | 93.4 | 98.9 | 338 |
| | Glu-C | EDTA | 4 | 94.9 | 3.6 | 87.0 | 100.0 | 596 | 100.0 | 0.0 | 100.0 | 100.0 | 173 |
| | | Pellet | 4 | 99.0 | 0.6 | 97.8 | 99.9 | 138 | 95.4 | 2.1 | 90.8 | 99.0 | 110 |
| Unspecific | Trypsin | EDTA | 1 | 100.0 | 0.0 | 100.0 | 100.0 | 7 | 100.0 | 0.0 | 100.0 | 100.0 | 8 |
| | | Pellet | 2 | 96.3 | 1.8 | 92.4 | 99.4 | 196 | 96.1 | 1.9 | 91.8 | 99.3 | 154 |
| | Glu-C | EDTA | 4 | 100.0 | 0.0 | 100.0 | 100.0 | 281 | 95.2 | 4.7 | 85.0 | 100.0 | 80 |
| | | Pellet | 4 | 99.2 | 0.5 | 98.0 | 100.0 | 138 | 96.4 | 2.1 | 91.9 | 100.0 | 70 |
| | Digestion-free | – | 3 | 100.0 | 0.0 | 100.0 | 100.0 | 9 | 100.0 | 0.0 | 100.0 | 100.0 | 6 |

**Supplementary Table 4_2. Summary of the calculated deamidation rates of the endogenous proteins of Penghu 3.**

| Search | Digestion | Fraction | Batch | N | | | | | Q | | | | |
|---|---|---|---|---|---|---|---|---|---|---|---|---|---|
| | | | | Mean | SD | 95% CI, lower | 95% CI, upper | n | Mean | SD | 95% CI, lower | 95% CI, upper | n |
| Specific | Trypsin | EDTA | 1 | – | – | – | – | 0 | 0.0 | 0.0 | 0.0 | 0.0 | 1 |
| | | EDTA | 2 | 89.6 | 4.6 | 79.7 | 97.4 | 102 | 80.8 | 5.2 | 70.5 | 90.6 | 106 |
| | | Pellet | 2 | 91.4 | 3.2 | 84.8 | 96.9 | 774 | 85.4 | 3.0 | 78.9 | 90.7 | 7199 |
| | Glu-C | EDTA | 4 | 95.9 | 0.8 | 94.2 | 97.0 | 1008 | 80.6 | 11.6 | 54.0 | 99.1 | 73 |
| | | Pellet | 4 | 88.4 | 4.8 | 78.4 | 97.2 | 693 | 69.4 | 9.3 | 50.0 | 86.1 | 182 |
| Semi-specific | Trypsin | EDTA | 1 | – | – | – | – | 0 | 0.0 | 0.0 | 0.0 | 0.0 | 1 |
| | | EDTA | 2 | 91.8 | 2.8 | 86.2 | 96.7 | 177 | 77.6 | 4.1 | 69.6 | 85.3 | 208 |
| | | Pellet | 2 | 92.4 | 2.1 | 88.2 | 96.3 | 726 | 82.1 | 2.5 | 77.0 | 86.8 | 5302 |
| | Glu-C | EDTA | 4 | 89.3 | 6.2 | 76.1 | 100.0 | 630 | 74.8 | 6.8 | 61.5 | 88.0 | 97 |
| | | Pellet | 4 | 93.7 | 2.8 | 87.8 | 98.6 | 533 | 66.8 | 4.8 | 57.6 | 76.4 | 261 |
| Unspecific | Trypsin | EDTA | 1 | – | – | – | – | 0 | 0.0 | 0.0 | 0.0 | 0.0 | 1 |
| | | EDTA | 2 | 93.7 | 2.1 | 89.1 | 97.4 | 207 | 77.7 | 3.6 | 70.9 | 84.6 | 281 |
| | | Pellet | 2 | 92.6 | 2.1 | 88.2 | 96.3 | 880 | 81.9 | 2.4 | 77.1 | 86.2 | 5838 |
| | Glu-C | EDTA | 4 | 96.7 | 1.7 | 93.0 | 99.5 | 1336 | 82.4 | 2.7 | 77.1 | 87.7 | 512 |

| | | | | | | | | | | | | | |
|---|---|---|---|---|---|---|---|---|---|---|---|---|---|
| | | Pellet | 4 | 94.9 | 2.1 | 90.3 | 98.3 | 780 | 77.9 | 3.0 | 71.4 | 83.6 | 531 |
| | Digestion-free | – | 3 | 100.0 | 0.0 | 100.0 | 100.0 | 27 | 94.2 | 2.7 | 88.6 | 98.7 | 108 |

**Supplementary Table 4_3. Summary of the calculated deamidation rates of the exogenous proteins of experimental blanks.**

| **Search** | **Digestion** | **Fraction** | **Batch** | **N** | | | | | **Q** | | | | |
|---|---|---|---|---|---|---|---|---|---|---|---|---|---|
| | | | | Mean | SD | 95% CI, lower | 95% CI, upper | n | Mean | SD | 95% CI, lower | 95% CI, upper | n |
| Specific | Glu-C | EDTA | 4 | 0.0 | 0.0 | 0.0 | 0.0 | 21 | 0.0 | 0.0 | 0.0 | 0.0 | 12 |
| | | Pellet | 4 | 25.1 | 10.6 | 6.4 | 47.1 | 1115 | 18.7 | 10.4 | 0.0 | 41.7 | 131 |
| Semi-specific | Trypsin | EDTA | 1 | 0.0 | 0.0 | 0.0 | 0.0 | 2 | 0.0 | 0.0 | 0.0 | 0.0 | 2 |
| | | Pellet | 2 | 0.0 | 0.0 | 0.0 | 0.0 | 3 | 0.0 | 0.0 | 0.0 | 0.0 | 3 |
| | Glu-C | EDTA | 4 | 16.2 | 11.1 | 0.0 | 40.0 | 23 | 0.0 | 0.0 | 0.0 | 0.0 | 12 |
| | | Pellet | 4 | 25.0 | 5.9 | 14.0 | 37.3 | 1198 | 25.2 | 7.5 | 11.1 | 40.8 | 167 |
| Unspecific | Trypsin | Pellet | 2 | 0.0 | 0.0 | 0.0 | 0.0 | 1 | 0.0 | 0.0 | 0.0 | 0.0 | 1 |
| | Glu-C | EDTA | 4 | 16.2 | 13.3 | 0.0 | 45.4 | 31 | 0.0 | 0.0 | 0.0 | 0.0 | 10 |
| | | Pellet | 4 | 16.1 | 5.8 | 5.3 | 28.0 | 319 | 32.8 | 9.9 | 14.7 | 53.2 | 124 |

### Supplementary Note 5:
### Evaluation of variants

Amino acid residues that have variations among Penghu 2, Penghu 3, Denisova 3, Neanderthals, and modern humans are especially important to investigate the phylogenetic relationship and evolutionary history of Penghu 2 and Penghu 3. Therefore, such positions were evaluated in detail below.

First, covered positions of proteins identified in Penghu 2 or Penghu 3 with inconsistencies of corresponding AA residues among any of the reference human proteome (UP000005640), a Denisovan proteome translated from the ancient genome of Denisova 3[20], the partial proteome of Xiahe 1[16], Xiahe 2[17], Penghu 1[2], and Harbin cranium[18], the reference proteome of three Neanderthal individuals with high-coverage ancient genomes (Denisova 5[21], Vindija 33.19[22], and Chagyrskaya 8[23]), and *H. sapiens* (Ust'-Ishim[24] and Loschbour[25]) were listed. The corresponding residues of apes (chimpanzees, gorillas, orangutans, and gibbons) were also checked to estimate the ancestral allele. Then, if the position is registered as a SNP, the allele frequency of the corresponding positions was checked in a modern human SNP database[26], and Neanderthal ancient genomes to investigate the possible variation of the position. This analysis allows us to estimate whether the variation is truly unique to specific taxa or whether it just happens to appear informative in randomly sequenced variations that exist in both humans and archaic hominins. When derived variants that appeared specific to Denisovans or ancient hominins (i.e., Denisovans and Neanderthals) were observed in SNP polymorphisms in *H. sapiens*, we judged that the polymorphism was likely to have been maintained among hominins and therefore did not subject them to further investigation. This applied to AHSG M248T, COL1A2 P549A, and COL9A1 Q621R (**Supplementary Table 5_1**). When the derived SAP was inferred to be specific to Denisovans or to the Penghu hominin, we carried out further investigations. For such variants, the proteomic authenticity of peptides or PSMs covering such positions was evaluated for Penghu 2 and Penghu 3 (**Supplementary Table 5_2**). If the peptide has an exogenous origin, the variation is no longer valid.

Peptides or PSMs covering such sites were evaluated based on the following criteria:

1. Depth: This is the number of peptides that cover the position. If the variant position was covered by ≥2 different peptides with support from tandem mass spectra, the variant can be regarded as confident.
2. Deamidation rate: Considering the overall mean deamidation rate of peptides from Penghu 2 and Penghu 3 was ≥88% in N and ≥67% in Q, most N and Q residues should be deamidated in ancient endogenous peptides. If the deamidation rate of N or Q residues in the PSMs covering the subject residue was too low (e.g., <50%), the corresponding peptides likely originated from modern contaminant proteins. To calculate the deamidation rates, PSMs covering the subject variant and including Q and/or N residues were evaluated [19]. The number of corresponding PSMs, the count of covered N and Q sites, and each deamidation status (deamidated or not) were retrieved from the evidence.txt file of the final MaxQuant outputs. The mean deamidation rates were calculated for all PSMs containing N and/or Q that cover the subject variant position. The calculated deamidation rates may be

the subject of various biases that can be caused when (i) the exact same molecules were identified in different MaxQuant searches (computational multiplicity) and (ii) the same molecules with a high concentration were selected for collision dissociation and analyzed in MS2 multiple times (measurement multiplicity). However, these biases have minimal effect on the results. Results from the exact same molecules were counted once for the calculation, which eliminates the computational multiplicity. The dynamic exclusion of mass spectrometry acquisition mostly suppresses the bias caused by measurement multiplicity. It is also possible to incorporate the information on ion intensity to achieve semi-quantitative information on peptide concentration[19], but we decided not to incorporate the intensity information. The deamidation of the obtained peptides is more than 50% (suggesting it is endogenous to Penghu hominins) or relatively low (i.e., <40%, suggesting modern contaminant). A simple calculation that does not consider peptide/PSM intensity was therefore sufficient for this study.

3. Codon degeneracy: For the observed variants of amino acid residues, it is highly unlikely that the DNA codons encoding the corresponding amino acid before and after the mutation differ by more than two bases. Variations in the amino acid sequence occur because of variations in the DNA sequence, and mutations in the DNA sequence are rare events. Therefore, if the subject amino acid variant requires more than two bases of DNA mutation to occur, the result is likely an artifact.
4. BLAST: This measure evaluates the subject sequence from the viewpoint of wider phylogenetic comparison. If the variant sequence matches other proteins from other taxa, it is possible that the peptide was of exogenous origin. If the sequence of the subject peptide matches the sequence of typical contaminants, such as hair keratin and bovine albumin, in the BLAST search, the peptide likely originated from contaminant proteins.
5. Collagen-specific features: If the subject variation is in collagenous proteins, the corresponding peptides would follow the typical two features of collagens. First, most of the collagen sequence consists of a G-X-Y repeat, where the first glycine appears in every three amino acid residues and is important for the tight wrapping of the triple-helix structure[27]. Therefore, if the variation appears in the relatively conserved G position of the G-X-Y repeat, we need to be cautious about the source of the subject peptide. Second, the Y position of the G-X-Y repeat is often hydroxylated proline, which facilitates the stability of the triple-helix structure of collagen by providing hydrogen bonds[28,29]. Therefore, if the proline in the peptide that covers the subject position was hydroxylated, it is more likely that the peptide originated from collagen, not from common laboratory contaminants such as albumin or keratin.

Based on these criteria, two variants were confidently identified as described below. We decided conservatively, and if the variant was not clearly supported, it was declined.

**The 266th position of COL1A2 (unlikely variant)**

The 266th position of COL1A2 is E in humans, archaic hominins, and apes. Two peptides supporting the E residue were sequenced from Penghu 3, as well as one peptide supporting the derived G residue (**Supplementary Table 5_2**). The SNP corresponding to the G variant does not exist in the database.

The G residue for this position is considered unlikely. Only one supporting peptide was obtained for this derived variant, and its deamidation rate for N was relatively low (i.e., 33.3%) for Pleistocene proteins (**Supplementary Table 5_2**). The amino acid change from E to G can occur with a single base mutation in the DNA sequence. The BLAST search showed that the peptide sequence containing this derived variant is present in *H. sapiens*. The G-X-Y repeat was not hindered by the change from E to G, and some proline residues were hydroxylated in PSMs with either supported E or supported G. Therefore, based on the low depth and low deamidation rates of peptides with derived G variant, we assigned only ancestral E for this position in Penghu 3.

**The 578th position of COL1A2 (Penghu 3-specific)**

The 578th position of COL1A2 is E in hominins and apes, and peptides supporting both E and Q were obtained from Penghu 3 (**Supplementary Table 5_2**). No SNP corresponding to the derived Q residue is registered.

Although only E residue was obtained from Penghu 2 (**Supplementary Fig. 5_1**), peptides are well-supported for both E and Q at this position in Penghu 3, as shown below (**Supplementary Fig. 5_2, Supplementary Table 5_2**). The depths of E and Q are both 13 and 5 for peptides supported by MS2 spectra, respectively, supporting both residues. The deamidation rate of Q was 75% for PSMs with the Q variant, but this figure appears imprecise, as the deamidated Q cannot be distinguished from E. The amino acid change from E to V can occur with a single base mutation in the DNA sequence. BLAST search showed that the identified peptide sequences with Q cannot be seen in other taxa. The G-X-Y repeat was not hindered by the change from E to Q, and some proline residues were hydroxylated in PSMs with either E or Q.

We concluded that this Penghu 3-specific Q variant exists as a heterozygous position with the shared variant of E. Since several criteria equally support peptides with either E or Q, we cannot dismiss either of these residues. For the phylogenetic analyses, we conservatively chose E for this position because of its greater depth. Only the E variant was identified from Penghu 2, but its depth of supporting peptides was one.

**The 996th position of COL1A2 (Denisovan-specific)**

The 996th position of COL1A2 is K in all five Denisovans and R in other hominins and apes (**Supplementary Table 5_2**). No SNP causing this variation is registered in the gnomAD database, suggesting the frequency of the K variant is almost zero or zero in modern human populations. Sequenced Neanderthal individuals have R at this position.

The K residue has abundant support in Penghu hominins, as described below. The depth of K was 4 and 10 for non-redundant peptides with MS2 spectral support at the subject position of Penghu 2 (**Supplementary Fig. 5_1**) and Penghu 3 (**Supplementary Fig. 5_2**), respectively. The N and Q deamidation rates were 100% and ≥86.7%, supporting their ancient endogenous origin. The amino acid change from R to K can occur with a single base mutation in the DNA sequence. BLAST search showed that the peptide sequence with K at the subject position cannot be seen in other taxa if the part towards the C-terminus end is included. The G-X-Y repeat was not hindered by the change from R to K, and some proline residues were hydroxylated in the subject PSMs.

We accepted this K variant in Penghu 2 and Penghu 3 as a Denisovan marker. This K variant is supported by the ancient genomic sequence of Denisova 3 and the ancient protein sequence of Xiahe 1, Xiahe 2, Penghu 1, and the Harbin cranium. In contrast, humans and Neanderthals have R at this position. The deamidation rates were almost 100% for N and Q residues.

**The 611th position of COL2A1 (Penghu 2- and Penghu 3-specific)**

The 611th position of COL2A1 is N in hominins and apes, and peptides supporting S were obtained from Penghu 2 and Penghu 3 (**Supplementary Table 5_2**). No SNP corresponding to this position is registered in the gnomAD database. Among Denisovans, this position was sequenced only in Denisova 3 and the Harbin cranium, and both of them have N. Three Neanderthal individuals have N at this position.

Peptides supporting S at this position were obtained, but the depth was 1 for both Penghu 2 (**Supplementary Fig. 5_3**) and Penghu 3 (**Supplementary Fig. 5_4**). The deamidation rate cannot be calculated because there was no N or Q residue covered by the subject PSMs. The amino acid change from N to S can occur with a single base mutation in the DNA sequence. BLAST search showed that the identified peptide sequences with S cannot be seen in other taxa. The G-X-Y repeat was not hindered by the change from N to S, and some proline residues were hydroxylated in PSMs.

We concluded that this is a potential Penghu 2- and Penghu 3-specific S variant, but the low depth limits a definitive conclusion. For the phylogenetic analyses, although we retained this S variant for this position, the tree topology was the same if this position was removed along with other positions with MS2 support in ≤1 depth (**Supplementary Note 6**).

**The 1211th position of COL5A2 (Penghu 2-specific)**

The 1211th position of COL5A2 is E in hominins and apes, and a peptide supporting V was obtained from Penghu 2 (**Supplementary Table 5_2**). A SNP corresponding to the derived V residue (rs752139160) can be seen in only one individual from South Asia at the worldwide level (<0.01%, n = 1,608,970) in the gnomAD database. For Denisovans, although Denisova 3 and the Harbin cranium have E at this position, Penghu 1 showed heterozygosity of E and V. Three Neanderthal individuals have E at this position.

A peptide supporting V at this position was obtained in Penghu 2, but the depth was 1 (**Supplementary Fig. 5_5**). The deamidation rate cannot be calculated because there was no N or Q residue covered by the subject PSMs. The amino acid change from E to V can occur with a single base mutation in the DNA sequence. BLAST search showed that identified peptide sequences with V cannot be seen in other taxa. The G-X-Y repeat was not hindered by the change from E to V, and some proline residues were hydroxylated in PSMs.

We concluded that this is a potential Penghu 2-specific V variant, but the low depth limits a definitive conclusion. One interesting point was that this V variant was also independently identified in Penghu 1 mandible[2]. For the phylogenetic analyses, although we retained this V variant for this position, the tree topology was the same if this position was removed along with other positions with MS2 support in ≤1 depth (**Supplementary Note 6**).

**Summary of the identified variants**

Among the 1856 and 3283 amino acid residues from 21 and 36 proteins recovered in Penghu 2 and Penghu 3, respectively, four residues from three proteins were confidently or potentially related to Denisovan-specific variation (**Supplementary Table 5_2**).

One derived variant specific to Denisovans was identified in COL1A2 of Penghu 2 (**Supplementary Fig. 5_1**) and Penghu 3 (**Supplementary Fig. 5_2**). The derived variant COL1A2 R996K was observed only in Penghu 1, Xiahe 1, Xiahe 2, the Harbin cranium, and Denisova 3 among the sequenced hominin individuals. This Denisovan-type variant was identified at peptide depths of 4× and 10× from Penghu 2 and Penghu 3, respectively, indicating that they are molecularly identified Denisovans.

Three derived variants, observed only in specific Denisovan individuals, were potentially identified in COL1A2, COL2A1, and COL5A2. The COL1A2 E578Q variant was observed in Penghu 3 in a heterozygous state (**Supplementary Fig. 5_4**). The COL2A1 N611S variant was observed in Penghu 2 (**Supplementary Fig. 5_5**) and Penghu 3 (**Supplementary Fig. 5_6**). The variant COL5A2 E1211V was seen in Penghu 2, as well as the Penghu 1 mandible[2]. These variants were not identified in other hominin individuals sequenced to date. However, the depth of peptides with MS2 support for the positions of COL2A1 and COL5A2 was only 1, and thus the calls for these derived variants are not confident. A derived variant, COL2A1 E582G, reported in Xiahe 1[16], was not covered in Penghu 2 and Penghu 3.

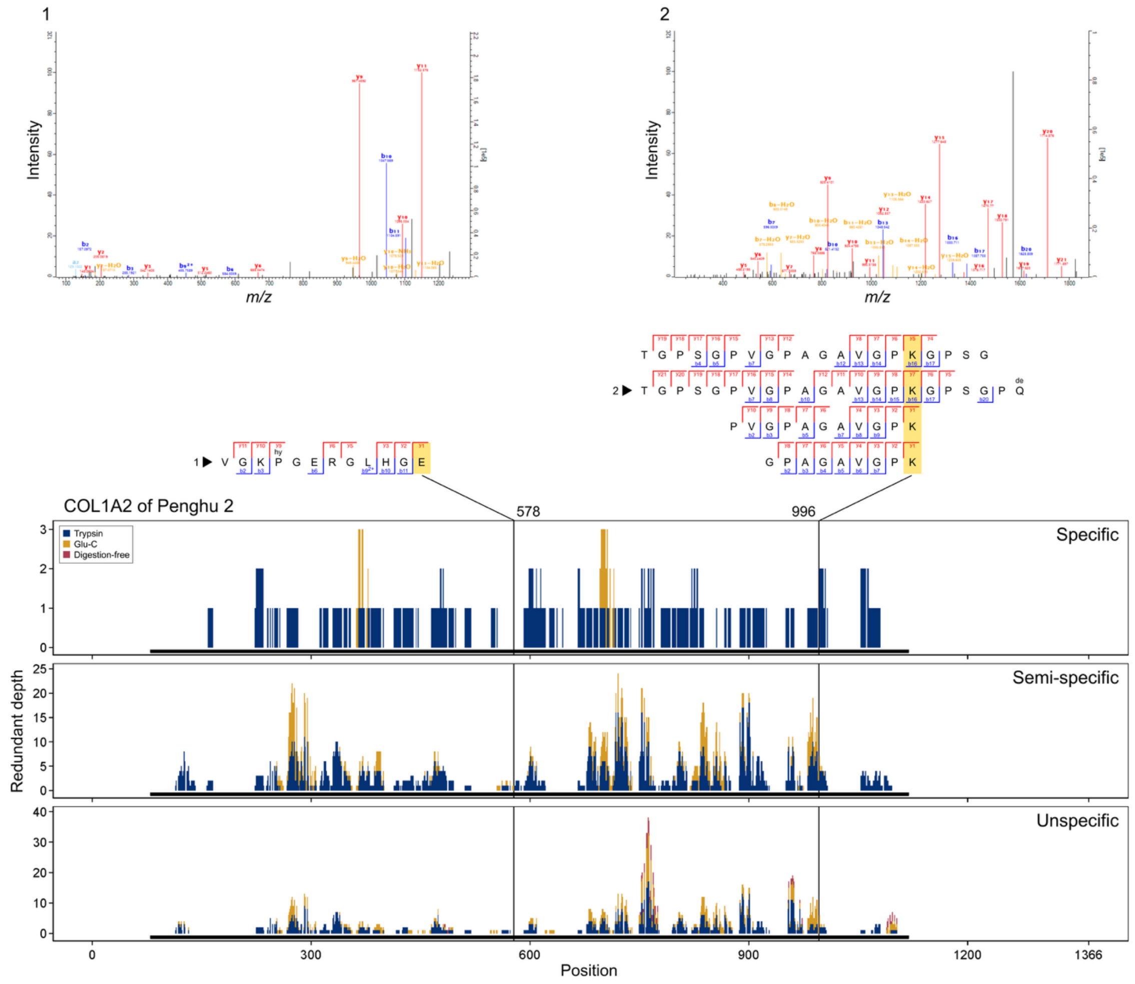


**Supplementary Fig. 5_1. Details of the variant in COL1A2 of Penghu 2.** The depth of supporting peptides across the entire protein sequence was shown for each search and extraction method. The regions of mature protein without signal sequence and propeptide are shown with black solid bars under the histograms. Supporting peptides were aligned, and the subject position was highlighted in yellow. The corresponding PSMs with the displayed MS2 spectra were marked with a triangle and a number.

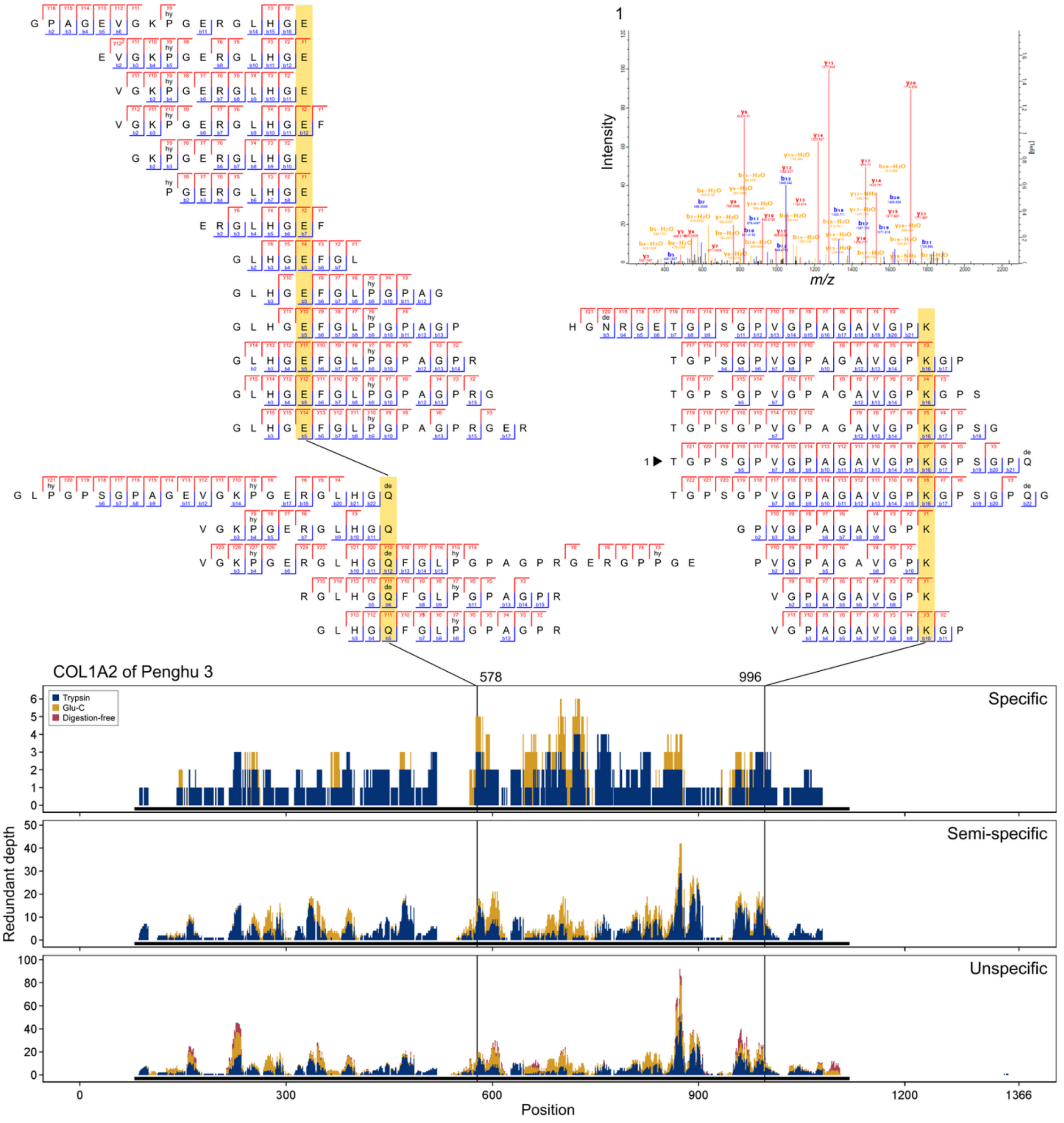


**Supplementary Fig. 5_2. Details of the variant in COL1A2 of Penghu 3.** The depth of supporting peptides across the entire protein sequence was shown for each search and extraction method. The regions of mature protein without signal sequence and propeptide are shown with black solid bars under the histograms. Supporting peptides were aligned, and the subject position was highlighted in yellow. The corresponding PSMs with the displayed MS2 spectra were marked with a triangle and a number.

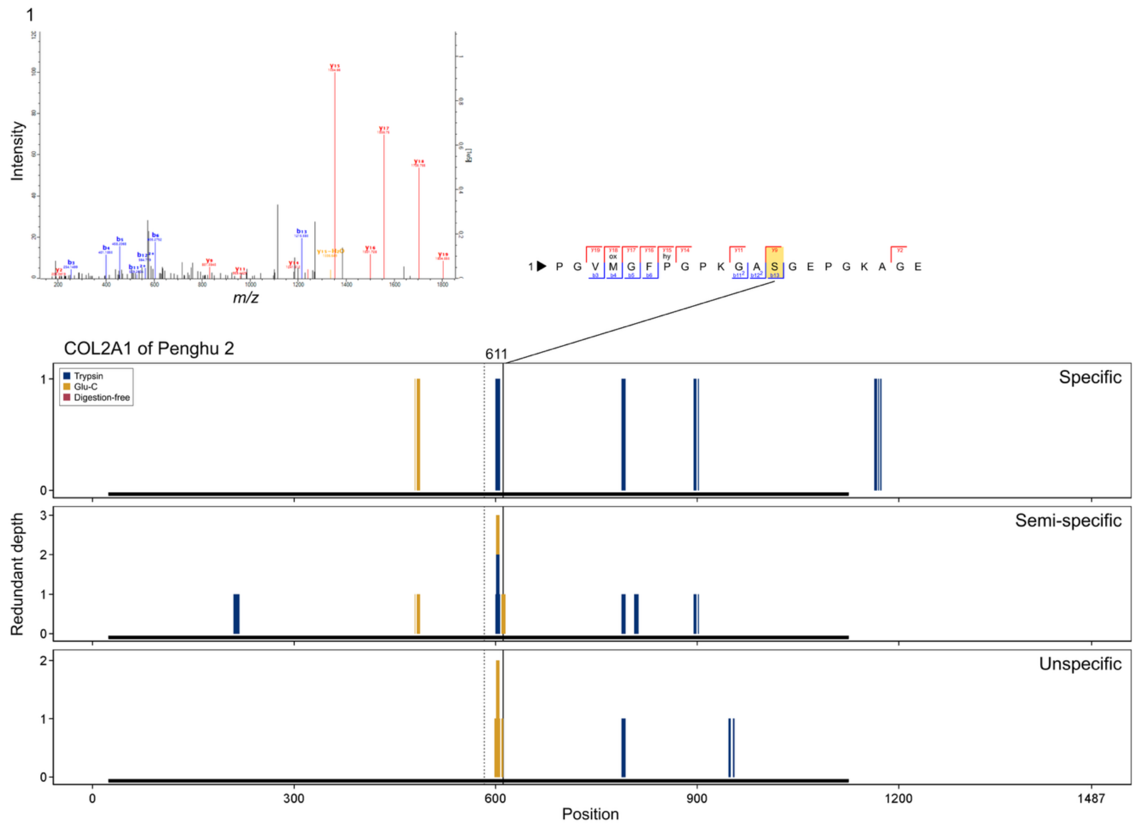


**Supplementary Fig. 5_3. Details of the variant in COL2A1 of Penghu 2.** The depth of supporting peptides across the entire protein sequence was shown for each search and extraction method. The regions of mature protein without signal sequence and propeptide are shown with black solid bars under the histograms. Supporting peptides were aligned, and the subject position was highlighted in yellow. The corresponding PSMs with the displayed MS2 spectra were marked with a triangle and a number.

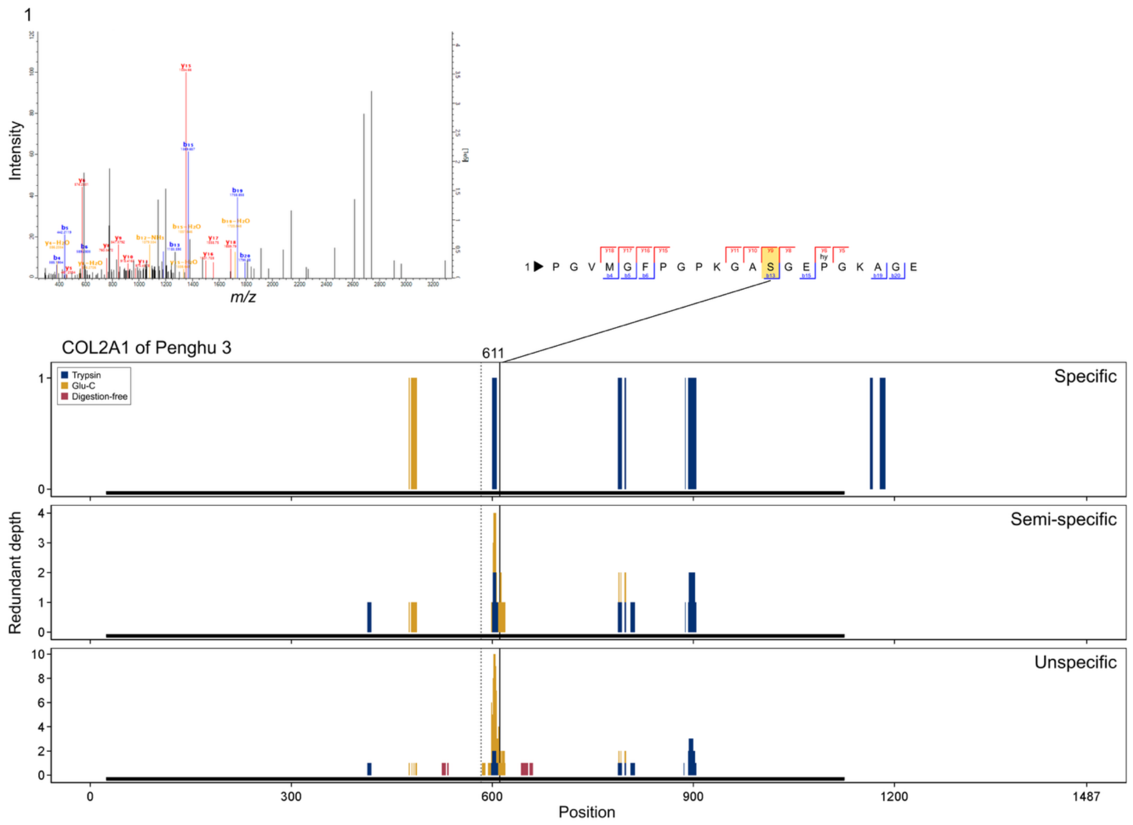


**Supplementary Fig. 5_4. Details of the variant in COL2A1 of Penghu 3.** The depth of supporting peptides across the entire protein sequence was shown for each search and extraction method. The regions of mature protein without signal sequence and propeptide are shown with black solid bars under the histograms. Supporting peptides were aligned, and the subject position was highlighted in yellow. The corresponding PSMs with the displayed MS2 spectra were marked with a triangle and a number.

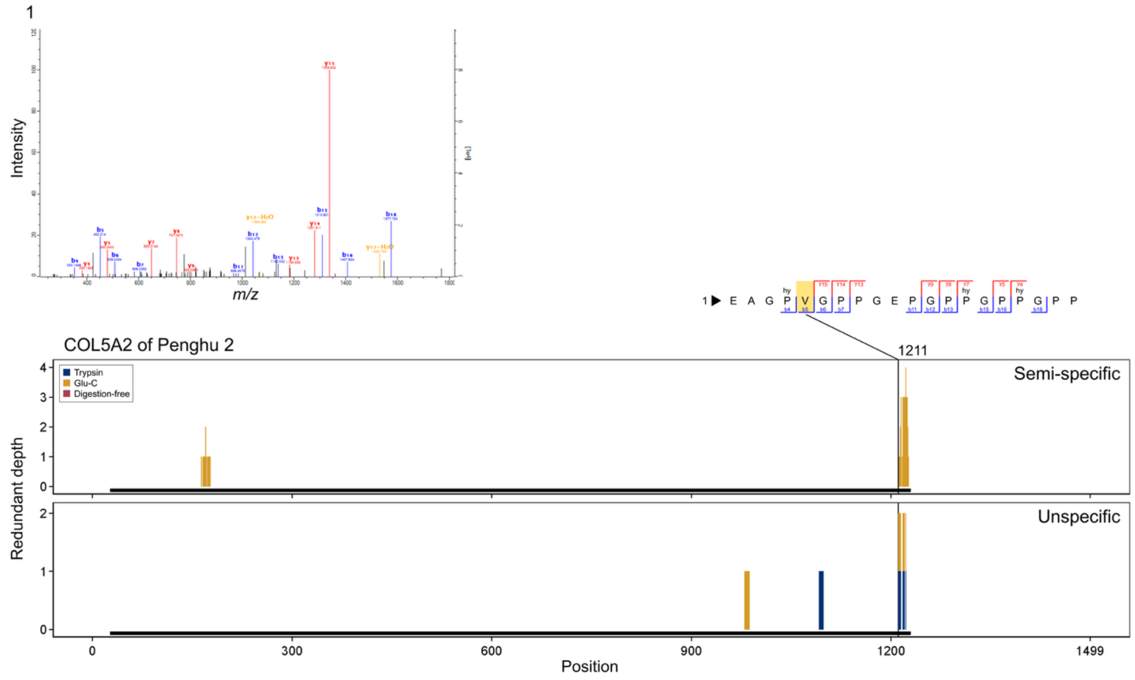


**Supplementary Fig. 5_5**. Details of the variant in COL5A2 of Penghu 2. The depth of supporting peptides across the entire protein sequence was shown for each search and extraction method. The regions of mature protein without signal sequence and propeptide are shown with black solid bars under the histograms. Supporting peptides were aligned, and the subject position was highlighted in yellow. The corresponding PSMs with the displayed MS2 spectra were marked with a triangle and a number.

**Supplementary Table 5_1. Comparison of AA residues at the positions of phylogenetically informative variant candidates.** Two identical or different AA residues that are shown with a hyphen mean homozygosity or heterozygosity, respectively. Numbers shown with some AA residues mean the depth of the non-redundant peptides that cover the subject position in the Penghu 2 or Penghu 3. The sequence around the variant is also shown in bold at the subject AA residue.

| # | Gene name | Uniprot ID | Position | GrCh38 Position | GrCh37 Position | SNP ID | SNP that can cause this change | SNP frequency |
|---|---|---|---|---|---|---|---|---|
| 1 | AHSG | P02765 | 248 | Chr3:186619924 | Chr3:186337713 | rs4917 | T/A, C | 0.66 |
| 2 | COL1A2 | P08123 | 266 | Chr7:94409326 | Chr7:94038638 | - | (A/C) | - |
| 3 | COL1A2 | P08123 | 549 | Chr7:94413927 | Chr7:94043239 | rs42524 | G to C (GCT -> A) | 0.76 |
| 4 | COL1A2 | P08123 | 578 | Chr7:94415238 | Chr7:94044550 | - | (G/C) | - |
| 5 | COL1A2 | P08123 | 996 | Chr7:94426041 | Chr7:94055353 | rs1792270869 | G to A | 0.00000062 |
| 6 | COL2A1 | P02458 | 611 | Chr12:47984996 | Chr12:48378779 | rs142492439 | T/C | 0.00001426 |
| 7 | COL5A2 | P05997 | 1211 | Chr2:189041587 | Chr2:189906313 | rs752139160 | A/G | 0.000000622 |
| 8 | COL9A1 | P20849 | 621 | Chr6:70252130 | Chr6:70961833 | rs1135056 | T to C | 0.4 |

| # | n | Decision | Notes | Sequence | Penghu 2 | Penghu 3 |
|---|---|---|---|---|---|---|
| 1 | 1611398 | Polymorphism | 1070970/1611398 | VTC**T**VFQ | T1 | T2 |
| 2 | No SNPs reported in *H. sapiens* | Unique | No SNPs reported in H. sapiens | PKG**E**IGA | – | E2G1 |
| 3 | 1612780 | Polymorphism | C/G (G/C) 1231835/1612780 | QGP**A**GPP | A1 | A1 |
| 4 | No SNPs reported in *H. sapiens* | Unique | No SNPs reported in H. sapiens | LHG**Q**FGL | E1 | E13Q5 |
| 5 | 1614086 | Unique | 1/1614086=6.195e-7 (1 in African/African American) | VGP**K**GPS | K4 | K10 |
| 6 | 1613176 | Unique | 23/1613176=0.00001426 (18 in non-Finnish European) | KGA**S**GEP | S1 | S1 |
| 7 | 1608970 | Unique | 1/1608970 (1 in South Asian) | AGP**V**GPP | V1 | – |
| 8 | 1613484 | Polymorphism | 640959/1613484 | RGP**Q**GLP | Q1 | Q1 |

| # | Denisova 3 | Penghu 1 | Xiahe 1 | Xiahe 2 | Harbin | Human SNP | Denisova 5 | Vindija 33.19 | Chagyrskaya 8 | Ust'-Ishim | Loschbour | Apes |
|---|---|---|---|---|---|---|---|---|---|---|---|---|
| 1 | T/T | – | – | – | – | M/T | T/T | T/T | T/T | T/T | M/T | T |
| 2 | E/E | E | E | E | E | E | E/E | E/E | E/E | E/E | E/E | E |
| 3 | A/A | A | A | A | A | A/P | A/A | A/A | A/A | A/A | A/A | A |
| 4 | E/E | E | E | E | E | E | E/E | E/E | E/E | E/E | E/E | E |
| 5 | K/K | K | K | K | K | R | R/R | R/R | R/R | R/R | R/R | R |
| 6 | N/N | – | – | – | N | N | N/N | N/N | N/N | N/N | N/N | N |
| 7 | E/E | V/E | – | – | E | E | E/E | E/E | E/E | E/E | E/E | E |
| 8 | R/R | Q | Q | Q | – | Q/R | Q/R | Q/Q | Q/Q | Q/R | Q/Q | Q |

**Supplementary Table 5_2. Summary of the proteomic authenticity of peptides and PSMs covering the positions of phylogenetically informative variant candidates.** The “Supported peptides” and “Supported PSMs” are the sum of unique sequences or molecules that only contain peptides or PSMs with MS2 spectral support at the subject position. Deami: Deamidation of N or Q residues. n_total: total number of N or Q counted for all PSMs that cover the subject variant position. The sequence around the variant is also shown in bold at the subject AA residue.

| # | Gene name | Uniprot ID | Position | Residue | Derived | Specimen | Depth | | | |
|---|---|---|---|---|---|---|---|---|---|---|
| | | | | | | | Peptide | PSM | Supported peptide | Supported PSM |
| 1 | COL1A2 | P08123 | 266 | E | No | Penghu 2 | 13 | 56 | 0 | 0 |
| 2 | | | | | | Penghu 3 | 15 | 54 | 2 | 3 |
| 3 | COL1A2 | P08123 | 266 | G | Yes | Penghu 2 | 1 | 2 | 0 | 0 |
| 4 | | | | | | Penghu 3 | 1 | 3 | 1 | 1 |
| 5 | COL1A2 | P08123 | 578 | E | No | Penghu 2 | 2 | 4 | 1 | 3 |
| 6 | | | | | | Penghu 3 | 22 | 93 | 13 | 43 |
| 7 | COL1A2 | P08123 | 578 | Q | Yes | Penghu 2 | 0 | 0 | 0 | 0 |
| 8 | | | | | | Penghu 3 | 6 | 16 | 5 | 10 |
| 9 | COL1A2 | P08123 | 996 | R | No | Penghu 2 | 0 | 0 | 0 | 0 |
| 10 | | | | | | Penghu 3 | 2 | 3 | 0 | 1 |
| 11 | COL1A2 | P08123 | 996 | K | Yes | Penghu 2 | 18 | 51 | 4 | 4 |
| 12 | | | | | | Penghu 3 | 27 | 97 | 10 | 19 |
| 13 | COL2A1 | P02458 | 611 | S | Yes | Penghu 2 | 2 | 8 | 1 | 3 |
| 14 | | | | | | Penghu 3 | 5 | 29 | 1 | 1 |
| 15 | COL5A2 | P05997 | 1211 | V | Yes | Penghu 2 | 6 | 7 | 1 | 2 |
| 16 | | | | | | Penghu 3 | 2 | 5 | 0 | 0 |

| # | Deami N | | Deami Q | | Consistent codon degeneracy | BLAST hit |
|---|---|---|---|---|---|---|
| | Percent | n_total | Percent | n_total | | |
| 1 | 100.0 | 47 | – | 0 | – | Theria |
| 2 | 91.2 | 34 | – | 0 | | |
| 3 | 100.0 | 2 | – | 0 | Yes | *Homo sapiens* |
| 4 | 33.3 | 3 | – | 0 | | |
| 5 | – | 0 | – | 0 | – | *Homo sapiens*, Klebsiella pneumoniae |
| 6 | – | 0 | – | 0 | | |
| 7 | – | – | – | – | Yes | No hit |
| 8 | – | 0 | 75.0 | 16 | | |
| 9 | – | – | – | – | – | *Homo sapiens* |
| 10 | – | 0 | – | 0 | | |
| 11 | 100.0 | 2 | 100.0 | 3 | Yes | No hit |
| 12 | 100.0 | 34 | 86.7 | 15 | | |
| 13 | – | 0 | – | 0 | Yes | No hit |
| 14 | – | 0 | – | 0 | | |
| 15 | – | 0 | – | 0 | Yes | No hit |
| 16 | – | 0 | – | 0 | | |

| # | G-X-Y repeat | Hydroxyproline | Confidence | Note | Sequence |
|---|---|---|---|---|---|
| 1 | Yes | Yes | Moderate | | PKG**E**IGA |
| 2 | | Yes | High | | |
| 3 | Yes | No | Low | | PKG**G**IGA |
| 4 | | Yes | Low | | |
| 5 | Yes | Yes | Moderate | | LHG**E**FGL |
| 6 | | Yes | High | | |
| 7 | Yes | – | No hit | | LHG**Q**FGL |
| 8 | | Yes | High | | |
| 9 | Yes | – | No hit | | VGP**R**GPS |
| 10 | | Yes | Low | | |
| 11 | Yes | Yes | High | | VGP**K**GPS |
| 12 | | Yes | High | | |
| 13 | Yes | Yes | Moderate | | KGA**S**GEP |
| 14 | | Yes | Moderate | | |
| 15 | Yes | Yes | Moderate | Also identified in Penghu 1 | AGP**V**GPP |
| 16 | | Yes | Low | | |

**Supplementary Note 6:**
**Phylogenetic analysis**

Phylogenetic trees were constructed based on the sequences of endogenous proteins (1627 residues in Penghu 2 and 2892 residues in Penghu 3) that have ≥7 razor+unique peptides in either Penghu 2 or Penghu 3. This corresponds to 15 proteins (AHSG, ALB, BGLAP, BGN, CHAD, COL1A1, COL1A2, COL2A1, COL5A1, COL5A2, COL9A1, F2, LUM, OMD, SERPINF1) as a whole. Corresponding sequences from Penghu 1 were included for comparison. Protein sequences of modern humans from the Yoruba population (NA19092, NA19117, and NA19121; 1000 Genomes Project) and archaic humans (Vindija 33.19, Chagyrskaya, Altai Neanderthal, and Denisova 3) were obtained from the Hominid Palaeoproteomic Reference Dataset[30,31]. Yoruba individuals were included as representative present-day African individuals to minimize the influence of archaic admixture in non-African populations[32]. Regarding sequences of great apes (*Pan troglodytes*, *Gorilla gorilla*, *Pongo abelii*) that were used in ref. 2, we used the same sequences from UniProt and Genbank. Other sequences of great apes were retrieved from the Hominid Dataset.

Alignment was conducted using MAFFT[33], and regions that might be affected by isoforms were excluded from AHSG, BGN, and COL5A1. Subsequent sequences were concatenated by each individual. Maximum-likelihood and Bayesian phylogenetic analyses were performed using IQ-TREE 2 version 2.3.6 (Model Finder, 1000 bootstrap replicates)[34] and BEAST 2 v2.7.7 (10,000,000 MCMC generations, with 10% burn-in with default parameters)[35], respectively (**Supplementary Fig. 6_1**; **Fig. 3**). The same tree topology was obtained for three different hominin taxa from the analysis performed with stricter sequencing criteria that require at least two peptides with spectral support to assign the amino acid residue at the position (**Supplementary Fig. 6_2**).

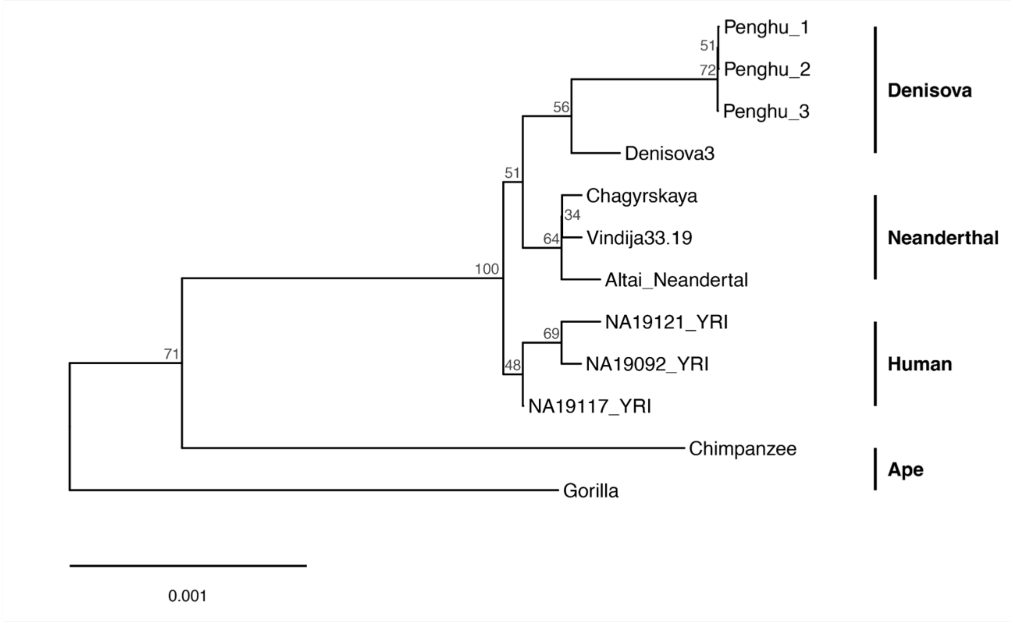


**Supplementary Figure 6_1. Maximum-likelihood phylogenetic tree of hominins and great apes inferred using IQ-TREE 2.** Node labels indicate bootstrap support values (0–100). The orangutan sample was excluded from the displayed tree.

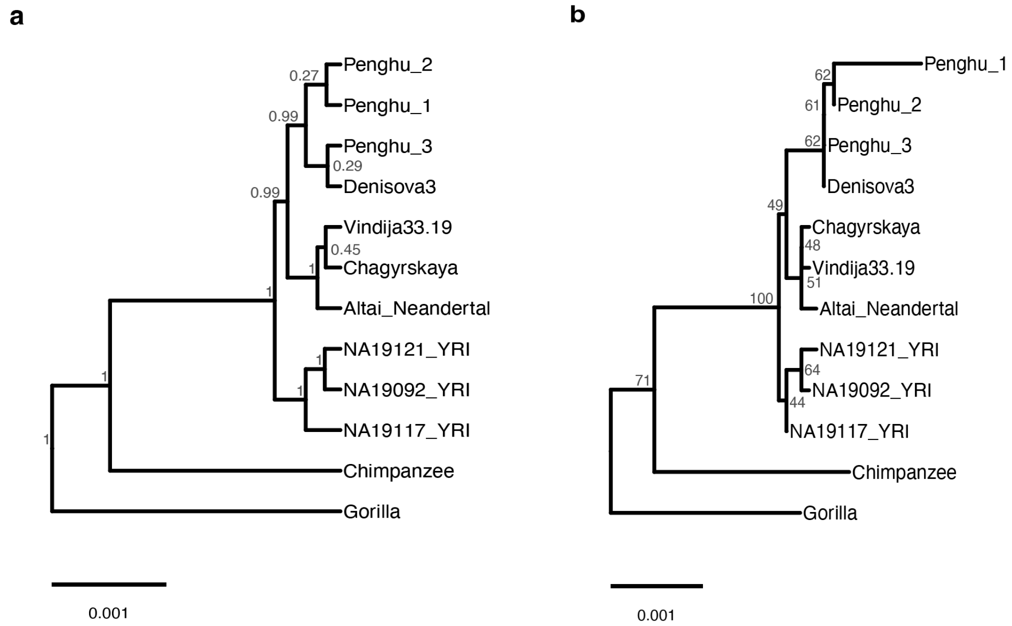


**Supplementary Figure 6_2. Bayesian (a) and maximum-likelihood (b) phylogenetic trees of hominins and great apes inferred using BEAST 2 and IQ-TREE 2, respectively.** Only amino acid residues supported by at least two peptides in Penghu 2 and Penghu 3 were included in the analyses. Node labels indicate posterior probabilities in the Bayesian tree and bootstrap support values in the maximum-likelihood tree. The orangutan sample was excluded from the displayed trees.

### Supplementary Note 7:
### Materials and methods of proteomic analysis

#### Specimen and sample collection

The Penghu 2 and Penghu 3 specimens were used with permission from NMNS. Following the best practices of palaeoproteomics[36], sampling was conducted inside clean laboratories at the Graduate University of Advanced Studies (SOKENDAI) and the University of Tokyo dedicated to the analysis of ancient biomolecules.

The broken distal ends of the specimens of Penghu 2 and Penghu 3 were removed for taking samples for dating and ancient DNA analyses, and the exposed inner parts of the compact bone were sampled for proteomic analyses. A total of 37.9 mg and 22.4 mg of bone powder samples were drilled from the inner part of Penghu 2 and Penghu 3, respectively, and collected on aluminum foil, then transferred into protein LoBind tubes after weighing. A hand-held dental drill with a carbide burr was used. First, ~5 mg of powder samples were sampled for trypsin digestion to evaluate protein preservation. After confirming the good preservation of ancient proteins, additional bone powders were sampled for protein extraction with the digestion-free method[14,37] and multiple proteases[6].

#### Extraction of bone proteins

The sampled bone powder was divided into four fractions and treated with digestion-free, non-filter trypsin digestion (two batches), and non-filter Glu-C digestion methods (**Supplementary Table 7_1**). The detailed procedures of each extraction method are described below. All fractions were treated along with experimental blanks that did not contain any sample.

*Digestion-free method*

Based on the previously developed method[14,37], approximately 5 mg bone powder aliquots were decalcified with 500 µL of 10% TFA at 4°C overnight. The solution was centrifuged to pellet tiny particles, and the supernatant was processed by C18 StageTips.

*Non-filter trypsin digestion method*

Based on the previously developed method[2,38], proteases were directly added to the decalcified supernatant without filtration and buffer exchange. Bone powder samples of 4.7–10.2 mg were decalcified with 500 µL of 0.5 M EDTA for 1–2 days under rotation. The samples were centrifuged (10,000 ×g, 5 min) and the supernatant was collected. The pellet fraction was washed 3 times with 100 µL of 50 mM Tris solution each time, and the solution was merged with the supernatant. TCEP and CAA were added to the final concentrations of 10 mM and 20 mM, respectively, to the supernatant fraction. For the pellet fraction, 250 µL Gu buffer (2M GuHCl, 10 mM TCEP, 20 mM CAA, and 100 mM Tris) was added. Supernatant and pellet fractions were incubated at 80°C for 1.5–2.5 hours. After the incubation, 0.2 µg of Lys-C (the approximate weight ratio of >1:128 relative to the total protein) was added and incubated at 37°C for 2 hours. The pellet fraction was diluted with the dilution solution (25 mM Tris and 10% ACN) to reduce the concentration of guanidinium chloride to less than 0.6 M. Then, 0.4 µg of trypsin (the approximate weight ratio of >1:64 relative to the total protein) was added and incubated at 37°C overnight for both fractions. After digestion, only the pellet fraction was acidified to pH <2.0 using 10% TFA to terminate the digestion. The supernatant fraction was not acidified to prevent the precipitation of EDTA. These digested solutions were then processed using C18 StageTips.

*Non-filter Glu-C digestion method*

The procedures were mostly the same as those in the non-filter trypsin digestion method. The non-filter Glu-C digestion method differs from the non-filter trypsin digestion method in the following steps. The washed pellet fraction was resuspended in 300 µL of 50 mM triethyl ammonium bicarbonate (TEAB) buffer containing 10 mM TCEP and 20 mM CAA, and incubated at 60°C for 2 hours. After incubation, 0.4 µg Glu-C (the approximate weight ratio of >1:64 relative to the total protein) was added to both fractions and incubated at 37°C overnight. After digestion, the pellet fraction was acidified by 10% TFA to pH <2 to inactivate the protease.

**Peptide purification by StageTips**

Decalcified or digested samples were purified and desalted using in-house-made StageTips with two stacked C18 membranes (CDS Analytical, USA)[9]. The C18 membranes were washed with 150 µL methanol and equilibrated with 150 µL of 80% ACN containing 0.1% TFA and then 150 µL of 0.1% TFA. The sample solution was incrementally added and passed through the C18 membranes. After passing the sample, the C18 membranes were washed twice with 150 µL of 0.1% TFA. Purified peptides were eluted from the StageTips using 50 µL of 40% ACN with 0.1% TFA into a new protein LoBind tube. Samples were placed in a vacuum centrifuge at 45°C until <3 µL of solution was left.

**LC-MS/MS measurement**

LC-MS/MS analysis was performed under similar conditions described in ref. 39. The dried peptides were resuspended in 2% acetonitrile and 0.1% trifluoroacetic acid in water. Peptides were separated using an Ultimate 3000 RSLCnano system (Thermo Fisher Scientific) with a reverse-phase Zaplous alpha Pep-C18 column (3 µm, 120 Å, 0.1 × 150 mm, AMR). The gradient was composed of solvent A (0.1% formic acid in water) and solvent B (100% acetonitrile). At the start of the analysis, the mobile phase was 5% solvent B at a flow rate of 500 nL per minute. The concentration of solvent B was linearly increased to 45% over 100 min. The column temperature was set at 35 °C. For peptide ionization, an electrospray voltage of 1.7 kV was applied via a nano-electrospray ion source (Dream Spray, AMR). The Orbitrap Fusion Tribrid mass spectrometer (Thermo Fisher Scientific) was operated in the positive ion mode, and the ion transfer tube temperature was set at 250 °C. All MS spectra were obtained using an Orbitrap mass analyzer (*m/z* range 350–1800; resolution 120,000 full width at half maximum) with EASY-IC internal mass calibration. MS/MS spectra resulting from collision-induced dissociation (CID) fragmentation were obtained in an Orbitrap mass analyzer (resolution 60,000 full width at half maximum , *m/z* range was auto).

**Database search for protein identification**

By conducting the preliminary searches using pFind and MaxQuant against the human proteome database, we aimed to list up the proteins and to identify the potential de novo substitutions to be included in the database for the final search.

*Preliminary search with pFind*

pFind version 3[40] was used in this study. Fractions with protease digestion were searched with the parameter "Enzyme" set to the respective proteases and "Full-Specific" protease specificity. All fractions, including digested and undigested bone, were also analyzed by specific and unspecific searches where the parameter "Enzyme" was set to "No_Enzyme". The error tolerance was set to 10 ppm for the precursor and to 0.07 Da for the fragment ion, respectively. The variable modifications were oxidation (M), deamidation (NQ), and hydroxylation (P). Carbamidomethylation (C) was set as a fixed modification for enzyme-specific search and as a variable modification for unspecific search. Peptide FDR was set to 1%. The parameter "Peptide Mass" was set to 600–5000, and "Peptide Length" was set to 6–30. Protein FDR was set to 100%. The database was the reference human proteome (UP000005640), downloaded on 2026-03-23.

*Preliminary search with MaxQuant*

MaxQuant version 2.6.3.0[10] was used in this study. The parameter "Enzyme" was set to the respective protease digestion conditions. Two types of searches with "Specific" and "Unspecific" protease specificity were conducted, and digestion-free fractions were only included in the unspecific search. The variable modifications were oxidation (M), deamidation (NQ), pyro-Glu (EQ), and hydroxylation (P). Carbamidomethylation (C) was set as a fixed modification for digested fractions. Error tolerances were kept at the default for Orbitrap mass spectrometers. Peptide and protein FDRs were set to 1%. Protein sequences of common laboratory contaminants provided by MaxQuant were included in the search database. The database was the reference human proteome (UP000005640), downloaded on 2026-03-23, for specific search and the subset of 32 proteins (AHSG, ALB, APOA1, BGLAP, BGN, C3, CHAD, CLEC3B, COL10A1, COL11A1, COL11A2, COL12A1, COL1A1, COL1A2, COL22A1, COL2A1, COL3A1, COL5A1, COL5A2, COL5A3, COL9A1, DCN, F10, F2, LUM, MGP, OMD, POSTN, SERPINF1, SPARC, SPP1, THBS1), that are typically identified from ancient bone/dentine samples, for unspecific search.

**Protein identification**

The above-mentioned preliminary searches against the human proteome using pFind and MaxQuant identified 88 proteins with at least two razor-unique peptides, excluding possible modern contaminants, and were merged with an additional 8 typical bone proteins to generate a tailored database (Human_Bone). Known archaic variants corresponding to these 96 proteins were also extracted using PaleoProPhyler (56). Then, the known archaic variants of the 96 identified proteins and the sequences with 35 possible de novo substitutions were added to the Human_Bone database to create the HomininVars_Bone database (**Supplementary Table 7_2**). All .raw files generated from Penghu 2 and 3 bone were searched against this

HomininVars_Bone database with "Specific", "Semi-specific", or "Unspecific" protease specificity using MaxQuant software to obtain the final identification results.

**Protein sequence reconstruction**

Ancient protein sequences were reconstructed from the outputs of the final MaxQuant searches using the HomininVars_Bone database. Results from different fractions and searches with different protease specificity were combined to create a unified consensus sequence. First, tandem mass spectra of all peptides listed in the MaxQuant output msms.txt files were computationally validated with a newly developed R script[11], and amino acid residues not supported by any peak were converted into "X" (unknown amino acid) in the sequence of the peptide. Peptides with a score of <70 were not used for the sequence reconstruction. Second, validated sequences of all peptides were mapped onto the entire length of each subject protein to create a table showing the position, corresponding amino acid, and depth indicating how many peptides cover the site. Since dynamic exclusion was used to increase sequence coverage, not all extracted molecules were measured by mass spectrometry analysis. Therefore, 'depth' in this study was not calculated based on the number of molecules as in the case of typical next-generation sequencing of DNA, but instead by the number of peptides covering each amino acid site. Results of proteins sharing almost identical sequences (e.g., isoforms and archaic variants) were aligned using MAFFT software and combined into a single table. Heterozygous positions were identified at this stage. Then, a unified consensus sequence was created for each protein from the table, with unsupported or unidentified positions converted to "X" (**Supplementary Data 1 and 2**). These sequences were used for phylogenetic analyses. The "Chain" region registered in UniProt for the corresponding human protein was used to identify the regions of mature protein that do not contain the signal sequence or the propeptide. The R script reported in ref. 17 and developed in ref. 2 was modified and used for sequence reconstruction, and the modified version of R script is accessible via doi: 10.5281/zenodo.21203230[41].

<u>Phylogenetic analysis</u>

Proteins with more than ≥7 razor+unique peptides were used for phylogenetic analysis. After this filtering, 15 proteins were used for the analysis. To compare phylogenetic relationships, protein sequences from Denisova 3, Neanderthals (n = 3), *H. sapiens* (n = 3), and apes (Chimpanzee, Gorilla, and Orangutan) were used. Sequences from hominins were analyzed using PaleoProPhyler[31]. Protein sequences of apes were sourced from UniProt and Genbank (Supplementary Table 8_3). For each protein, alignment was conducted using MAFFT, and regions affected by isoforms were excluded. Subsequent sequences were concatenated by each individual, and constructed phylogenetic trees using maximum likelihood method (iqtree2, model:, 1000 bootstrap repeats) and Bayesian approach (BEAST, 10,000,000 MCMC generations, with 10% burnin). The scripts[42] used for phylogenetic analysis are accessible in Zenodo repository.

**Supplementary Table 7_1. Summary of the Penghu 2 and Penghu 3 samples used for proteomic analysis.** These specimens were processed with four experimental batches.

| Specimen | ID | Element | Part | Amount (mg) | Digestion | Batch |
|---|---|---|---|---|---|---|
| Penghu 2 | NMNS006392 F051716 | Femur | Small piece | 4.7 | Trypsin | 1 |
| | | | Main body | 10.2 | Trypsin | 2 |
| | | | Main body | 5.1 | Digestion-free | 3 |
| | | | Main body | 17.9 | Glu-C | 4 |
| Penghu 3 | NMNS006392 F051713 | Tibia | Small piece | 5.9 | Trypsin | 1 |
| | | | Main body | 10.2 | Trypsin | 2 |
| | | | Main body | 5.4 | Digestion-free | 3 |
| | | | Main body | 0.9 | Glu-C | 4 |

**Supplementary Table 7_2. Corresponding gene names of the proteins that were used to create the HomininVars_Bone database for Penghu 2 and Penghu 3.** Entries listed in "Identified" correspond to those identified in the first MaxQuant and pFind searches against the human proteome. Entries listed in "Skeletal" correspond to those typically identified from ancient bone/dentine samples but are not included in the "Identified" entries. Entries listed in "De novo" correspond to those that include variant candidates that were inferred from the results of the pFind analysis.

| Classification | Gene names |
|---|---|
| Identified | Bis(monoacylglycero)phosphate synthase CLN5 (Fragment) (PID: A0A1B0GU22), Cofactor required for Sp1 transcriptional activation subunit 6 (PID: A0A1W2PRB8), ABCA13, ACTN3, AHSG, ALB, ANO5, AVL9, BAHD1, BGLAP, BGN, C1QB, C1QC, C21orf58, C9, CD36, CD63, CD81, CD9, CHAD, CLEC11A, COL1A1, COL1A2, COL10A1, COL11A1, COL12A1, COL15A1, COL16A1, COL17A1, COL18A1, COL2A1, COL21A1, COL22A1, COL23A1, COL24A1, COL26A1, COL28A1, COL3A1, COL4A3, COL4A4, COL4A6, COL5A1, COL5A2, COL5A3, COL7A1, COL8A1, COL8A2, COL9A1, COL9A2, COL9A3, COLEC12, DCAF7, DES, DUSP28, EMID1, EPPK1, F10, F2, F9, FAN1, FCN2, GOLGB1, KCNT2, KIF26A, KNG1, KRT87P, LRRC9, LUM, LZTFL1, MYH14, NUCB2, OLFML3, OMD, PROC, SERPINF1, SLC8A3, SPARC, SRPX, STK11IP, STXBP5, THBS1, TMEM119, TNFSF12, TRIM46, TRRAP, VIM, VIPR2, VTN |
| Skeletal | APOA1, C3, CLEC3B, COL11A2, DCN, MGP, POSTN, SPP1 |
| De novo | BGN, COL1A1, COL1A2, COL2A1, COL4A3, COL4A4, COL5A2 |

**Supplementary Data 1. Reconstructed protein sequences of Penghu 2.** (Separated data file)

**Supplementary Data 2. Reconstructed protein sequences of Penghu 3.** (Separated data file)

**Supplementary Note 8:**
**Morphological description**

Comparative fossil sample is listed in **Supplementary Table 8**.

**Penghu 2**

The specimen is a proximal half of the right femur that lacks much of the neck and greater trochanter (**Extended Data Fig. 1**, **Extended Data Table 2**). No postmortem distortion is evident and the surface preservation is excellent. The posterior base of the missing trochanteric fossa is marginally preserved at the proximal break. The distal break occurs at some distance below the midshaft level.

Viewed anteriorly or posteriorly, the medial margin of the basal neck and the lateral margin of the greater trochanter flare nearly equally, assuming a large funnel-like shape on top of the proximal shaft. The preserved root of the neck has an anteroposterior width of ~30 mm. It orients not transversely but rather superiorly. Viewed posteriorly, the slightly concave basal neck surface is diagonally extensive. Lateral to it, the preserved inferior two-thirds of the intertrochanteric crest is strong and also extensive. These and the large lesser trochanter (~27.7 mm long) suggest a great size of the missing proximal articular end of this specimen (**Extended Data Fig. 4b**).

The relatively straight shaft is robust throughout. Its mediolateral width continues to decrease or remains stable toward the distal break. The proximal shaft is not platymeric (**Fig. 4**). Viewed anteriorly, its lateral border expands slightly laterally (lateral buttress[43]) with a weak, blunt ridge formed lateral to the hypotrochanteric fossa (gluteal buttress[44]). The posterior surface of the proximal shaft is marked by moderately developed gluteal tuberosity, hypotrochanteric fossa (a 17 mm-long, longitudinal shallow depression), pectineal line and other muscle markings. The development of third trochanter is minimal, if any. The anterior, medial and lateral surfaces of the midshaft are convex and rounded overall. Its posterior face is marked by a strongly developed pilaster with the angular posterior edge. It emerges from ~45 mm distal to the inferior margin of the lesser trochanter, and gradually increases the degree of posterior extension toward the distal break of the shaft.

**Penghu 3**

This specimen is a right tibia that lacks the distalmost segment, with the maximum preserved length of 340 mm (**Extended Data Fig. 1**, **Extended Data Table 2**). No postmortem distortion is evident and the surface preservation is excellent. We have cleaned much of the nodules that had originally covered the proximal articular surface and some parts of the shaft (**Extended Data Fig. 1**), to reliably extract the bone surface data. Remains of nodule obscure the morphology of the anterior and central parts of the tibial plateau. The proximal epiphysis is complete except for sporadic damage around the posterior edge. Distally, the fibular notch is entirely missing.

The specimen is very large in any dimensions including the transverse proximal articular breadth, horizontal dimensions of the shaft, and length (**Extended Data Fig. 5**). The inferior half of the superior fibular articular facet is preserved at the superolateral corner of the proximal epiphysis. It faces not inferiorly but laterally and posteriorly. In anterior view, the proximal metaphysis shows ‘medial buttressing’ (R24) to form only a gentle concavity along the medial margin. The diaphysis is relatively straight in both anterior and lateral views, and shows only a moderate degree of retroversion of the tibial plateau. The diaphysis exhibits a mild degree of mediolateral compression and some concavity of the lateral face (see below), but does not show marked angularity and concavity often seen in modern human hunter-gatherers. The anterior crest is sharp, but the anterior margin is thickened throughout. The medial surface is gently convex. Approximately 50 mm distal to the tibial tuberosity on this surface is a bulging healed trauma of 40 mm long and 16 mm wide. The medial surface is undulated in transverse section. Posterior to the convolution along the anterior margin is ~18 cm-long, extensive longitudinal furrow that continues from top of the shaft downward beyond the midshaft level. Nearly parallel to the posterior border of this furrow is a weakly raised interosseus crest. The posterior surface is convex throughout and is marked by a strong tibial pilaster. It emerges from around the middle of the moderately raised soleal line, forming a distinct, triangular posterior projection with a blunt posterior margin, and gradually disappears onto the convex posterior surface of the distal diaphysis.

**Supplementary Table 8**. Comparative fossil sample.

| Specimen | Region | Bone | EFA | Data source |
|---|---|---|---|---|
| **early *Homo*/*Homo* sp. (2.1-1.7 million years ago)** | | | | |
| KNM-ER 5881 | Kenya | F | F | 45,46 |
| KNM-ER 1472 | Kenya | F | F | 45,46 |
| KNM-ER 1481 | Kenya | F, T | F | 45,46 |
| OH62 | Tanzania | F | F | 4 |
| KNM-ER 813 | Kenya | F | | 45 |
| D4167/D3901 | Georgia | F, T | | 45,47 |
| ***H. erectus*/*ergaster* (1.6-0.9 million years ago)** | | | | |
| KNM-ER 1808mn | Kenya | F | F | 45,46 |
| KNM-ER 737 | Kenya | F | F | 45,46 |
| KNM-ER 736 | Kenya | F | F | 45,46 |
| KNM-ER 803 | Kenya | F, T | F | 45,46 |
| BOU-VP-2/15 | Ethiopia | T | F | 48 |
| OH 28 | Tanzania | F | F | 45,46 |
| ***H. erectus* (1.1-0.1 million years ago)** | | | | |
| Trinil II | Indonesia | F | F | 49 |
| Trinil III | Indonesia | F | | 49 |
| Trinil IV | Indonesia | F | F | 49 |
| Trinil V | Indonesia | F | F | 49 |
| Kresna 11 | Indonesia | F | F | 50 |
| Ngandong 13 | Indonesia | T | | 45 |
| Ngandong 14 | Indonesia | T | | 45 |
| Zhoukoudian Femur I | China | F | F | 45,51 |
| Zhoukoudian Femur II | China | F | F | 45,51 |
| Zhoukoudian Femur IV/V | China | F | F | 45,51 |
| Zhoukoudian Femur VI | China | F | | 45 |
| PA65 | China | T | | 45 |
| **Middle Pleistocene (MP) archaic *Homo* (late archaic *Homo*)** | | | | |
| **Eastern Asia** | | | | |
| Hualongdong 11 | China | F | F | 52 |
| Hualongdong 15 | China | F | | 52 |
| Hualongdong 16 | China | F | | 52 |
| **West Asia** | | | | |
| Gesher-B.-Y. 1 | Israel | F | | 45 |

| | | | | |
|---|---|---|---|---|
| Gesher-B.-Y. 2 | Israel | F | | 45 |
| Tabun E1 | Israel | F | | 45 |
| **Africa** | | | | |
| Broken Hill E689 (femur) | Zambia | F | | 45 |
| Broken Hill E690 (femur) | Zambia | F | | 45 |
| Broken Hill E691 (tibia) | Zambia | T | | 45 |
| Broken Hill E709 (femur) | Zambia | F | | 45 |
| Broken Hill E793 (femur) | Zambia | F | | 45 |
| Hoedjiespunt 1 | South Africa | T | | 45 |
| Berg Aukas 1 (age unknown) | Namibia | F | F | 53 |
| KNM-ER 999 (age/group unknown) | Kenya | F | F | 54 |
| Aïn Maarouf 1 | Morocco | F | | 45 |
| **Europe** | | | | |
| Arago 48 | France | T | | 45 |
| Arago 53 | France | T | | 45 |
| Arago 57 | France | T | | 45 |
| Arago 141 | France | F | F | 45, T. Chevalier |
| Arago 120 | France | T | T | 45, T. Chevalier |
| Boxgrove 1 | England | T | T | 45 |
| Castel del Guido 1 | Italy | F | | 45 |
| La Chaise-BD 5 | France | F | | 45 |
| Ehringsdorf 5 | Germany | F | F | 45, T. Chevalier |
| Mammolo 1 | Italy | F | | 45 |
| Sedia-del-Diavolo 1 | Italy | F | | 45 |
| Atapuerca SH AT-616 | Spain | F | | 43,55 |
| Atapuerca SH AT-1020 | Spain | F | | 43,55 |
| Atapuerca SH F-IX | Spain | F | F | 43,55 |
| Atapuerca SH F-X | Spain | F | F | 43,55 |
| Atapuerca SH F-XI | Spain | F | | 43 |
| Atapuerca SH F-XII | Spain | F | | 43 |
| Atapuerca SH F-XIII | Spain | F | F | 43,55 |
| Atapuerca SH F-XIV | Spain | F | F | 43,55 |
| Atapuerca SH F-XVI | Spain | F | F | 43,55 |
| Atapuerca SH Tib I | Spain | T | T | 55,56 |
| Atapuerca SH Tib III | Spain | T | T | 55,56 |
| Atapuerca SH Tib IV | Spain | T | T | 55,56 |
| Atapuerca SH Tib VI | Spain | T | T | 55,56 |

| | | | | |
|---|---|---|---|---|
| Atapuerca SH Tib XI | Spain | T | T | 55,56 |
| Atapuerca SH Tib XII | Spain | T | T | 55,56 |
| Atapuerca SH AT-848 | Spain | T | T | 55,56 |
| **Neanderthal** | | | | |
| Krapina 213 | Croatia | F | | 45 |
| Krapina 214 | Croatia | F | | 45 |
| Krapina 257.15 | Croatia | T | | 45 |
| Krapina 257.20 | Croatia | T | | 45 |
| Krapina 257.32 | Croatia | F | | 45 |
| Krapina 257.33 | Croatia | F | | 45 |
| Tabun 1 | Israel | F, T | F, T | 45,57,58 |
| Tabun 3 | Israel | F | F | 45,57 |
| Amud 1 | Israel | F, T | F, T | 45,57,58 |
| Shanidar 1 | Iraq | F, T | | 45 |
| Shanidar 2 | Iraq | T | T | 45,58 |
| Shanidar 4 | Iraq | F | | 45 |
| Shanidar 5 | Iraq | F | F | 45,57 |
| Shanidar 6 | Iraq | F, T | | 45 |
| La Chapelle-aux-Saints 1 | France | F, T | T | 45, T. Chevalier |
| Feldhofer 1 | Germany | F | F | 45,59 |
| Ferrassie 1 | France | F, T | F, T | 45, T. Chevalier |
| Ferrassie 2 | France | F, T | | 45 |
| Fond-de-Forêt 1 | Belgium | F | F | 45,60 |
| Hortus 34 | France | F | | 45 |
| Kiik-Koba 1 | Crimea | T | | 45 |
| Oliveira 4 | Italy | T | | 45 |
| Palomas 52 | Spain | F | | 45 |
| Palomas 92 | Spain | F | | 45 |
| Palomas 96 | Spain | F, T | | 45 |
| Pofi 1 | Italy | T | | 45 |
| Quina 5 | France | F | | 45 |
| Quina 38 | France | F | | 45 |
| Rochers-de-V. 1 | France | F | | 45 |
| Saint-Césaire 1 | France | F, T | | 45 |
| Santa Croce 1 | Italy | F | | 45 |
| Spy 2 | Belgium | F, T | F | 45,60 |
| Stadelhöhle 1 | Germany | F | | 45 |

| | | | | |
|---|---|---|---|---|
| Zafarraya 1 | Spain | F | | 45 |
| **MIS 6/5/4 *H. sapiens*** | | | | |
| Omo-Kibish 1 | E Africa | T | | 45 |
| Omo-Kibish 158-1a | E Africa | T | | 45 |
| Qafzeh 3 | W Asia | F, T | | 45 |
| Qafzeh 6 | W Asia | F | | 45 |
| Qafzeh 8 | W Asia | F, T | F, T | 45,57,58 |
| Qafzeh 9 | W Asia | F, T | F | 45,57 |
| Skhul 3 | W Asia | F, T | | 45 |
| Skhul 4 | W Asia | F, T | T | 45,58 |
| Skhul 5 | W Asia | F, T | F, T | 45,57,58 |
| Skhul 6 | W Asia | F, T | T | 45,58 |
| Skhul 7 | W Asia | F | F | 45,57 |
| Skhul 9 | W Asia | F | | 45 |
| Skhul '4' | W Asia | T | | 45 |
| Skhul '7' | W Asia | F | | 45 |
| Skhul '9' | W Asia | F | | 45 |
| **MIS 3/2 *H. sapiens*** | | | | |
| Dolní Věstonice 3 | Czeck | F, T | F, T | 45,61 |
| Dolní Věstonice 13 (adolecent) | Czeck | F, T | F, T | 45,61 |
| Dolní Věstonice 14 | Czeck | F, T | F, T | 45,61 |
| Dolní Věstonice 16 | Czeck | F, T | F, T | 45,61 |
| Dolní Věstonice 35 | Czeck | F | F | 45,61 |
| Dolní Věstonice 40 | Czeck | F | | 45 |
| Pavlov 1 | Czeck | F | F | 45,61 |
| Předmostí 3 | Czeck | F, T | | 45 |
| Předmostí 4 | Czeck | F, T | | 45 |
| Předmostí 9 | Czeck | F, T | | 45 |
| Předmostí 10 | Czeck | F, T | | 45 |
| Předmostí 14 | Czeck | F, T | | 45 |
| Mladeč 27 | Czeck | F | | 45 |
| Mladeč 28 | Czeck | F | | 45 |
| Sunghir 1 | Russia | F, T | F, T | 45,62 |
| Sunghir 4 | Russia | F | F | 45,62 |
| Willendorf 1 | Austria | F | | 45 |
| Arene Candide 1 | Italy | F, T | | 45 |
| Barma Grande 1 | Italy | F, T | | 45 |

| | | | | |
|---|---|---|---|---|
| Barma Grande 2 | Italy | F, T | | 45 |
| Barma Grande 5 | Italy | T | | 45 |
| Barma Grande 6 | Italy | F, T | | 45 |
| Caviglione 1 | Italy | F, T | T | 45,63 |
| Grotte des Enfants 4 | Italy | F, T | | 45 |
| Grotte des Enfants 5 | Italy | F, T | | 45 |
| Paglicci 25 | Italy | F, T | | 45 |
| Paviland 1 | Wales | F, T | F | 45,59 |
| Veneri 1 | Italy | F, T | | 45 |
| Veneri 2 | Italy | F, T | | 45 |
| Cro-Magnon Alpha | France | F, T | F, T | 45,64 |
| Cro-Magnon Beta | France | F | F | 45,64 |
| Cro-Magnon Gamma | France | F, T | F, T | 45,64 |
| Rochette 2 | France | F | | 45 |
| Bruniquel 24 | France | T | | 45 |
| Bichon 1 | Swiss Alps | T | | 45 |
| Cap Blanc 1 | France | T | | 45 |
| Ohalo 2 | Israel | F, T | | 45 |
| Nahal ‘En-Gev 1 | Israel | F, T | | 45 |
| Ust’-Ishim | West Siberia | F | F | 24,45 |
| Tianyuan 1 | China | F, T | F, T | 45,65 |
| Zhoukoudian-UC 67 (Femur II) | China | F, T | F | 45,51 |
| Zhoukoudian-UC 68 (Femur I) | China | F, T | F | 45,51 |
| Liujiang 1 | China | F | | 45 |
| Maomaodong GM7506 | China | F | F | 66 |
| Maomaodong GM7507 | China | F | F | 66 |
| Maomaodong GM7508 | China | F | F | 66 |
| Niah | Malaysia | F, T | F | 67 |
| Trinil 3 (Femur I) | Indonesia | F | F | 49 |
| Trinil 9 | Indonesia | F | F | 68 |
